# Multiscale analysis and functional validation of the cellular and genetic determinants of skeletal disease

**DOI:** 10.1101/2024.12.16.628792

**Authors:** Ryan C. Chai, Mischa Lundberg, Bernard Freudenthal, James T. Smith, Andrew P. Boughton, Yuandan Zhang, Kaitlyn A. Flynn, Monika Frysz, Alexander P. Corr, Weng Hua Khoo, Davide Komla-Ebri, Michael R.G. Dack, Siobhan E. Guilfoyle, John G. Logan, Natalie C. Butterfield, Victoria D. Leitch, Andrea S. Pollard, Riikka E. Mäkitie, Nathaniel Bradford, Lorenzo Ramos-Mucci, Amaia Vilas-Zornoza, Yunshun Chen, Raymond K.H. Yip, Jeremy Er, Siew Zhuan Tan, Michelle M. McDonald, Scott E. Youlten, C. Marcelo Sergio, Ariel Castro-Martinez, Shelley G. Young, Elena Skorokhodova, David M. Evans, Joseph E. Powell, Christiaan A. de Leeuw, Adam D. Ewing, John A. Eisman, Robert D. Blank, Tri Giang Phan, International Federation of Musculoskeletal Research Societies (IFMRS) Big Data Working Group, Anne K. Lagendijk, Edwin D. Hawkins, Horng Lii Oh, Rebecca McIntyre, Edith M. Hessel, Jake P. Taylor-King, Paul A. Baldock, Emma L. Duncan, Graham R. Williams, J. H. Duncan Bassett, Peter I. Croucher, John P. Kemp

**Author notes:** These authors contributed equally: Ryan C. Chai, Mischa Lundberg, Bernard Freudenthal & James T. Smith. These authors jointly supervised this work: Graham R. Williams, J. H. Duncan Bassett, Peter I. Croucher & John P. Kemp.

## Abstract

Musculoskeletal diseases are a major health burden. Development of bone-active therapies has been hindered by limited understanding of the cells and genes that regulate the skeleton. We exploited the value of cross-species analysis and developed single-cell methodologies in skeletal tissues to define the critical endosteal compartment that regulates bone turnover. Thirty-four distinct cell types were identified, and disease-relevant cells prioritised by enrichment for rare skeletal disorder genes and bone mineral density-associated genes in an extended UK Biobank GWAS. Functional validation was undertaken in over one thousand genetically modified mouse models. Endothelial and vascular smooth muscle cells were identified as novel skeletal disease-relevant cells alongside osteoblast, chondrocyte and osteoclast cell lineages. Hundreds of cell-specific genes with unappreciated roles in skeletal pathophysiology were identified. This comprehensive cellular and molecular framework underpins skeletal physiology and disease, and will help prioritise new therapeutic targets to accelerate development of novel therapies to treat musculoskeletal disease.

## Introduction

Musculoskeletal diseases such as osteoporosis, affect half of individuals over 50 years of age and are a major health burden^1,2^. Whilst existing treatments prevent further skeletal deterioration, they rarely restore skeletal integrity, creating an urgent need for new therapies^3^, which has been hindered by limited understanding of the cellular and molecular mechanisms that maintain bone mass and strength.

Knowledge from human genetics can accelerate discovery programs^4^. Gene-mapping of rare disorders and common disease traits have uncovered new mechanisms of skeletal regulation^5^. Genome-wide association studies (GWAS) recently identified over five hundred loci associated with estimated bone mineral density (eBMD), a measure of skeletal integrity^6–9^. Many of these loci contain effector genes, defined as genes with a functional role in regulating the skeleton, yet the genes underlying the associations at most loci remain unknown. Methods that integrate GWAS with expression quantitative trait locus analysis (eQTL) of bulk tissues can help identify effector genes^10^, but is constrained by limited knowledge of gene function in specific cell types^11–13^. Application of eQTL methods to single-cell transcriptomic datasets has the potential to address this constraint^14^. However, it is difficult to obtain bone cells from healthy individuals without underlying disease^15^. Furthermore, a key site of bone turnover is the endosteal compartment, which is located at the interface between bone and bone marrow. Isolation of cells from this compartment is challenging and data from bone is lacking or poorly represented in Genotype Tissue Expression (GTEx) project and the Human Cell Atlas^16,17^. Given that genes in humans and mice are highly conserved, along with recent success in analysing GWAS and transcriptomic data from mouse tissue^18–21^, we reasoned that isolating cells from the endosteal compartment of mice could address these challenges.

We hypothesised that integrating gene expression data from a map of cells isolated from the endosteal compartment of murine bone with human gene-mapping studies of rare skeletal disorders and common skeletal disease traits would identify genes with important functional roles in the skeleton that contribute to the pathogenesis of human skeletal diseases and might be amenable to therapeutic intervention. We generated a single-cell RNA-sequencing (scRNA-seq) map of cells isolated from the endosteal compartment of healthy mice. We integrated this dataset with analyses of: (i) the current nosology of rare monogenic skeletal disorders^22^, (ii) a new and now largest GWAS of eBMD, (iii) skeletal phenotyping data from the mouse genome informatic (MGI) database, (iv) an in-depth functional analysis of the skeleton in over one thousand mouse lines with unselected single gene deletions and (v) validated outcomes in a new scRNA-seq dataset from adult human bone.

Our integrated, multiscale and cross-species approach defined the cellular landscape of the endosteal compartment and identified effector genes in osteoblasts, chondrocytes, osteoclasts, vascular smooth muscle cells and endothelial cells that cause rare and common skeletal disorders. To make our comprehensive datasets accessible to the scientific community we developed an open access platform that is available at www.musculoskeletal-genomics.org.

## Results

### Defining the repertoire of cell types in the endosteal compartment

To identify genes that regulate the skeleton and define the cells in which they function, we developed a multiscale framework (Fig. 1a). This began by isolating cells from the endosteal compartments of the metaphysis and diaphysis, and from bone marrow from femurs of adult male C57BL/6J mice (Fig. 1b) for scRNA-seq analysis. Gene programs (the set of genes that are differentially upregulated in one cell cluster relative to all others) of individual cell types found within the endosteal bone compartment were then defined (Supplementary Table 1. Glossary).

**Fig. 1.**
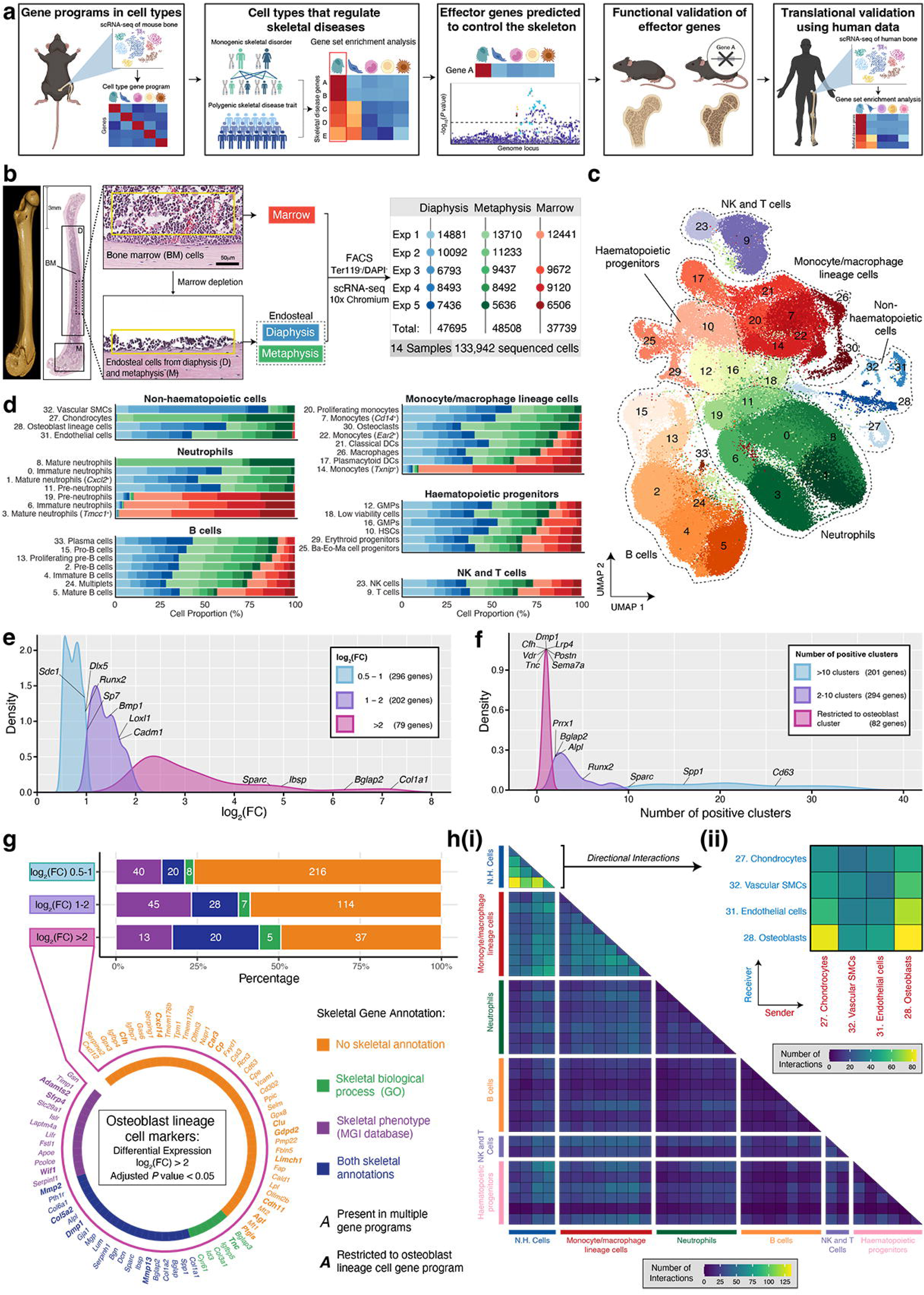
Mapping the cell types in the endosteal compartment. (**a**) Outline of the workflow used to identify and functionally validate effector genes at loci associated with eBMD. Created with BioRender (www.biorender.com) with permission to publish. (**b**) Anatomical regions of the mouse femur from which endosteal bone cells and bone marrow cells were cells isolated and sequenced from individual experiments. Numbers in table indicate total number of cells present post-filtering. (**c**) UMAP plot derived from scRNA-seq data generated from (**b**) showing 34 cell clusters and 6 broad cell categories. (**d**) Distribution of cell types in the 6 cell categories from (**c**) in diaphysis (blue shades), metaphysis (green shades) and marrow (red shades). Each shade represents an individual experiment as detailed in (**b**). (**e**) Density plot of log_2_ fold change [log_2_(FC)] of genes that define the osteoblast cluster. (**f**) Density plot of the number of other cell clusters in which genes from the osteoblast cluster gene program were detected. (**g**) Bar plots showing proportion of genes in the osteoblast lineage cell gene program that are annotated with roles in the skeleton, stratified by log_2_(FC) 0.5-1 (top), log_2_(FC) 1-2 (middle) and log_2_(FC) >2 (bottom) change in expression. Genes annotated with a skeletal process in the gene ontology (GO) database (green), the mouse genome informatics (MGI) database (purple), in both databases (blue) or are not annotated in either (orange) are shown. Numbers denote the number of genes in each category. Circos plot showing individual genes with a log_2_(FC) >2 in expression, with those found only in the osteoblast lineage cluster gene program indicated in bold. (**h**) Heatmaps displaying the number of predicted intercellular interactions between each cell cluster, as determined by CellPhoneDB. Panel (i) shows all interactions between every pair of clusters within the dataset, whilst panel (ii) shows interactions between non-haematopoietic clusters separated according to the directionality of the interaction (see Methods).

A total of 133,942 cells were included in subsequent analyses. These cells formed 34 cell clusters (Fig. 1c) comprising six categories: non-haematopoietic cells (4 clusters), monocytes and macrophages (8 clusters), neutrophils (7 clusters), haematopoietic progenitors (6 clusters), B cells (7 clusters), and NK and T cells (2 clusters) (Supplementary Table 2). Cell annotation identified clusters that corresponded to distinct cell types (Supplementary Note 1; Supplementary Table 2). Non-haematopoietic cells, which included chondrocytes, osteoblast lineage cells, endothelial cells, and vascular smooth muscle cells (VSMCs) originated largely from the endosteal compartments of the diaphysis and metaphysis (Fig. 1d). Chondrocytes were derived exclusively from the metaphysis, consistent with their known location in the growth plate. Cells of the monocyte/macrophage lineage, including dendritic cells and macrophages, which may include osteal macrophages^23^, were found in both endosteal and bone marrow compartments. *Txnip*^+^ monocytes were found predominantly in the bone marrow compartment; whilst proliferating monocytes and *Cd14*^+^ monocytes were more abundant in the endosteal compartments (Fig. 1d), an enrichment confirmed by flow cytometry (Supplementary Fig. 1). This lineage also included cells denoted as osteoclasts, which were found in both endosteal compartments. While large multinucleated cells are normally excluded from scRNA-seq, these cells express genes typical of osteoclasts and may represent precursors, mononuclear osteoclasts, or osteomorphs^24^. Of note, cells of this lineage can fragment during disaggregation and remnants of these cells can be associated with unrelated cells^25,26^. Neutrophils at different stages of maturation, haematopoietic progenitor cells (haematopoietic stem cells, basophil-eosinophil-mast cell progenitors and granulomyeloid progenitors), B cells, T and NK cells were also identified, showing varying distribution among compartments.

Osteoblast lineage cells mediate bone formation and regulate skeletal integrity. Therefore, we determined the gene programs that define osteoblasts and other cell types enriched in the endosteal compartment. The osteoblast gene program consisted of 577 genes (Supplementary Table 2). Seventy-nine genes (13.7%), including *Col1a1*, *Bglap2*, and *Ibsp*, were highly upregulated (log2 FC>2) (Fig. 1e). While most genes (495 genes, 85.8%) were expressed in at least one other cell cluster, 82 genes (14.2%) were considered to have restricted expression (see Supplementary Table 1 and Methods for criteria), including *Dmp1*, *Vdr*, and *Postn* (Fig. 1f). Notably, 387 genes (67.1%) in the osteoblast program were neither annotated in the gene ontology (GO) database for skeletal terms nor had an abnormal skeletal phenotype in the MGI database of knockout mice (Fig. 1g).

The gene programs of other key endosteal cells were also defined. The chondrocyte program included 546 genes, with 71 highly expressed genes, including *Col2a1*, *Comp* and *Acan*, and 136 genes with restricted expression (Extended Data Fig. 1a,b; Supplementary Table 2). The vasculature is also important for bone development, regeneration and remodelling^27^. The endothelial cell program included 610 genes, with 78 highly expressed genes (Extended Data Fig. 1c,d). The VSMC program included 636 genes, with 112 highly expressed genes (Extended Data Fig. 1e,f). Importantly, 26.9% and 27.7% of highly expressed genes in endothelial cells and VSMCs, respectively, were annotated in the GO database with a skeletal biological process or had an abnormal skeletal phenotype in mice in the MGI database (Extended Data Fig. 1d,f). Finally, the osteoclast program included 687 genes, of which 19 genes (2.8%) were highly expressed and forty-nine (7.1%) were expressed only in osteoclasts (Extended Data Fig. 1g,h), including genes crucial for osteoclast differentiation and resorption such as *Tnfrsf11a*, *Dcstamp*, and *Atp6v0d2*.

Since these cells may regulate the skeleton through direct cell-cell communication, we performed interaction analysis. This revealed the greatest number of interactions among non-haematopoietic cells, including between osteoblast lineage cells and the vasculature (Fig. 1h; Supplementary Table 3). Interactions were mediated by ligand-receptor pairs, adhesion molecules, and extracellular matrix components (Extended Data Fig. 2a). Osteoblasts interact with other cell types via common (e.g. TGF-β) and distinct pathways (e.g. WNT with chondrocytes, NOTCH with endothelial cells, IGF-1R with VSMCs; Extended Data Fig. 2b).

Our approach identified the repertoire of cell types and gene programs that define cells in the endosteal bone compartment. Osteoclasts, osteoblast lineage cells, chondrocytes, endothelial cells, and VSMCs originated almost exclusively (>98%) from this compartment. Of the 1886 genes defining the gene programs of these five clusters, 1374 genes (72.9%) had not previously been implicated in skeletal regulation.

### Non-haematopoietic cell differentiation in the endosteal compartment

Since the cells in the endosteal compartments are at different stages of differentiation, we used high-resolution sub-clustering of the non-haematopoietic cells to define these stages. Sixteen clusters were identified (Extended Data Fig. 3a,b; Supplementary Table 2), comprising 6 clusters within the osteoblast lineage, 3 chondrocyte clusters, 4 endothelial cell clusters, 2 VSMC clusters, and a single neuronal cluster.

The osteoblast lineage formed a continuum: MSCs progressed through fibroblasts and osteoprogenitors to pre-osteoblasts, mature osteoblasts and early osteocytes (Extended Data Fig. 3c, Supplementary Table 2). Pre-osteoblast number was higher in the diaphysis relative to the metaphysis (Supplementary Fig. 2). Gene expression was progressively modulated throughout differentiation (Extended Data Fig. 3b,d). *Cxcl12*^+^*/Adipoq*^+^ MSC and *Apod*^+^/*Igfbp5*^+^ fibroblast programs were enriched for GO processes in cell proliferation and migration, respectively (Extended Data Fig. 3e). *Postn*^+^*/Col3a1*^+^ osteoprogenitors and *Car3*^+^*/Mmp13*^+^ pre-osteoblasts were associated with extracellular matrix organisation, *Bglap2*^+^/*Col1a1*^+^ mature osteoblasts with collagen fibril organisation, and *Mepe*^+^/*Phex*^+^ early osteocytes with ossification. This trajectory coincided with the sequential increase in expression of key transcription factors (TFs) and their target genes as predicted by SCENIC analysis, including *Cebpa* and *Foxc1* in MSCs, which are associated with adipogenic and osteogenic progenitors^28,29^, as well as *Runx2* and *Sp7* in committed cells^30,31^ (Extended Data Fig. 4a). Parallel single-nucleus ATAC-seq (snATAC-seq) analysis identified unique and overlapping TF-binding motifs in osteoblast lineage cells (Extended Data Fig. 4b-e; Supplementary Table 4). SnATAC-seq identified *Tfap2a*, *Ebf2* and *Zic1* with known roles in the skeleton^32–34^, as well as novel TFs such as *Ebf3*, *Maz* and *Ctcfl*. TFs identified by both methods included *Runx1* and *Creb3l1*, as well as *Klf15*, *Etv4* and *Glis2* which have unappreciated roles in bone (Extended Data Fig. 4e).

Critically, 56.3%−72.6% of genes within these osteoblast sub-cluster gene programs were not annotated with skeletal terms in the GO or MGI database, indicating that many genes involved in skeletal differentiation and regulation have yet to be studied (Extended Data Fig. 3f).

Chondrocytes also showed a trajectory of cell states, transitioning from *Ucma*^+^/*Serpina1b*^+^ resting chondrocytes to *Ppa1*^+^/*Scrg1*^+^ pre-hypertrophic chondrocytes and *Col10a1*^+^/*Ihh*^+^ hypertrophic chondrocytes (Extended Data Fig. 3a,b,g,h; Supplementary Fig. 2; Supplementary Table 2). Chondrocytes formed a distinct differentiation trajectory, separate from the osteoblast lineage and were restricted to the metaphysis (Extended Data Fig. 3i, Supplementary Fig. 2). The endothelial cell clusters included *Ubd*^+^/*Tfpi*^+^ sinusoidal cells, two clusters of arteriolar cells (*Cldn5*^+^/*Gkn3*^+^ and *Ly6c1*^+^/*Glul*^+^) and *Plvap*^+^/*Aplnr*^+^ type H cells, which have been implicated in skeletal development^35^ (Extended Data Fig. 3a,b; Supplementary Table 2). VSMC clusters included smooth muscle cells (SMCs) expressing myosin and actin (*Myl9*^+^/*Acta2*^+^) and pericytes expressing *H2-M9* and *Rgs5*. Arteriolar cells (*Cldn5*^+^/*Gkn3*^+^) and SMCs were enriched in the diaphysis, whereas type H ECs were enriched in the metaphysis (Supplementary Fig. 2), consistent with previous reports^35^.

This high-resolution analysis identified the stages of differentiation in non-haematopoietic cells in the endosteal compartment. Many of the genes identified in these cells are not currently known to regulate the skeleton.

### Non-haematopoietic cell and osteoclast gene programs are enriched with genes that cause rare human skeletal disorders

We next investigated whether cell type gene programs were enriched with genes that cause rare monogenic skeletal disorders when their function is altered by pathogenic mutations (causative genes, Supplementary Table 1). We used the International Skeletal Dysplasia Society (ISDS) nosology, a reference set of 533 protein-coding genes with pathogenic mutations causing 719 rare disorders^22^. Whilst multiple cell types expressed these causative genes (Fig. 2a), gene-set enrichment analyses showed that only some gene programs were highly enriched with causative genes, and these included osteoblast lineage cells, chondrocytes, osteoclasts, proliferating monocytes, and proliferating pre-B cells were highly enriched with monogenic skeletal disorder genes (Fig. 2a; Supplementary Table 5).

**Fig. 2.**
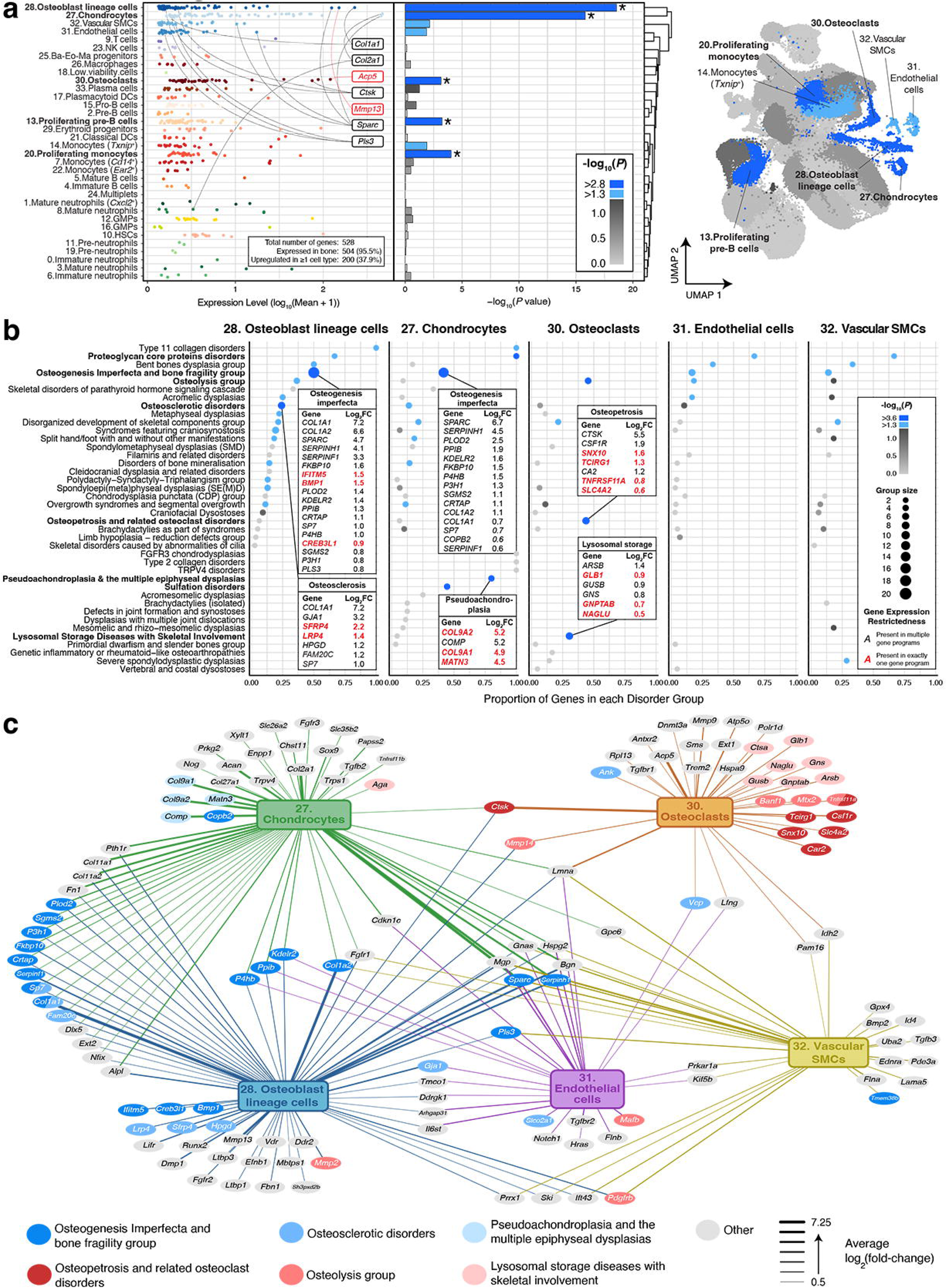
Gene programs of specific cell types are enriched with genes involved in human monogenic skeletal disorders. (**a**) Dotplot showing the mean expression level (log_10_ mean+1) of known monogenic skeletal disorder genes in individual gene programs. Exemplar genes have been annotated with gene symbols. Red text denotes a gene present in a single gene program and not present in any other gene program. Scale bar in bar plot and UMAP plot indicates the *P* value. Light blue bars in bar plot and light blue dots in UMAP correspond to observations that have nominal evidence of enrichment: *P* value of <0.05 [-log_10_ (*P* value) of >1.3]. Dark blue bars and asterisks in bar plot and dark blue dots in UMAP correspond to observations that have robust evidence of enrichment and meet the *Bonferroni* corrected significance threshold: *P* value of < 1.5 ×10^-3^ [-log_10_(*P* value) > 2.8]. (**b**) Bubble plot showing the number and proportion of monogenic skeletal disorder genes from each disorder group that are present in gene programs of selected cell types. 39 disorder groups are identified. Those enriched with disorder-causing genes in at least one of the five identified cell types are indicated in bold. Size of the circles represent the number of disorder group genes within the gene program of the corresponding cell type. Scale bar indicates the *P* value of enrichment, as determined by hypergeometric tests of over-representation. Light blue dots indicate nominal evidence of enrichment: *P* value of <0.05 [-log_10_(*P* value) of >1.3]. Dark blue dots denote robust evidence of enrichment with *Bonferroni*-corrected significance threshold of *P* value < 2.4 × 10^-4^ [-log_10_(*P* value) of >3.6]. Boxes identify the causative genes for the indicated disorder group present in a cell type gene program. Red text denotes a gene found only in that gene program and not shared with other gene programs. (**c**) Network plot showing the pattern and magnitude of gene expression of skeletal disorder genes among selected cell types. Genes are colour coded based on disorder group and the magnitude of differential expression [log_2_(FC)] is indicated by the thickness of the connecting lines.

In osteoblast lineage cells, 61 genes (out of 553 human orthologs) cause rare skeletal disorders, representing a four-fold enrichment (*P*=2.8×10^−19^, Supplementary Table 5). All differentiation stages except MSCs were enriched (Extended Data Fig. 5a). Osteoblast lineage cells were primarily enriched with genes causing osteogenesis imperfecta (OI) and other bone fragility disorders, and osteosclerotic disorders (Fig. 2b,c). Eighteen of 36 known OI/bone fragility genes were present in the osteoblast lineage program, with causative genes over-represented in pre-osteoblasts and mature osteoblasts (Extended Data Fig. 5b; Supplementary Table 5). Genes causing osteosclerotic and bone mineralisation disorders were enriched in pre-osteoblasts and early osteocytes.

Chondrocytes were enriched with causative genes for proteoglycan core protein disorders, sulfation disorders, pseudoachondroplasia, multiple epiphyseal dysplasias, and OI/bone fragility (Fig. 2b,c). Enrichment patterns varied among chondrocyte subtypes (Extended Data Fig. 5c,d). Osteoclasts were enriched with disease genes affecting bone mass, including those causing osteopetrosis and osteolysis/lysosomal storage disorders (Fig. 2b,c). Proliferating monocytes and pre-B cells were enriched with genes causing dwarfism, potentially reflecting expression of associated cell cycle genes^36^ (Supplementary Table 5). Endothelial cells and VSMCs showed only nominal evidence (*P*<0.05, not significant after Bonferroni-correction) of enrichment for certain disorders (e.g., proteoglycan core protein disorders and bent bone disease), possibly due to shared gene expression with osteoblast lineage cells and chondrocytes (Fig. 2b,c).

This analysis showed that gene programs defining key endosteal cells including osteoblasts, chondrocytes, osteoclasts were highly enriched with genes causing rare skeletal disorders. While many causative genes were cell type-restricted, others were expressed in multiple cell types.

### Non-haematopoietic cell gene programs are enriched with genes associated with eBMD

We then investigated whether gene programs that define cells in the endosteal compartment are also enriched with eBMD-associated genes. We conducted: (1) GWAS to identify variants associated with eBMD; (2) gene-based tests using GWAS results to identify genes associated with eBMD, and (3) competitive gene set analysis (GSA) to determine if gene programs are enriched with eBMD-associated genes.

GWAS of eBMD encompassed 448,010 individuals from the UK-Biobank study (Supplementary Table 6). This included an additional 21,186 individuals, analysis of 36% more genetic variants and more detailed analysis of the X chromosome than was possible in our previous GWAS^8^. 169,618 variants were associated at genome-wide significance (*P*<6.6×10^-9^) ^7^. Twenty-five variants were classified as high impact and predicted to alter gene function (Supplementary Table 1. Glossary, Fig. 3a; Supplementary Table 6). This included variants in genes such as *MEPE*, which has been reported to be associated with low eBMD^37^, and variants associated with increased eBMD including a ‘stop/lost’ variant in the NHERF family PDZ scaffold protein 2 (*NHERF2,* formerly *SLC9A3R2*), a regulator of sodium transport^38^ (Fig. 3a,b; Supplementary Table 6).

**Fig. 3.**
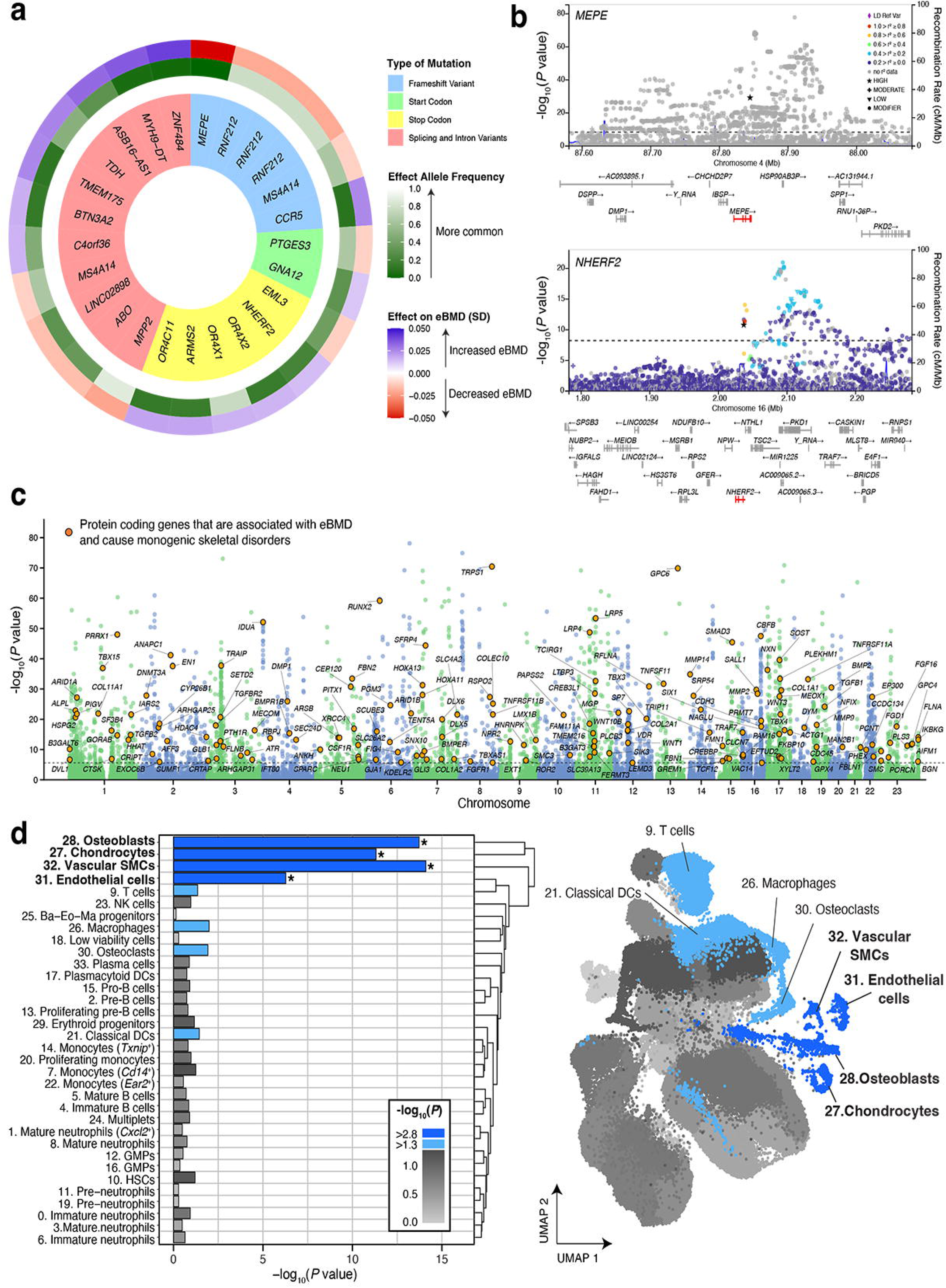
Gene programs of specific cell types are enriched with genes associated with eBMD. (**a**) Circos plot of high impact variants that are predicted to alter gene function. The inner ring denotes the types of variant, the middle ring denotes the frequency of the alternate allele, and the outer ring denotes the effect of the alternate allele on eBMD. (**b**) LocusZoom plots showing eBMD associated variants mapping closest to *MEPE* and *NHERF2*. High impact variants are represented by stars. Pairwise linkage disequilibrium (LD) could not be estimated for the lead eBMD associated variant mapping to *MEPE*. (**c**) Manhattan plot summarising the results of the genome-wide gene-based tests of eBMD conducted in the UK Biobank study using MAGMA. The dotted line denotes the threshold of statistical significance (*P* < 2.5 × 10^-6^). Orange circles indicate genes that are associated with eBMD and are monogenic skeletal disorder genes. (**d**) Bar plot and UMAP plot showing enrichment of eBMD-associated genes in gene programs for different cell types. Scale bar in bar plot and UMAP plot indicates the *P* value. Light blue bars in bar plot and light blue dots in UMAP correspond to observations that have nominal evidence of enrichment: *P* value of <0.05 [-log_10_(*P* value) of >1.3]. Dark blue bars and asterisks in bar plot and dark blue dots in UMAP correspond to observations that have robust evidence of enrichment and meet the *Bonferroni*-corrected significance threshold: *P* value of < 1.5 ×10^-3^ [-log_10_(*P* value) > 2.8].

Fine mapping identified 1,294 statistically independent lead genetic variants, defined as variants most strongly associated with eBMD that mapped closest to 903 unique protein-coding genes (Extended Data Fig. 6a; Supplementary Table 6). Sixteen lead variants represented novel associations, with 13 mapped to the X chromosome (Extended Data Fig. 6a). Three were located closest to genes involved in monogenic disorders with abnormal BMD, including: plastin-3 (*PLS3),* which causes X-linked juvenile osteoporosis^39^; inhibitor of nuclear factor kappa B kinase regulatory subunit gamma (*IKBKG)*, which causes osteopetrosis^40^; and phosphate regulating endopeptidase x-linked (*PHEX),* which causes abnormal mineralisation and X-linked hypophosphatemic rickets (Extended Data Fig. 6b; Supplementary Table 6) ^41^. The set of protein coding genes located closest to lead variants was enriched with genes that cause rare skeletal disorders (*P*=2.1×10^-19^) (Extended Data Fig. 6c; Supplementary Table 6). These included genes involved in disorders with abnormal BMD, such as OI and low bone mass, sclerosing bone disorders, and osteopetrosis. Thus, our new GWAS extends the identification of new eBMD-associated variants from 1,103 (518 loci) ^8^ to 1,294 (533 loci) and highlights causal variants and new genes that may regulate skeletal integrity.

Since gene-based tests of association offer a more powerful approach to estimate gene-trait associations than GWAS, we conducted a Multi-marker Analysis of GenoMic Annotation (MAGMA) analysis^42^. Gene-based tests detected robust eBMD associations for 3,883/19,695 protein-coding genes (*P*=2.5×10^-6^, Fig. 3c; Supplementary Table 6). 788 of the eBMD-associated genes were among the 903 genes located closest to lead variants identified by GWAS (Supplementary Table 6). eBMD-associated genes were enriched with genes that cause skeletal disorders (Supplementary Fig. 3a; Supplementary Table 6). This included genes that cause disorders with high and low BMD. Independent analysis of pulse rate, a trait unrelated to the skeleton, showed no enrichment for causative genes, suggesting enrichment was not a chance effect (Supplementary Fig. 3b; Supplementary Table 6).

GSA showed that the gene programs of osteoblast lineage cells, chondrocytes, endothelial cells and VSMCs, were enriched with eBMD-associated genes (Fig. 3d; Supplementary Table 6). Analysis of individual stages of cell differentiation showed that all osteoblast, chondrocyte, VSMC and one of four endothelial cell states were enriched with eBMD-associated genes (Extended Data Fig. 7; Supplementary Table 6).

Since 57% of genes in non-haematopoietic cell gene programs were present in at least one other gene program, we performed pairwise conditional GSA to determine whether enrichment was confounded by genes shared between gene programs (Supplementary Fig. 4a,b; Supplementary Table 6). Osteoblast lineage cells, chondrocytes, endothelial cells and VSMCs remained enriched with genes associated with eBMD (*P*_conditional_<0.05). Cells at all stages in the osteoblast lineage and hypertrophic chondrocytes remained enriched with genes associated with eBMD (*P*_conditional_<0.05; Supplementary Table 6). Given a subset of genes strongly associated with eBMD could influence enrichment, we also performed a post-hoc permutation analysis. 93–99% of genes contributed to the enrichment, suggesting that small numbers of genes were not unduly influential (Supplementary Fig. 4c).

Together, these data demonstrate that the gene programs of osteoblast lineage cells, chondrocytes, endothelial cells and VSMCs, are enriched with genes associated with eBMD and susceptibility to common polygenic skeletal diseases.

### Non-haematopoietic cells and osteoclasts are enriched with eBMD effector genes that control bone structure

We next investigated whether eBMD-associated genes in non-haematopoietic cell and osteoclast programs influence skeletal structure, as they were implicated in monogenic and polygenic skeletal diseases. We screened the MGI database to identify 1347 protein-coding genes with human orthologs that cause ‘abnormal bone structure’ when mutated in mice (Supplementary Table 1. Glossary; Supplementary Note 4). Gene programs of osteoblasts, chondrocytes, osteoclasts, and VSMCs, but not endothelial cells, were enriched with these ‘abnormal bone structure’ genes (Extended Data Fig. 8a; Supplementary Table 7). Magnitude of enrichment increased when the analysis was restricted to eBMD-associated genes from the MAGMA analysis (Extended Data Fig. 8a,d; Supplementary Table 7). Sensitivity analyses to exclude the possibility that genes causing monogenic skeletal disorders were responsible for enrichment indicated that enrichment persisted for osteoblasts, chondrocytes, and VSMCs (Extended Data Fig. 8b; Supplementary Table 7). Lastly, we showed that the magnitude of enrichment increased for genes with greatest expression in osteoblast, chondrocyte, and osteoclast gene programs (Extended Data Fig. 8c,e; Supplementary Table 7). Enrichment increased further when restricted to genes associated with eBMD (Extended Data Fig. 8c,e; Supplementary Table 7).

### Non-haematopoietic cells and osteoclasts express novel eBMD effector genes that regulate bone structure and function

Since the MGI database typically lacks biomechanical measurements of bone, we determined whether eBMD-associated genes expressed in non-haematopoietic cells and osteoclasts have functional roles in the skeleton and are indeed eBMD effector genes. We screened adult female mice from a cohort of over 1000 mouse lines with single gene deletions that have undergone detailed structural and functional skeletal phenotyping by our Origin of Bone and Cartilage Disease (OBCD) study for genes present in each of the five gene programs (Fig. 4a). This is a significant extension to the original report of 100 lines^43^ and the 596 lines reported in 2019^8^. The detailed functional bone phenotyping and phenotype definitions are described in Methods and Supplementary Table 1.

**Fig. 4.**
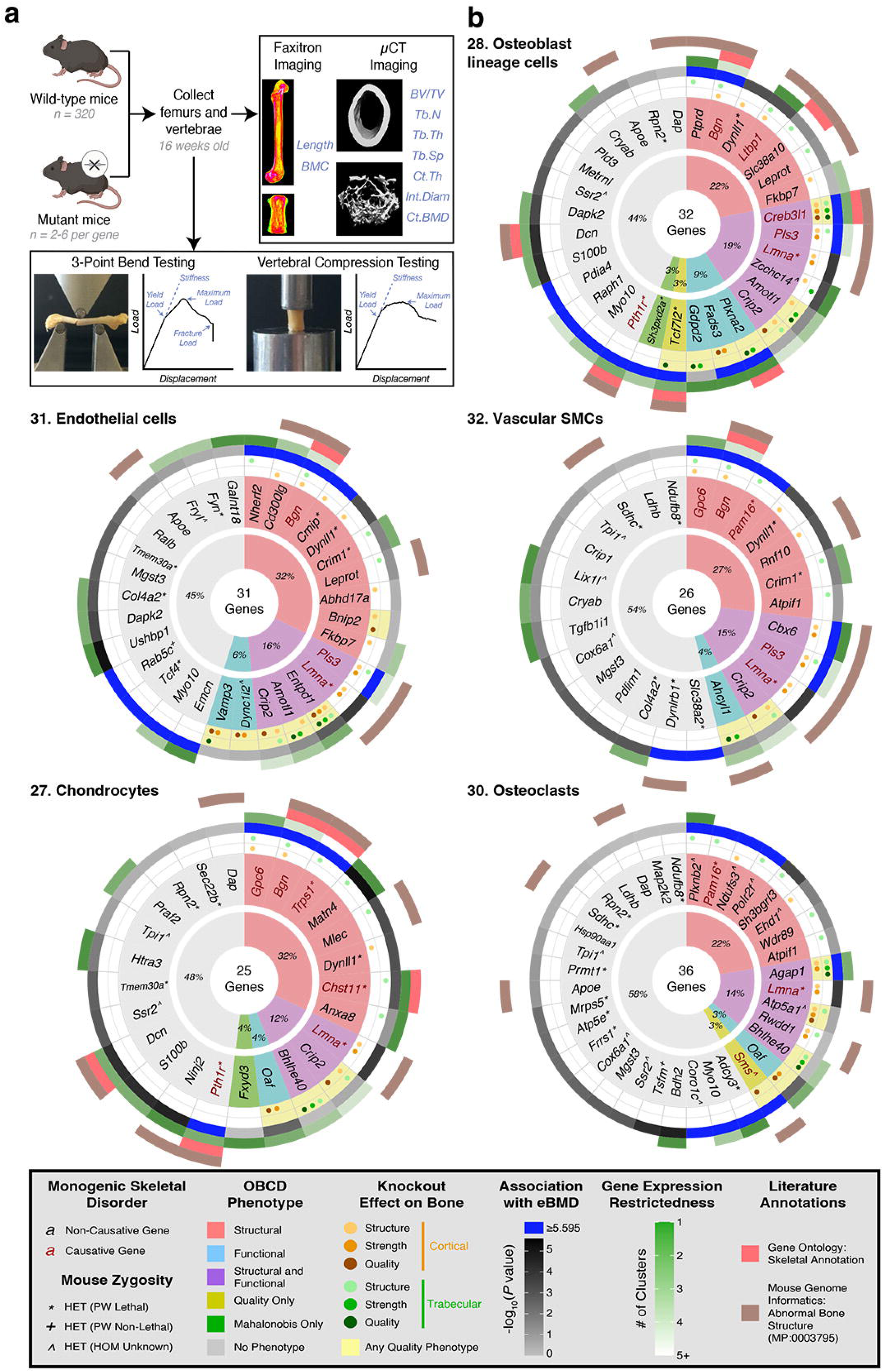
Genes that cause abnormal skeletal phenotypes when deleted in mice. (**a**) The Origins of Bone and Cartilage Disease phenotyping platform. Created with BioRender (www.biorender.com) with permission to publish. Parameters for faxitron and micro-CT imaging are BMC = bone mineral content; BV/TV = trabecular bone volume/tissue volume; Tb.N = trabecular number; Tb.Th = trabecular thickness; Tb.Sp = trabecular separation; Ct.Th = cortical thickness; Int.Diam = internal diameter of bone; Ct.BMD = cortical bone derived BMD. (**b**) The effect of deleting individual genes found in the gene programs of non-haematopoietic cells and osteoclasts in mice on the phenotype of the skeleton. Each circos plot represents genes from a single cell type and the number corresponds to the cell clusters identified in Fig. 1. The total number of genes deleted in the gene programs that define individual cell types is noted in the centre of each plot. The central ring shows the proportion of mice (%) with specified skeletal phenotypes. The next ring shows the individual genes deleted in each gene program. The nature of the skeletal phenotype is colour coded according to the key. Gene names in red are those that also cause monogenic skeletal disorders. For instances where heterozygous knockout mice were used, the lethality of homozygous knockouts are shown by the indicated symbols. The next two rings show the impact of gene deletion on cortical and trabecular bone. This is followed by information on which genes are associated with eBMD from the MAGMA analysis (grey and blue ring) and which are restricted to a single cell type (green ring). The outer two rings identify genes that appear in the GO with a skeletal term and in MGI with abnormal bone structure. The key summarises the colour coding for each phenotype, association with monogenic diseases and eBMD, gene expression restrictedness and skeleton-related annotations.

Of the 1886 genes in the five gene programs, 101 genes (5.3%) had corresponding deletions in the OBCD study (Fig. 4b), including 32 in the osteoblast program, 25 in chondrocytes, 31 in endothelial cells, 26 in VSMCs, and 36 in osteoclasts. Fifty-one (50.5%) of the 101 knockout lines exhibited an abnormal structural, functional, or bone quality phenotype (Supplementary Table 1 Glossary for definitions; Fig. 4b; Supplementary Table 8).

Some genes had established skeletal roles, including those that cause skeletal disorders and/or are associated with eBMD, such as *Bgn* (biglycan) ^44^, *Creb3l1* (cAMP responsive element binding protein 3 like 1) ^45^, *Gpc6* (glypican 6) ^7^ and *Pls3*^39^. *Pls3* is an example of a disease-causing gene for which the cellular and molecular mechanisms of disease pathogenesis are unclear. Many other identified genes were not annotated with a skeletal term in the GO database or had no or limited description of an impact on bone structure in the MGI database, suggesting they are previously unknown regulators of bone structure and/or function. These included cell type-restricted genes like *Ptprd* (protein tyrosine phosphatase receptor type D, in osteoblasts), *Nherf2* (in endothelial cells), *Cbx6* (chromobox 6, in VSMCs), and *Agap1* (ArfGAP With GTPase Domain, Ankyrin Repeat And PH Domain 1, in osteoclasts) (Fig. 4b; Supplementary Table 8). This analysis suggests that the gene programs defining cells in the endosteal compartment include many new and important eBMD effector genes.

#### PLS3 is a widely expressed eBMD effector gene that controls bone structure and function

To explore an effector gene that is expressed widely, we performed studies on *Pls3*. While *PLS3* loss-of-function mutations cause OI and X-linked osteoporosis in children^39^, we also found its common genetic variation is associated with eBMD in adults (Fig. 3c; Extended Data Fig. 6a; Supplementary Table 6). *Pls3* was expressed in the osteoblast lineage cells, endothelial cells, and VSMCs in bone (Fig. 5a), and in epithelial, mesenchymal, and vascular cells outside the skeleton (Supplementary Fig. 5; Supplementary Table 9). Whilst *Pls3* has been studied in several of these cell types, it is still unclear which cells it acts through to affect bone integrity^46–49^.

**Fig. 5.**
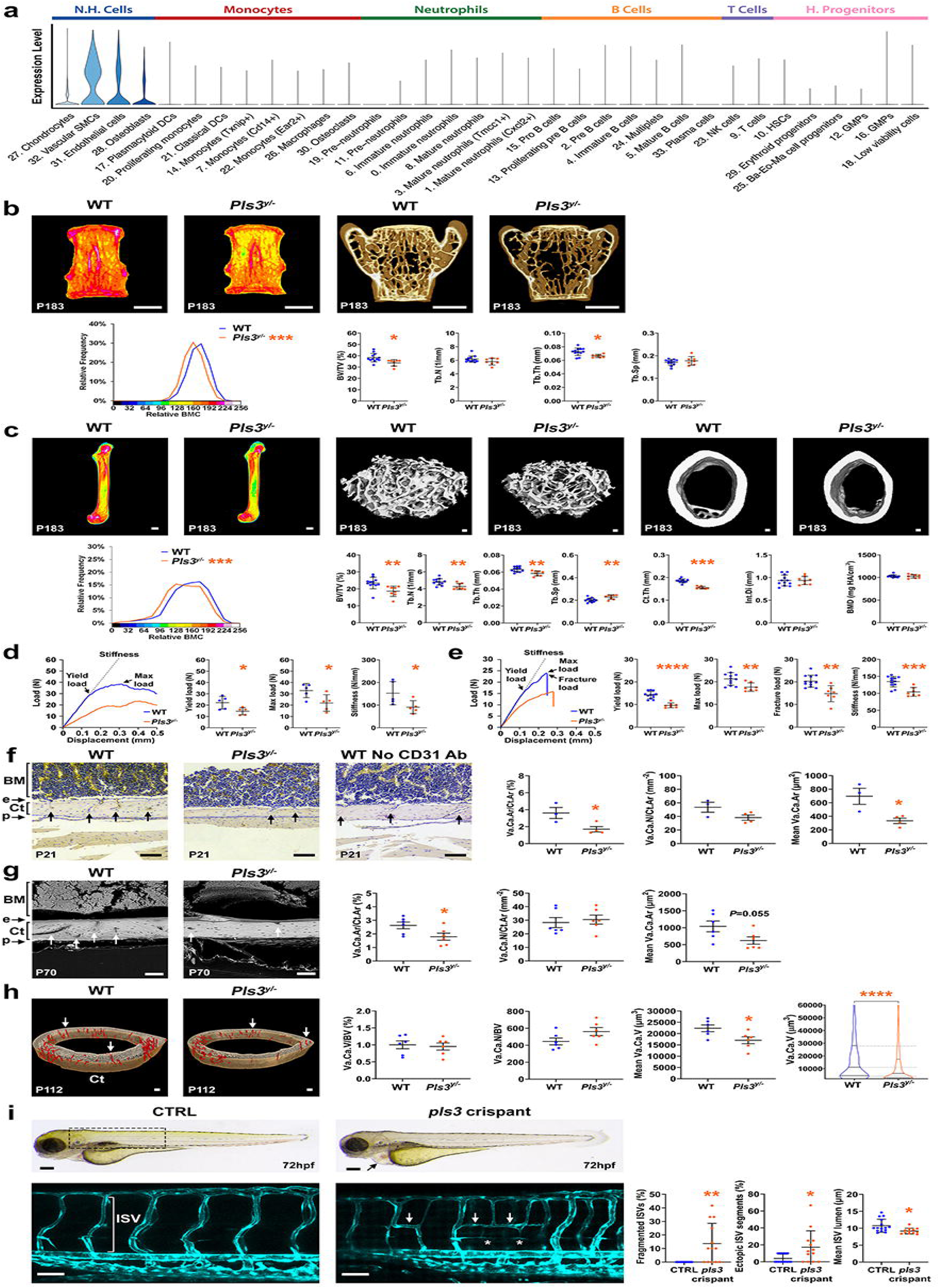
Skeletal phenotype of male *Pls3^y/-^* mice. (**a**) Violin plots showing *Pls3* gene expression in the endosteal bone and bone marrow scRNAseq dataset. N.H. = non-haematopoietic. (**b**) Pseudocoloured X-ray microradiography of lumbar vertebrae (L5) from P183 (WT n=12; *Pls3^y/-^* n=8) mice; scale bar = 1mm. Relative frequency histograms of BMC are shown for each age comparison; Kolmogorov-Smirnov test, ***P<0.001. Micro-CT images of mid-coronal sections of lumbar vertebrae (L5) from P183 (WT n=12; *Pls3^y/-^* n=7) mice; scale bar = 1mm. Graphs show bone volume as a proportion of tissue volume (BV/TV), trabecular number (Tb.N), trabecular thickness (Tb.Th), and trabecular separation (Tb.Sp); mean±SD; Student’s *t*-test; \**P*<0.05. (**c**) Pseudocoloured X-ray microradiography of femurs from P183 (WT n=12; *Pls3^y/-^*n=8) mice. Low bone mineral content (BMC) is blue/green and high BMC is pink; scale bar = 1mm. Relative frequency histograms of BMC are shown for each age comparison; Kolmogorov-Smirnov test, ***P<0.001. Micro-CT images of distal femur trabecular bone from P183 (WT n=12; *Pls3^y/-^* n=8) mice; scale bar = 100μm. Graphs show bone volume as a proportion of tissue volume (BV/TV), trabecular number (Tb.N), trabecular thickness (Tb.Th), and trabecular separation (Tb.Sp); mean±SD; Student’s *t*-test; \*\**P*<0.01. Micro-CT images of femur mid-shaft cortical bone from P183 (WT n=12; *Pls3^y/-^* n=8) mice; scale bar = 100μm. Graphs show cortical thickness (Ct.Th), internal diameter (Int.Di), and bone mineral density (Ct.BMD); mean±SD; Student’s *t*-test; \*\*\**P*<0.001. (**d**) Representative load displacement curves from compression testing of lumbar vertebrae (L5) from P183 (WT n=5; *Pls3^y/-^* n=5) mice. Graphs show yield load, maximum load, and stiffness; mean±SD; Student’s *t*-test; \**P*<0.05. (e) Representative load displacement curves from three-point bend testing of femurs from P183 (WT, n=12; *Pls3^y/-^*, n=7) mice. Graphs show yield load, maximum load, fracture load, and stiffness; mean±SD; Student’s *t*-test; \*\**P*<0.01, \*\*\**P*<0.001. (**f**) Anti-CD31 immunohistochemistry of mid-femur sections from P21 (WT n=3; *Pls3^y/-^* n=4) mice; scale bar = 100μm. BM (bone marrow); Ct (cortical bone); e (endosteum); p (periosteum); black arrows indicate cortical vascular canals. Graphs show vascular canal area as a proportion of cortical area (Va.Ca.Ar/Ct.Ar), vascular canal number per cortical area (Va.Ca.N/Ct.Ar), and mean vascular canal area (Mean Va.Ca.Ar); mean±SD; Student’s *t*-test;. (**g**) Backscattered electron SEM images of triiodide stained mid-femur sections from P70 (WT n=6; *Pls3^y/-^* n=6) mice; scale bar = 100μm. BM (bone marrow); Ct (cortical bone); e (endosteum); p (periosteum); white arrows indicate cortical vascular canals. Graphs show vascular canal area as a proportion of cortical area (Va.Ca.Ar/Ct.Ar), vascular canal number per cortical area (Va.Ca.N/Ct.Ar), and mean vascular canal area (Mean Va.Ca.Ar); (**h**) Micro-CT images of a 250µm mid-femur ROI from P112 (WT n=6; *Pls3^y/-^*n=6) mice; showing cortical vessels in red; scale bar = 100μm. Ct (cortical bone); white arrows indicate examples of cortical vascular canals. Graphs show vascular canal volume as a proportion of bone volume (Va.Ca.V/BV), vascular canal number per bone volume (Va.Ca.N/BV), mean vascular canal volume (Mean Va.Ca.Ar); mean±SD; Student’s *t*-test;. \**P*<0.05. The violin plot shows the distribution of Va.Ca.V; Kolmogorov-Smirnov test; \*\*\*\**P*<0.0001. (**i**) Upper images show overall body morphology of uninjected control (CTRL) and *pls3 crispant* zebrafish embryos at 72 hours post fertilisation (hpf); scale bar = 100μm. Black arrow indicates pericardial oedema in *pls3 crispant*. Box shows region of interest (ROI) represented in lower images. Lower images are representative maximum intensity projections of *Tg(kdr-l:eGFP)* expression in the trunk vasculature and white bar indicates intersegmental vessel (ISV) in CTRL at 72 hpf. (CTRL n=15; *pls3 crispants* n=13); scale bar = 50μm. White asterisks indicate incomplete ISVs, disconnected from the dorsal aorta or posterior cardinal vein. White arrows indicate ectopic ISV segments connecting neighbouring ISVs in *pls3 crispants*. Graphs show percentage of fragmented ISVs, percentage of ectopic vessel segments, and mean ISV luminal diameter; mean±SD; Student’s *t*-test;. \**P*<0.05, ** *P*<0.01.

As X-linked osteoporosis is characterised by multiple vertebral fractures in boys, we characterised lumbar vertebrae from male *Pls3* deficient male mice (*Pls3^y/-^*). Bone mineral content (BMC) was reduced in vertebrae at both P70 and P183 in *Pls3^y/-^* mice, while micro-CT analysis demonstrated decreased bone volume (BV/TV) and trabecular thickness (Tb.Th) in *Pls3^y/-^* vertebrae at P183 (Fig. 5b; Supplementary Fig. 6a). In long bones, *Pls3^y/-^* mice also showed decreased BMC, reduced BV/TV, trabecular number (Tb.N), and Tb.Th and increased trabecular spacing (Tb.Sp) at P183 (Fig. 5c; Supplementary Fig. 6c,d). Cortical thickness (Ct.Th) was also reduced at P70 and P183 in *Pls3^y/-^* mice (Fig. 5c; Supplementary Fig. 6e). Biomechanical testing demonstrated maximum load was impaired in vertebrae at P70, and yield load, maximum load, and stiffness were all reduced at P183 in *Pls3^y/-^*mice (Fig. 5d; Supplementary Fig. 6b). Long bones also exhibited a reduced yield load, maximum load, and stiffness at P70 and P183 and impaired fracture load at P183 in *Pls3^y/-^* mice (Fig. 5e; Supplementary Fig. 6f). Female *Pls3* deficient (*Pls3^-/-^*) mice demonstrated a similar osteoporotic phenotype (Supplementary Fig. 7).

To investigate the underlying cellular mechanism of disease pathogenesis, we analysed the major bone cell lineages. Analysis of the postnatal day 1 (P1) skeleton revealed no differences in endochondral ossification between *Pls3^y/-^* and WT mice (Extended Data Fig. 9a). There were no differences in growth plate reserve, proliferative and hypertrophic zones among P21 *Pls3^y/-^* and WT mice (Extended Data Fig. 9a). Linear growth between P1 and P21 did not differ between *Pls3^y/-^* and WT mice (Extended Data Fig. 9a). Static and dynamic histomorphometry and serum measures of osteoclastic resorption, osteoblastic bone formation and osteocyte-derived proteins were unaffected in *Pls3*^y/−^ mice (Extended Data Fig. 9b-d). Cortical porosity and osteocyte lacuna size and number were increased in *Pls3^y/-^* mice (Extended Data Fig. 9e).

Since *Pls3* was most highly expressed in VSMCs and endothelial cells in bone (Fig. 5a), we examined whether *Pls3* deletion affected the vasculature in bone. *Pls3^y/-^* mice showed smaller blood vessels in bone compared to control mice (Fig. 5f), which was confirmed by backscattered electron SEM, and micro-CT analysis (Fig. 5g,h; Supplementary Fig. 8). Furthermore, knockdown of *pls3* in zebrafish embryos using a *Crispr*-cocktail approach^50^ resulted in mild cardiac oedema and significant defects in intersegmental vessel (ISV) formation, including smaller lumens, disconnections from the dorsal aorta or posterior cardinal vein, and ectopic vessel connections compared to controls. The absence of bone cells at this early developmental stage of zebrafish indicates *pls3* might act directly to control angiogenesis and vascular patterning (Fig. 5i). These data suggest that the severe phenotype of X-linked osteoporosis might result from abnormalities in the bone vasculature, rather than a defect in cells of any one individual bone cell lineage.

#### eBMD effector genes with restricted expression determine bone structure and function

To understand the impact of eBMD-associated genes with restricted expression in individual skeletal cell types, we investigated the impact of deleting *Ptprd*, *Nherf2*, *Cbx6*, and *Agap1* on bone structure and function. *Ptprd* expression was restricted to osteoblast lineage cells in the skeleton (Fig. 6a) but was expressed in mesenchymal, epithelial and glial cells in the lung, brain, and ovary (Supplementary Fig. 5; Supplementary Table 9). P112 female *Ptprd*^−/−^ mice had increased vertebral BMC and decreased vertebral length compared to the wild-type (WT) mice reference range derived from 320 P112 female mice from an identical genetic background housed in the same animal facility (Fig. 6b,c). Femurs from *Ptprd*^−/−^ mice showed decreased BV/TV and Tb.N with increased Tb.Sp but normal cortical bone parameters (Fig. 6d,e). Femoral and vertebral bone strength were unchanged in *Ptprd*^−/−^ mice (Fig. 6f,g). *Nherf2* expression was restricted to endothelial cells in the endosteal compartment (Fig. 6a) and also expressed in endothelial cells in other tissues (Supplementary Fig. 5; Supplementary Table 9), suggesting this gene is an important regulator of endothelial cell function. *Nherf2*^−/−^ mice had decreased vertebral length (Fig. 6b,c) and reduced femoral BV/TV and Tb.N with increased Tb.Sp (Fig. 6d,e). Femoral and vertebral bone strength and stiffness parameters were unchanged in *Nherf2*^−/−^ mice (Fig. 6f-g). *Cbx6* expression was restricted to VSMCs in the skeleton (Fig. 6a) but also expressed in the lung, brain, and pancreas and stomach (Supplementary Fig. 5; Supplementary Table 9). *Cbx6*^−/−^ mice exhibited decreased vertebral length (Fig. 6b,c) and reduced femoral cortical thickness (Ct.Th) (Fig. 6d,e). Accordingly, femur yield load and maximum load were decreased compared to WT mice, while vertebral strength was not different from control (Fig. 6f,g). Finally, *Agap1* expression was primarily restricted to osteoclasts in bone but also expressed in the heart, brain, and liver (Supplementary Fig. 5; Supplementary Table 9). *Agap1^-/-^* mice had decreased femur BMC and decreased vertebral BMC and length, compared to WT mice (Fig. 6b-c). Femurs from *Agap1^-/-^* mice also had decreased BV/TV and Ct.Th (Fig. 6d-e). Femur maximum load and vertebral yield load, maximum load, and stiffness were all decreased compared to WT mice (Fig. 6f-g).

**Fig. 6.**
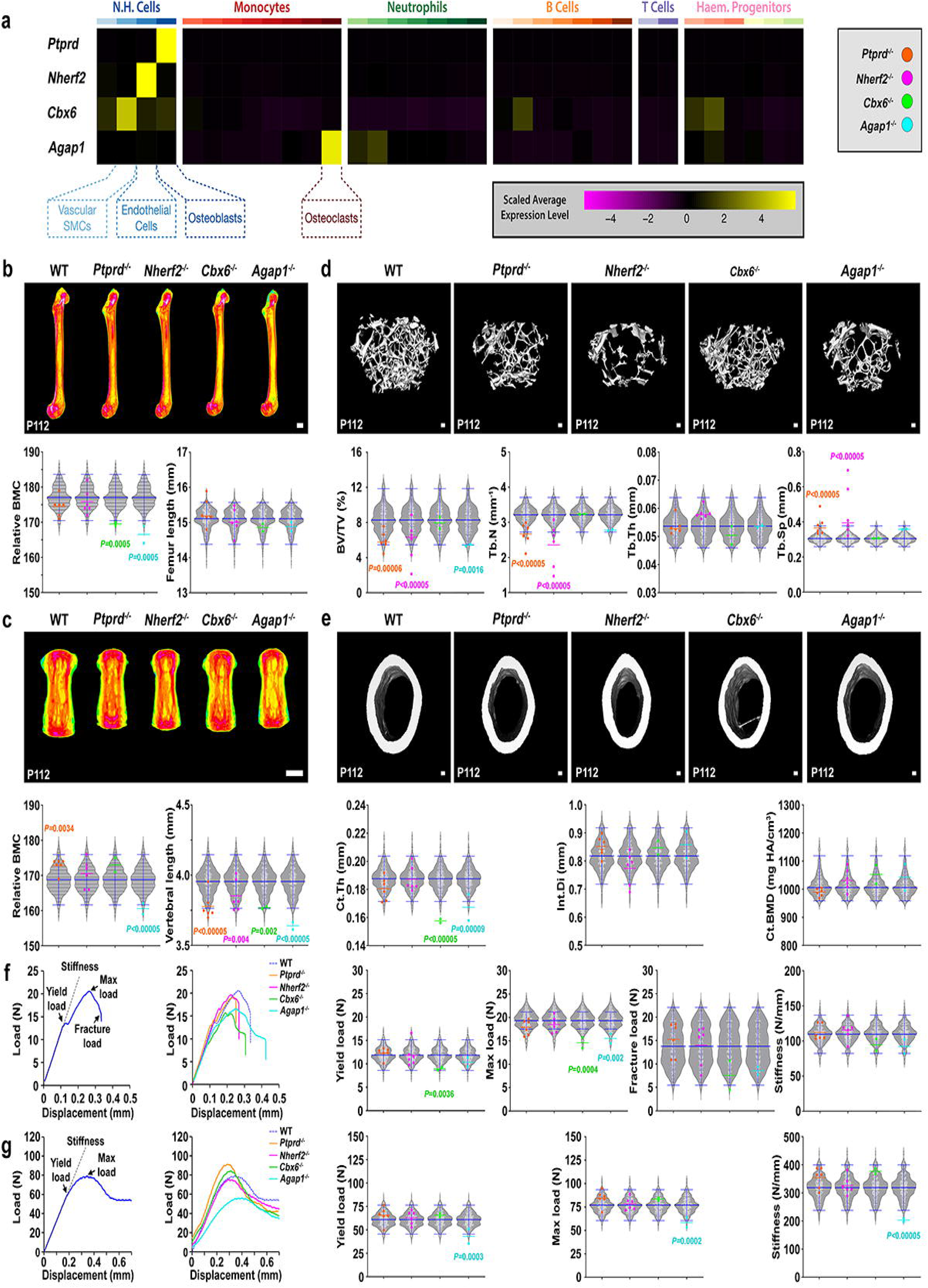
Skeletal phenotypes of female *Ptprd^-/-^*, *Cbx6^-/-^*, *Nherf2^-/-^*, and *Agap1^-/-^* mice. (**a**) Heatmaps showing *Ptprd*, *Nherf2*, *Cbx6,* and *Agap1* gene expression in the endosteal bone and bone marrow scRNA-seq dataset. N.H. = non-haematopoietic. (**b**) Pseudocoloured X-ray microradiography of femurs from P112 WT, *Ptprd^-/-^*, *Nherf2^-/-^*, *Cbx6^-/-^*, and *Agap1^-/-^* mice. Low bone mineral content (BMC) is blue/green and high BMC is pink; scale bar = 1mm. Graphs show relative bone mineral content (BMC, mean±SD) and femur length (median±2.5^th^ and 97.5^th^ percentiles) with WT reference range represented as grey violin plots with individual values shown as white dots (n=320). Individual values, together with the mean or median, from *Ptprd^-/-^* (orange n=6), *Nherf2^-/-^* (pink n=6), *Cbx6^-/-^*(green n=2), and *Agap1^-/-^* (cyan n=2) mice are shown. Significant *P* values after permutation testing are indicated. (**c**) Caudal vertebrae from P112 WT, *Ptprd^-/-^*, *Nherf2^-/-^*, *Cbx6^-/-^*, and *Agap1^-/-^* mice; scale bar = 1mm. Graphs show BMC (mean±SD), and vertebral length (mean±SD) with WT reference range represented as grey violin plots with individual values shown as white dots (n=320). Individual values and mean from *Ptprd^-/-^* (orange n=6), *Nherf2^-/-^* (pink n=6), *Cbx6^-/-^* (green n=2), and *Agap1^-/-^* (cyan n=2) mice are shown. Significant *P* values after permutation testing are indicated. (**d**) Micro-CT images of distal femur trabecular bone from P112 WT, *Ptprd^-/-^*, *Nherf2^-/-^*, *Cbx6^-/-^*, and *Agap1^-/-^* mice; scale bar = 100µm. Graphs show bone volume as a proportion of tissue volume (BV/TV, mean±SD), trabecular number (Tb.N, mean±SD), trabecular thickness (Tb.Th, mean±SD), and trabecular separation (Tb.Sp, median±2.5^th^ and 97.5^th^ percentiles). Significant *P* values after permutation testing are indicated. (**e**) Micro-CT images of femur mid-shaft cortical bone from P112 WT, *Ptprd^-/-^*, *Nherf2^-/-^*, *Cbx6^-/-^,* and *Agap1^-/-^*mice; scale bar = 100µm. Graphs show cortical thickness (Ct.Th, mean±SD), internal diameter (Int.Di, mean±SD), and bone mineral density (Ct.BMD, median±2.5^th^ and 97.5^th^ percentiles). Significant *P* values after permutation testing are indicated. (**f**) Example (left) and representative (right) load displacement curves from three-point bend testing of femurs from P112 WT (dotted blue), *Ptprd^-/-^* (orange), *Nherf2^-/-^* (pink), and *Cbx6^-/-^*(green), and *Agap1^-/-^* (cyan) mice. Graphs show yield load (mean±SD), maximum load (mean±SD), fracture load (median±2.5^th^ and 97.5^th^ percentiles), and stiffness (mean±SD). Significant *P* values after permutation testing are indicated. (**g**) Example (left) and representative (right) load displacement curves from compression testing of caudal vertebrae from P112 WT (dotted blue), *Ptprd^-/-^* (orange) *Nherf2^-/-^*(pink), and *Cbx6^-/-^* (green), and *Agap1^-/-^*(cyan) mice. Graphs show yield load (mean±SD), maximum load (mean±SD), and stiffness (mean±SD). Significant *P* values after permutation testing are indicated.

### Mapping the cell types and genes involved in skeletal diseases in human bone

To investigate the translational importance of our findings we further analysed scRNA-seq data from human bone. We isolated and sequenced 125,063 cells from adult human femoral head bone from individuals undergoing hip replacement surgery (Fig. 7a). Human bone included similar populations of MSCs, pre-osteoblasts, mature osteoblasts, chondrocytes, fibroblasts, ECs and VSMCs to those found in mice (Fig. 7a,b; Supplementary Table 2). The human dataset also included haematopoietic cells including HSCs, GMPs, neutrophils, monocytes and macrophages, B cells and T and NK cells.

**Fig. 7.**
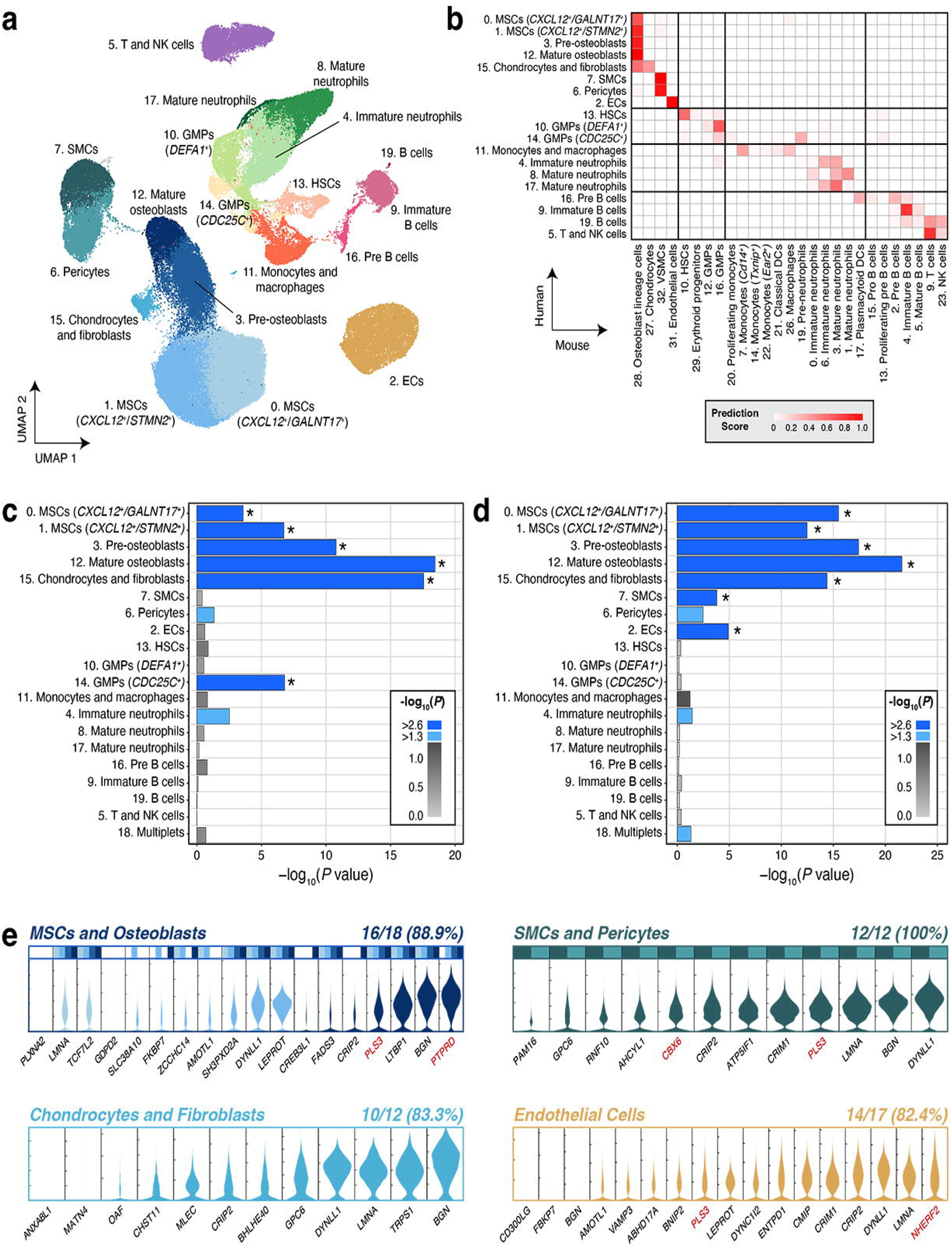
Mapping the cell types and genes involved in skeletal diseases in human bone. (**a**) UMAP plot derived from scRNA-seq data of cells isolated from human femoral head showing 20 cell clusters. (**b**) Label transfer analysis using mouse scRNA-seq data from Fig. 1 as a reference to compare the gene expression with human scRNA-seq data. Prediction scores reflect the probability assigned to a transferred label for a given query cell cluster. Bar plots showing enrichment of gene programs for (**c**) causative genes of monogenic skeletal disorders and (**d**) eBMD-associated genes. Scale bars in bar plots indicate the *P* value. Light blue bars in bar plot correspond to observations that have nominal evidence of enrichment: *P* value of <0.05 [-log_10_(*P* value) of >1.3]. Dark blue bars and asterisks in bar plot correspond to observations that have robust evidence of enrichment and meet the *Bonferroni*-corrected significance threshold: *P* value of < 2.5 ×10^-3^ [-log_10_(*P* value) > 2.6]. (**e**) Violin plots showing expression of exemplar genes (cell type-specific genes that are associated with eBMD and functionally validated in mice) in human bone. Where multiple clusters are present for a lineage, a single violin is shown corresponding to the cluster with the highest expression, with bars above indicating which clusters pass the threshold for active expression of the gene (see Methods), both coloured according to the UMAP in (**a**). Numbers above the plots for each lineage indicate the proportion of genes that pass the threshold for active expression. Selected exemplars from Fig. 5 and Fig. 6 are highlighted in red.

In common with the murine data, the gene programs that define human MSCs, pre-osteoblasts, mature osteoblasts, chondrocytes and fibroblasts clusters were enriched for genes that cause both monogenic skeletal disorders and are associated with eBMD (Fig. 7c,d; Supplementary Tables 5,6). This was confirmed in our analysis of osteoblasts from human embryonic joints and in cranium development^51^ (data not shown). VSMCs and ECs were also enriched for eBMD-associated genes (Fig. 7d). The majority of the eBMD effector genes that we functionally validated in mice from individual cell types (82-100%), were also found in their cellular counterparts in human adult bone (Fig 7e). This included *PLS3* in MSCs, osteoblasts, SMCs and pericytes, and ECs; *PTPRD* in MSCs and osteoblasts; *CBX6* in SMCs and pericytes; and *NHERF2* in ECs (Fig. 7e).

Finally, to validate our human scRNA-seq data, and examine cells not readily identified by microfluidics-based methods, we interrogated a novel spatial transcriptomic data of adult human bone^52^. In addition to osteoblasts, chondrocytes, cells of the vasculature and osteoclasts, this approach also identified a population of adipocytes (Extended Data Fig. 10a,b; Supplementary Table 2). The gene programs that define osteoblasts, chondrocytes and osteoclasts were each enriched for genes that caused monogenic skeletal disorders and eBMD-associated genes (Extended Data Fig. 10c,d). In common with scRNA-seq data, VSMCs were also enriched for eBMD-associated genes (Extended Data Fig. 10d). Adipocytes were not enriched for either genes that cause monogenic skeletal disorders or eBMD-associated genes. This is consistent with findings in adipocytes derived from the periosteum of mouse bones^53^ (data not shown). Together these findings demonstrated this approach has important translational potential.

### A platform for real time interrogation of multiscale data of genes that regulate eBMD

Our multiscale cross species analysis provides a systematic approach to identify the cells and genes with important functional roles in the skeleton. To ensure these novel datasets are accessible, we developed a web-based platform that integrates these data to enable users to identify effector genes and formulate cellular and mechanistic hypotheses relating to their role in skeletal health and disease. The platform can be accessed using the following hyperlink: https://www.musculoskeletal-genomics.org.

## Discussion

The rapid advance in our understanding of the pathophysiology and development of new treatments for many diseases has benefited from the application of cutting-edge single-cell technologies. Despite the global burden of skeletal disease, the challenges of working with mineralised tissues and difficulties in obtaining normal human bone samples has meant skeletal research has lagged behind other fields. In this study we address this need by exploiting the value of cross-species analysis to identify common, critical and conserved causal mechanisms. We developed and optimised single-cell methodologies in skeletal tissues to define the critical endosteal compartment that regulates bone turnover in the skeleton.

This study identified the repertoire of cells within the endosteal compartment, defined the gene programs that control them and enabled detailed lineage trajectory analysis. Cell types enriched with monogenic human disorder genes and genes associated with eBMD, identified in a new GWAS of 448,010 individuals that now includes data from the X-chromosome, were considered disease relevant and investigated for causality. Our analysis validated the key role of osteoblast, chondrocyte, and monocyte/macrophage/osteoclast lineages in skeletal pathophysiology but also identified previously underappreciated cell types including endothelial cells and VSMCs in both murine and human bone. Many of these cell types were localised to the endosteal compartment, and in some cases to different endosteal compartments, illustrating that this is a distinct and specialised location within bone.

Our identification of a role for endothelial cells and VSMCs in skeletal disease is exemplified by the finding of a novel abnormal vascular phenotype in the cortical bone of *Pls3^y/-^* mice and during *pls3* deficient zebrafish embryo development. The strength of our approach is highlighted by the fact that mutations of *PLS3* have been known to cause X-linked osteoporosis for more than a decade but the cellular and molecular mechanism of disease has remained elusive^39^. Furthermore, most genes defining disease relevant cells in the endosteal compartment have underappreciated or unknown roles in skeletal pathophysiology. The expression of these genes may be restricted to individual cell types or show broader expression patterns. Here we demonstrate causation for multiple exemplar genes by characterising skeletal phenotypes in knockout mice. Our results suggest that the pathological mechanisms underlying skeletal disease may be lineage restricted, involve contributions from multiple cell types, or may reflect complex systemic effects. To investigate further, we characterised the cellular interactions between cells within the endosteal compartment and showed that endothelial cells and VSMCs can interact directly with cells of the osteoblast lineage. These findings provide a potential explanation for how genes expressed in endothelial cells and VSMCs, such as *Pls3*, *Nherf2* and *Cbx6,* may regulate bone mass.

Thus, our novel unbiased approach has generated a comprehensive map of cell types across the entire endosteal compartment and expands the repertoire of endosteal cells that regulate bone to include endothelial cells and VSMCs, and populations of monocytes/macrophages, neutrophils, B and T cells. This complements analyses of specific bone and marrow cell populations^54–57^. To maximise accessibility to these new data and integrate them with available datasets we developed an interactive, open access portal (www.musculoskeletal-genomics.org).

As with all novel approaches, our multiscale study has some limitations. We focused on the endosteal compartment, and this does not include cells from the periosteum. Nevertheless, mesenchymal cells from these locations have been shown to be similar^58^. The single cell isolation and sequencing methodology may not capture fragile cells such as adipocytes, multinucleated osteoclasts and cells embedded within bone, such as mature osteocytes.

Despite this, cells with canonical osteoclast markers, likely representing mononuclear osteoclasts or osteomorphs were identified^24^ and early osteocytes were also isolated, which probably represent immature cells located at the endosteal surface prior to entombment. Moreover, the application of orthogonal methods, including spatial transcriptomics, further validated our approach and enabled identification of adipocytes, although these cells were not enriched with disease-causing genes. We only studied the endosteal compartment in the femur and whilst different bones have different developmental patterns osteoblastic cells and osteocytes from different bones express remarkably similar gene programs^59,60^. Our new eBMD GWAS was generated using UK-Biobank data, which is a cohort of European ancestry, and has not been replicated in an independent cohort. Furthermore, whilst genes shared between cellular gene programs could account for enrichment, pairwise conditional gene set analysis suggested this was not the case. Finally, whilst we phenotyped more than 1000 unselected knockout out mouse lines this did not include all candidate genes. Moreover, due to the scale of the analysis, outlier phenotypes were determined by comparison of 2-6 mice from each knockout line with combined data from 320 wildtype littermates that were generated and analysed contemporaneously with the knockout lines. Outlier parameters were also independently identified by permutation testing and all lines selected for further detailed study replicated the phenotype, confirming the strength of this approach. Future studies could explore cell specificity and sexual dimorphism using inducible conditional gene targeting approaches in male and female mice, for example following *Pls3* deletion.

In summary our approach integrates new and published genetic, transcriptional and functional data from humans, mice and zebrafish, and identifies a novel cellular and molecular framework through which to better understand skeletal physiology and disease. Although the GWAS focuses on eBMD and osteoporosis, the data have far wider implications for skeletal development, rare bone disorders, aging, fracture repair, and skeletal responses to systemic diseases including malignancy and inflammation. This foundational knowledge will help to identify and prioritise new therapeutic targets for the prevention and treatment of musculoskeletal disease.

## Methods

### Mice for the isolation of bone cells and single-cell RNAseq (scRNA-seq)

9–10-week-old male wildtype C57BL/6J mice were used in the isolation of cells from different bone compartments for scRNA-seq. Animal experiments were approved by the Garvan Institute of Medical Research Animal Ethics Committee (ARA16/01, ARA19/09 and ARA_22/12).

### Isolation of cells from the endosteal and bone marrow compartments in mice

To obtain endosteal compartment and bone marrow cells for scRNA-seq, mice were sacrificed via CO_2_ asphyxia. Femurs were harvested from 5 groups of 5 mice. Soft tissue and epiphysis were removed from the femurs before being separated into diaphysis and metaphysis. Marrow cells were collected by flushing the diaphysis with PBS. Marrow-depleted diaphyseal and metaphyseal bone were crushed and cut up gently, then cells that are adherent to the endosteal surface were removed by digestion using 2mg/ml of collagenase A and 2.5mg/ml of trypsin for 30 mins at 37°C. After digestion, bone fragments were vortexed for 10s and the supernatant containing digested endosteal cells was filtered through a 100μm filter into collection tubes containing 10% fetal calf serum (FCS; Bovogen Biologicals). Marrow cells and endosteal cells were collected by centrifugation at 400x g for 5 mins and resuspended in 200μl PBS supplemented with 2% FCS prior to staining for FACS sorting.

### Isolation of cells from human femoral head bone

Femoral head bone samples were obtained from patients with osteoarthritis undergoing total hip arthroplasty with approval from the St. Vincent’s Hospital Sydney Human Research Ethics Committee (2022/ETH00475). Data for the patients and samples are provided in Supplementary Note 2. Femoral head samples were collected in RPMI medium and stored at 4°C for up to 18 hours prior to cell isolation. Bone cores were isolated from the femoral head using an electric core drill. Bone fragments from the femoral neck regions were obtained by cutting using secateurs. Bone cores and fragments were cut up into smaller bone chips using surgical blades and scissors. Cells were removed from the bone chips using 2mg/ml of collagenase A and 2.5mg/ml of dispase for 30 mins at 37°C. After digestion, bone chips were vortexed for 10s and the supernatant containing digested cells was filtered through a 100μm filter into collection tubes containing 10% fetal calf serum (FCS; Bovogen Biologicals).

### FACS enrichment of mouse and human cells

Mouse cells were stained for Ter119-PE (Biolegend) at 4°C for 30 mins and rinsed with PBS supplemented with 2% FCS. Dead cells and debris were excluded by FSC, SSC and DAPI (ThermoFisher Scientific). Cells that were viable (DAPI-negative) and negative for erythroid marker (Ter119) were sorted into PBS supplemented with 2% FCS. The gating strategy is shown in Supplementary Note 3.

Human cells were stained for CD235-BUV395 and CD45-APC-H7 (BD Biosciences) at 4°C for 30 mins and rinsed with PBS supplemented with 2% FCS. Dead cells and debris were excluded by FSC, SSC and DAPI (ThermoFisher Scientific). Cells that were viable (DAPI-negative) and negative for erythroid marker (CD235a) were then sorted based on CD45 for haematopoietic (CD45-positive) and non-haematopoietic (CD45-negative) cells into PBS supplemented with 2% FCS. Gating strategy is shown in Supplementary Note 3.

### scRNA-seq

Single cells were encapsulated into emulsion droplets using the 10x Chromium platform (10x Genomics). scRNA-seq libraries were constructed using the Chromium Single Cell 3’ v2 Reagent Kit according to the manufacturer’s protocol. Briefly, FACS sorted cells were examined under a microscope and counted with a cell counter (Thermo Fisher Scientific). Cells were loaded into each channel with a target output of 10,000 cells. Reverse transcription and library preparation were performed on a C1000 Touch Thermal cycler with 96-Deep Well Reaction Module (Bio-Rad). Amplified cDNA and final libraries were evaluated on an Agilent Tapestation using a High Sensitivity D1000 ScreenTape (Agilent Technologies). Individual libraries were diluted to 4nM and pooled for sequencing. Pools were sequenced with 75 cycle run kits (26bp Read1, 8bp Index1 and 55bp Read2) on the Novaseq Sequencing System (Illumina) to 80-90% saturation level. scRNA-seq services were provided by the Garvan Genomics Platform at the Garvan Institute of Medical Research.

### Pre-processing of 10x scRNA-seq data

Raw sequencing data were processed using the CellRanger pipeline (versions 2-7, 10x Genomics). Count matrices were loaded into R (version 4.5.1) and further processed using Seurat (versions 2-5) ^61^. For scRNA-seq data we removed all cells with fewer than 300 distinct genes or cells with more than 10% unique molecular identifiers stemming from mitochondrial genes.

### Dimensionality reduction, clustering and sub-clustering of 10x scRNA-seq data

Dimensionality reduction was performed using gene expression data for the top 3000 variable genes. The variable genes were selected based on dispersion of binned variance to mean expression ratios using the FindVariableGenes function of the Seurat package. Next, principal component analysis (PCA) was conducted, and the first 40 principal components were included for the mouse scRNA-seq data, and the first 20 principal components were included for the human scRNA-seq data, for subsequent clustering and UMAP analysis based on manual inspection of a principal component variance plot (‘PC elbow plot’). Human scRNA-seq data was batch corrected based on donor using Harmony^62^. Graph-based clustering of the PCA reduced data with the Louvain Method was performed after computing a shared nearest neighbour graph. The clusters were visualized on a 2D map produced with Uniform Manifold Approximation and Projection (UMAP). Multiple resolutions of clustering were conducted and the selected resolution represents clusters that best approximate the cell types of interest based on well characterised markers (Supplementary Note 1). For high resolution sub-clustering of the non-haematopoietic cells, the same procedure of finding variable genes, dimensionality reduction, and clustering was applied to the restricted set of data.

### Cell type annotation

Cell types were identified using the Seurat clustering algorithm as described above, using both discrete and continuous variations in gene expression across cell clusters. Cell types were annotated based on canonical markers and markers identified in published bulk RNAseq and scRNA-seq datasets (Supplementary Note 1) ^61,63–65^. For cell clusters without previous annotations from the literature, we used cluster-specific gene programs detailed below to annotate cells at the resolution of Leiden clusters.

### Defining gene programs of each cell type using differential gene expression analysis

Differentially expressed genes for each cluster were identified using the FindAllMarkers function of Seurat and ROC-based test statistics for differential expression between every cluster versus all other clusters within the dataset. Genes with 1 or more UMI in at least 25% of cells within the two populations in the comparison (cluster-of-interest and all other clusters) were included in the analysis. Gene programs for each cluster included differentially expressed genes with log_2_fold-change [log_2_(FC)] of > 0.5 with a Bonferroni-adjusted *P* value of < 0.05. Gene programs can be found in Supplementary Table 2.

### Flow cytometry validation of enriched populations within the endosteal compartment

Flow cytometric analysis was used to investigate the enrichment of cell populations within the endosteal compartment relative to the bone marrow marrow. 9-week-old male C57BL/6J mice were sacrificed via CO_2_ asphyxiation and both femora and tibiae were collected. Soft tissue and the epiphyses were removed and marrow collected by flushing with PBS. Marrow-depleted bones were then crushed gently. To liberate adherent cells, marrow cells and bone fragments were subjected to enzymatic digestion with 2mg/ml collagenase A, 2mg/ml dispase and 0.1mg/ml DNaseI for 30 mins at 37°C. Following digestion, cells were filtered through a 100μm filter into collection tubes containing FCS (Bovogen Biologicals). Cells were collected via centrifugation at 400x g for 5 mins. Red blood cell lysis was then performed (Roche Diagnostics) and cells again collected by centrifugation. Cells were counted and incubated with Fc block (Biolegend; 101302) for 5 mins on ice. Cells were then stained with Zombie NIR Viability Stain (Biolegend; 423106), B220-BV510 (Biolegend; 103248), TCRβ-BV510 (Biolegend; 118131), CD45-BV650 (Biolegend; 103151), CD11b-BUV395 (BD Bioscience; 565976), Ly6C-BUV737 (BD Bioscience; 755201), Ly6G-APC (BD Bioscience; 560599) and CD14-PE (BD Bioscience; 569968) for a further 25 mins on ice. Cells were washed in PBS supplemented with 2% FCS. Samples were acquired using a BD FACSymphony machine and analysed with FlowJo v10.10.0 (BD Life Sciences).

### Defining genes with cell type-restricted expression

The restrictedness of gene expression was calculated by determining the number of clusters expressing a given gene. A cluster was deemed to express a gene if 1 or more UMI for that gene was detected in at least 25% cells within the cluster.

### Cell-cell interaction analysis

Putative intercellular interactions between clusters were identified via ligand-receptor analyses using CellPhoneDB v5.0.1^66^. Following Seurat clustering analysis, gene symbols were converted to HGNC symbols using the biomaRt package (version 2.64.0) ^67^. CellPhoneDB analysis was then performed using the “statistical_analysis_method” with default settings. A complete table of results is available in Supplementary Table 3. Within this process, all interactions are ascribed a directionality indicating which cell type is the “sender” cell and which cell type is the “receiver cell”.

To identify meaningful signalling pathways and processes associated with the identified ligands and receptors, we interrogated the ReactomeDB database of biological processes ^68^ for enriched terms using the “enrichPathway” function in the ReactomePA package (version 1.52.0) ^69^. All ligands and receptors involved in interactions between a given pair of cell types were used as input, with all genes present in the sequencing dataset used as the background universe. Enriched terms were annotated according to their root term within the ReactomeDB database and the level that they appear within that classification (root term = level 1). To minimise overlapping and redundant terms, Jaccard similarity values were calculated between each pair of terms based on the genes contributing to each term. Terms with a Jaccard similarity score of 0.8 and above were considered redundant and collapsed into a single term, with the term at level 3 selected for visualisation. Data were visualised using the ggplot2 (version 4.0.0) and viridis (version 0.6.5) R packages^70^.

### Reconstruction of cell differentiation trajectories

To infer the progression of cells across multiple differentiation stages and order them in pseudo-time, the algorithms implemented in the Monocle package (version 2) were used^71^. The top 1000 significantly differentially expressed genes were selected as the ordering genes for the trajectory reconstruction, using the nonlinear reconstruction algorithm DDRTree.

### Comparison between mouse and human scRNA-seq datasets

To compare scRNA-seq datasets we performed label transfer analysis with Seurat (version 5.3.0) ^72^, using the annotated mouse scRNA-seq dataset as the reference and the human scRNA-seq dataset as the query. A shared set of 3000 variable genes was identified using SelectIntegrationFeatures across both datasets. Transfer anchors were computed using FindTransferAnchors with the first 20 principal components. Cell-type annotations from the mouse reference were then transferred to the query datasets using TransferData, and prediction scores were added to the metadata. The prediction scores are derived from a softmax-like normalisation of label transfer anchors and reflect the maximum probability assigned to a transferred label for a given query cell. A prediction score of **>** 0.5 identifies high-confidence cell type predictions^72^.

### Identification of transcription factors regulating gene programs in cell clusters

To identify the transcription factors and their downstream target genes (regulons), the package pySCENIC (version 0.11.2) was used^73^. Gene regulatory network inference was performed with GRNBoost2, co-expression modules and potential direct targets of transcription factors were identified and then filtered using cisTarget. Regulon activity on single cells was scored using AUCell. The analysis was performed according to the standard SCENIC workflow.

### Single-nucleus ATAC-seq (snATAC-seq)

Endosteal cells were isolated from femora and tibiae of 9-week-old male *C57BL/6J* mice as described above. Single-cell suspensions were washed with PBS containing 1% BSA and pelleted (500 × g, 5 min, 4 °C). Cells were lysed in 100 µL custom lysis buffer (0.0075% digitonin, 0.05 mg/mL protease inhibitor [Millipore Sigma, 114298668001], 20 mM Tris-HCl [pH 7.4], 150 mM NaCl, 3 mM MgCl□, 2% BSA) on ice for 1–2 min with gentle pipetting to facilitate lysis. Nuclei were washed with washing buffer (0.025 mg/mL protease inhibitor, 20 mM Tris-HCl [pH 7.4], 150 mM NaCl, 3 mM MgCl□, 1% BSA) and pelleted (300 × g, 8 min, 4°C). Single-nucleus ATAC-seq (snATAC-seq) was performed using the 10x Genomics Chromium Single Cell ATAC v2 platform, following the manufacturer’s protocol. Briefly, nuclei were counted using a LUNA counter and haemocytometer to assess concentration and nuclear integrity. Nuclei were diluted to the recommended loading concentration and transposed using the Tn5 transposase in the ATAC transposition mix. Following transposition, gel bead-in-emulsions (GEMs) were generated using the Chromium Controller, enabling barcoding of accessible chromatin fragments in individual nuclei. Post-GEM amplification and library construction were carried out according to the 10x Genomics protocol. Final libraries were quantified using Qubit and assessed for fragment distribution using the Agilent Bioanalyzer or Tapestation. Sequencing was performed on an Illumina NovaSeq 6000 with paired-end 50□bp reads, targeting a depth of ∼25,000–50,000 fragments per nucleus. snATAC-seq services were provided by the Garvan Genomics Platform at the Garvan Institute of Medical Research.

Raw snATAC-seq sequencing data were processed using Cell Ranger ATAC (v2.1.0, 10x Genomics) for demultiplexing, alignment to the reference genome, barcode filtering, and peak calling. A median of 12,108 high-quality fragments per cell were detected, with 82.3% of fragments demonstrating high-quality overlap with peaks. The resulting fragment files were imported into R (v4.5.1) and analysed using the Signac (v1.15.0) and Seurat (v5.3.0) packages. Low-quality nuclei were filtered based on nucleosome signal, TSS enrichment score, and fragment count thresholds to remove likely debris or doublets. Peaks were quantified using the FeatureMatrix function, and term frequency–inverse document frequency (TF-IDF) normalization was applied, followed by dimensionality reduction using singular value decomposition (SVD) on the top components. Clustering was performed using the shared nearest neighbour (SNN) graph approach, and UMAP was used for two-dimensional visualization. Gene activity scores were computed by aggregating chromatin accessibility in gene bodies and promoter regions, and integrated with matched scRNA-seq data using label transfer via canonical correlation analysis (CCA), when applicable. Differential accessibility analysis between clusters or conditions was performed using logistic regression implemented in FindMarkers with fragment counts as covariates. Enriched transcription factor binding motifs within differentially accessible peak regions were identified using the FindMotifs function with default settings.

### Identification of genes within the gene programs that affect the skeleton

Genes from gene programs associated with biological processes important in the skeleton were identified using a curated list of gene ontology (GO) biological processes^74^ directly related to the skeleton^60^. Briefly, this list was constructed by filtering GO term descriptions using bone-related keywords. Genes associated with any of the 116 manually curated skeletal biological processes were then identified. This resulted in a final list of 663 genes. This list may be found in Supplementary Note 4.

Similarly, genes that cause a significant skeletal phenotype when mutated in mice were identified from the Mouse Genome Informatics (MGI) database^75^. Mouse gene ids (mgi_ids) with “abnormal skeletal phenotypes” were extracted using the mouse phenotype identifier MP:0005390 (date of access: 22^nd^ September 2023). Genes with alternative gene symbols or typographical errors were manually corrected and the list filtered for protein coding genes. This resulted in a final list of 2811 genes. This list may be found in Supplementary Note 4.

### Hypergeometric over-representation testing

To determine whether gene programs or lists were enriched for a specific subset of genes, over-representation analyses were performed using Fisher’s Exact Test under the hypergeometric distribution. Briefly, this test determines whether the number of genes of interest that are present within a given list of genes (successes in sample) is higher than would be expected by chance based on the total number of genes of interest within the background population (successes in population). For each test, a *P* value was calculated corresponding to the strength of evidence to reject the null hypothesis of no enrichment and adjusted for multiple testing via the Bonferroni correction method.

Parameters and details of *P* value adjustments for each over-representation test are outlined in Supplementary Note 5. For consistency, human Ensembl IDs were used as inputs for each test, using the biomaRt R package (version 2.64.0) ^76^ to convert from other gene identifiers where appropriate. All tests were performed using the RITAN package (version 1.24.0) in R^77^. Results are visualised with bar plots, bubble plots and FeaturePlots generated using the ggplot2 and Seurat R packages. Dark blue shades indicate significant enrichments (*P* ≤ 0.05 after Bonferroni correction); light blue shades indicate nominally significant enrichments (*P* ≤ 0.05 before Bonferroni correction and *P* ≥ 0.05 after Bonferroni correction).

#### Identifying gene programs enriched with monogenic skeletal disorder genes

To investigate whether skeletal disorder-causing genes were enriched among gene programs of bone and marrow cells, the current International Skeletal Dysplasia Society (ISDS) Nosology and Classification of Genetic Skeletal Dysplasias database was used^22^. This database encompasses 771 rare monogenic skeletal disorders categorised into 41 disorder groups (based on shared clinical, radiographic and molecular phenotypes), with pathogenic variants in 552 protein-coding genes or chromosomal aberrations attributed as causative. Genes with alternative gene symbols or typographical errors were manually corrected. The list was filtered for protein coding genes and genes that mapped to corresponding mouse orthologs using the biomaRt package in R (version 2.64.0) ^76^. This resulted in a final list of 528 genes which can be found in Supplementary Note 4.

Hypergeometric tests were performed and visualised as described above. Network plots were generated to allow for visual interpretation of enrichment analysis by depicting linkage between cells and the genes they expressed. Network plots were visualised using Cytoscape software (version 3.10.0) ^78^.

#### Quality control of ultrasound derived bone mineral density in the UK-Biobank Study

Between 2006 and 2010, the UK Biobank Study recruited 502,647 individuals aged between 37 and 76 years (99.5% were 40-69 years) located across the UK. The Northwest Multi-Centre Research Ethics Committee approved the UK Biobank Study, and informed consent was obtained from all participants. Quantitative ultrasound (QUS) assessment of calcanei was conducted on UK Biobank Study participants using a Sahara Clinical Bone Sonometer [Hologic Corporation (Bedford, Massachusetts, USA)]. Details of the complete QUS protocol is publicly available on the UK Biobank data showcase. Estimated heel bone mineral density [eBMD, (g/cm2)] was derived as a linear combination of two QUS parameters: speed of sound [SOS, (meters/second)] and bone ultrasound attenuation [BUA, (decibels/megahertz)] using methods described in Morris *et al*^8^. A total of 481,380 valid measures for eBMD passed QC (264,757 females and 216,623 males).

#### Identifying study participants with European ancestry

UK Biobank Study participants with high quality genotyping data (N=486,445) were projected onto the first 20 ancestry informative principal components (PCs) estimated from 1000 Genomes Project Phase 3 (1KG) individuals using GCTA version 1.93.2. PCA was based on a curated set of 38,512 LD-pruned HapMap3 (HM3) [20] bi-allelic SNPs that were shared between the 1KG and UK Biobank genotyped datasets [i.e. Minor allele frequency (MAF) > 1%, minor allele count (MAC) > 5, genotyping call rate > 95%, Hardy-Weinberg *P* > 1×10^−6^, and 13 regions with excessive linkage disequilibrium excluded]. The first 20 ancestry informative principal components of UK Biobank and 1KG individuals were projected into 3 components using Uniform Manifold Approximation and Projection (UMAP) using the following parameters: min_dist=0.000001, n_components=3, n_neighbors=35, random_state=10293082. Parameters were defined through supervised clustering by iterating over different parameter combinations and selecting parameters that clustered 1KG individuals with the same ancestry. UK Biobank Study participants that clustered with 1KG European ancestry individuals were identified by visual inspection. These individuals were then mapped back to PCA plots to ensure they co-localised with 1KG European ancestry individuals. Based on this analysis a total of 461,920 UK Biobank participants were deemed to have European ancestry.

### Genome-wide association analysis

A maximum of 448,010 European individuals (245,157 females and 202,853 males) with genotype and valid eBMD measures were available for analysis. As compared with a recent UK Biobank eBMD GWAS conducted by Morris and colleagues^8^, our study included 21,186 more individuals for the analysis of autosomal variants, and 85,085 more individuals for analysis of the sex chromosomes. The larger sample size was attributable to including individuals with European ancestry, as opposed to British ancestry analysed by Morris. Our study also included related individuals in the analysis of the X-chromosome, where Morris only analysed unrelated individuals due to software limitations. We also analysed the 3^rd^ version of UK Biobank imputed genetic data that had ∼10M more autosomal genetic variants and ∼730k more X chromosome variants than version 2 analysed by Morris.

Autosomal genetic variants and variants on the X, and X/Y chromosomes were tested for associations with eBMD assuming an additive allelic effect. A linear mixed non-infinitesimal model implemented in BOLT-LMM v2.3.4 was used to account for population structure and cryptic relatedness. Prior to GWAS eBMD had been transformed using a ranked based inverse normal transformation (mean=0, sd=1) and the following covariates were included as fixed effects in all models: age, sex, genotyping array, and ancestry informative principal components 1 to 20. Analysis was restricted to 21,684,109 high-quality imputed variants on the autosomal chromosomes, and 753,292 high-quality imputed variants on X-chromosome with a MAF > 0.05%, and imputation info score > 0.3.A Manhattan plot summarising genome-wide associations was generated using ggplot2 library within R^70^.

The genomic inflation factor (λ_GC_) was used to quantify the false positive rate and extent to which GWAS test statistics were inflated due to latent confounding. λ_GC_ was calculated as a ratio of the median of the empirically observed distribution of chi^2^ test statistics to the expected median (i.e., 0.456). The contribution of population stratification and other potential latent sources of bias to genomic inflation was quantified using stratified linkage disequilibrium score regression (LDSR) implemented in LDSC (version v1.0.1) in conjunction with pre-computed LD-Scores from European individuals from the 1000 Genomes project, assuming the BaselineLD model. LDSR analysis was limited to HapMap3 variants that passed QC using default software settings, supplemented with the --chisq-max 5000 flag to prevent disproportionate numbers of with small *P* values being excluded as a result of the software not being able to interpret double-precision floating-point number formats. The LDSC attenuation ratio statistic (RPS) was modest [LDSC_RPS_=0.02 (0.0195)], suggesting that polygenicity was the main driver of the observed inflation of test statistic (i.e. Genomic inflation factor λ_GC_□=□2.27). The high SNP heritability of eBMD (h2_SNP_=0.39 se=0.027) and large sample size of the study resulted in the LDSC intercept exceeding 1 (LDSR_I_=1.08 se = 0.073).

SNPTracker (version 1.0) was used to update reference SNP cluster IDs (rsID) and base pair positions from Hg19 to GRCh38h using dbSNP release 151^79^. SnpEff 5.1d^80^ was used in conjunction with GRCh38.p14 databases to assess the putative impact of all genetic variants and classify them as either high, moderate, or low impact. High impact variants are assumed to have high (disruptive) impact in the protein, causing protein truncation, loss of function or triggering nonsense mediated decay. Moderate impact variants are non-disruptive, but likely to change protein effectiveness. Low impact variants are assumed to be mostly harmless or unlikely to change protein behaviour. All high impact variants were cross referenced using Variant Effect Predictor (VEP) to confirm that SNP alleles and predictions were consistent^81^.

#### Fine mapping associated loci

Statistically independent genetic variants robustly associated with eBMD were identified using approximate conditional and joint association analysis (COJO) ^82^ implemented in GCTA (version 1.94.1) ^83^. The genome-wide significance threshold derived previously by Morris and colleagues was applied (*P* < 6.6×10^-9^). Genetic variants with high collinearity (multiple regression R^2^ > 0.9) were ignored and those situated more than 10 mega bases (MB) away were assumed to be in complete linkage equilibrium. To model patterns of linkage disequilibrium (LD) between genetic variants in GWAS, a reference sample of ∼50,000 unrelated European ancestry individuals from the UK Biobank Study was used^83,84^. The LD reference sample of unrelated individuals was identified using KING (kinship coefficient <0.044) ^85^ implemented in PLINK (version 1.9) ^86^. The following variant level QC was applied to resulting dataset, and variants were excluded if they deviated from Hardy Weinberg Proportions (*P* < 1×10^-6^), had a missing call rate less than 95%, were rare [(MAF) < 0.01%, and had a minor allele count (MAC) > 1]. The proportion of phenotypic variance explained by statistically independent association signals was calculated by summing the variance explained by primary and secondary signals using the formula described in Morris et al^8^:

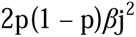

where p is the effect allele frequency, and *β*j is the joint and conditional effect of the allele on a standardized phenotype (mean□=□0, variance□=□1).

#### Annotation of independent association signals

Statistically independent variants reaching genome-wide significance (GWS, 6.6×10^-9^) in both marginal as well as conditional and joint analyses of eBMD were referred to as primary association signals, whereas those reaching GWS in conditional and joint analysis of BMD, but not marginal analysis were referred to as secondary association signals. Association signals were binned to ‘loci’ using a 1 MB sliding window. A locus was declared “novel” if it was located > 1 MB from any SNP reported to be associated with “bone density” in the NHGRI-EBI GWAS Catalog. For this annotation a reference list of BMD-associated SNP RSIDs meeting genome-wide significance (*P* < 5×10^-8^) was downloaded from the GWAS Catalog on 05/03/2024 using the term “EFO_000392”. The corresponding Hg38 co-ordinates were obtained using SnpTracker as previously described, and bedtools (version 2.29.2) ^87^ was used to estimate the minimum distances between each lead variant and the list of BMD associated variants. Lead association signals were also annotated to the closest protein coding gene using bedtools in conjunction with the Ensemble Genes 105 dataset that was downloaded from BioMart and based on genome-build GRCh38.p14^76^.

The closest protein coding gene to each lead variant were then followed up in the ISDS 2023 Nosology and Classification of Genetic Skeletal Diseases and further annotated if they caused monogenic skeletal disorders in humans when mutated^22^.

#### MAGMA gene-based tests of association

Gene-based tests of association were conducted using MAGMA software (version 1.10) in conjunction with GWAS summary statistics of variants with imputation quality score > 0.6 and MAF > 0.5%. Patterns of linkage disequilibrium (LD) were modelled with a reference sample of ∼50,000 unrelated European ancestry individuals from the UK Biobank Study. Genetic variants were annotated to protein coding genes based on Ensembl gene version 105 GRCh38. Variants were annotated to a gene if they were mapped ≤ 2kb upstream of the transcriptional start site and ≤ 1kb downstream of the stop site. A multi-model approach that combined the summary association results from 2 gene analysis models was used to derive an aggregate *P* value (*P*_Multi_) corresponding to the overall strength of evidence of association between each protein coding gene and eBMD. This multi-model approach was chosen as it yielded a more even distribution of statistical power and sensitivity over a wider range of different genetic architectures. The two gene-analysis models included in the multi-model were the ‘SNP-wise mean’ model that calculates the mean of χ2 statistics for all genetic variants annotated to a gene, as well as the ‘SNP-wise top’ model that uses the calculates the χ2 statistic for the lead genetic variant annotated to a gene. The threshold to declare statistical significance was determined using Bonferroni correction as follows: error rate of one test / number of genes tested = 0.05 / 19,856 = *P* < 2.54×10^-6^.

#### Enrichment analysis involving eBMD associated genes

Hypergeometric tests were performed and visualised as described above to investigate whether rare skeletal disorder causing genes were over-represented among the set of protein coding genes located closest to independent association signals, and separately the set of genes associated with eBMD as defined by MAGMA gene-based tests of association (Supplementary Table 6). Further hypergeometric tests were then used to determine whether enrichment was attributable to genes that caused individual skeletal dysplasia groups. The ISDS Nosology database was processed as outlined above with one alteration: the list was not filtered for genes with mouse orthologs. This resulted in a final list of 533 genes.

#### Identification of gene programs enriched with eBMD-associated genes

Competitive gene set analysis (GSA) was used to determine whether the gene-programs of different bone and marrow cells were enriched with eBMD associated genes. This was achieved by testing whether the set of genes making up the gene-program of a cell type was on average more strongly associated with eBMD, as compared with all other protein coding genes that were not in the gene-program under investigation. GSA accounted for several confounding factors including gene size, gene density and the inverse of the mean minor allele count in the gene, as well the log value of these three factors. Sensitivity analyses were also conducted in which adjustment was made for a set of protein coding genes that could be mapped between mouse and humans. This conditional analysis adjusted the marginal GSA results for the “mapped genes vs all human gene effect” that could potentially bias the analysis. The threshold to declare statistical significance was determined using Bonferroni correction as follows: Error rate of a single test / number of gene programs tested = 0.05 / 34 = *P* < 1.5×10^-3^.

GSA was also conducted on gene programs of the 16 non-haematopoietic cell sub-clusters - and using Bonferroni-corrected significance threshold of 0.05 / 16 = *P* < 3.1×10^-3^ was used to identify enriched cell subtypes. Post-hoc permutation analysis was conducted using R-scripts supplied with the MAGMA software. Exhaustive pairwise conditional analysis was also conducted on all enriched cell types to account for genes that were shared between gene programs of enriched cell types and cell states. A conditional *P* value < 0.05 was used to reject the null-hypothesis no-enrichment after correcting for shared genes.

#### Pulse rate as a negative control to for our gene enrichment workflow

We hypothesised that the cellular and genetic determinants of pulse rate were unlikely to be shared with those that regulate BMD. After conducting the analysis of eBMD, pulse rate was selected as a negative control for our study. We analysed pulse rate in the same set of individuals from the UK Biobank Study using the same workflow described above and compared results of enrichment analyses involving monogenic skeletal disorder genes, as well as MAGMA.

#### Identification of gene programs enriched with genes that cause abnormal bone structure when mutated in mice

Hypergeometric tests were performed and visualised as described above to investigate whether genes that regulate bone structural integrity were over-represented among gene programs of cell types and non-haematopoietic sub-clusters that were enriched with genes that cause monogenic skeletal disorders with abnormal bone structural integrity, and/or enriched with genes that were associated with eBMD (See Methods, Supplementary Note 4). Genes that cause a significant bone structural phenotype when mutated in mice were identified from the Mouse Genome Informatics (MGI) database^75^. Mouse gene ids (mgi_ids) with “abnormal bone structure” phenotypes were extracted using the mouse phenotype identifier MP:0003795 (date of access: 21^st^ May 2024). Genes with alternative gene symbols or typographical errors were manually corrected. The list was filtered for protein coding genes and genes that mapped to corresponding human orthologs using the biomaRt package in R^76^. This resulted in a final list of 1347 genes. This list may be found in Supplementary Note 4.

To determine whether genes that were more highly expressed in each gene program were more strongly enriched with genes that regulate bone structural integrity, we allocated genes to 6 nested groups based on the magnitude of fold-change in gene expression and repeated the gene-set enrichment analysis using all genes from each gene program, and then using only eBMD- associated genes from each gene program. The same methodology was applied to determine if gene programs of non-haematopoietic cell types were enriched for genes that regulate bone structural integrity.

### Non-Osseous Tissue Gene Expression Analysis

To interrogate expression of genes across tissues in mice, the TabulaMuris database was used^88^. Processed scRNA-Seq data was downloaded for each of the SMARTSeq FACS-based datasets from FigShare (https://figshare.com/articles/dataset/Single-cell_RNA-seq_data_from_Smart-seq2_sequencing_of_FACS_sorted_cells/5715040).

To interrogate expression of genes across tissues in human, the TabulaSapiens database was used^89^. Processed scRNA-Seq data were downloaded from FigShare (https://figshare.com/articles/dataset/Tabula_Sapiens_v2/27921984) and converted to Seurat objects using the “convertFormat” function from the sceasy package in R (version 0.0.7) ^90^. Datasets were filtered to include only cells analysed via SMARTSeq.

Each tissue was analysed individually. Cell clusters with <10 cells were excluded from analysis. Remaining clusters were manually assigned to a cell group (e.g. epithelial, mesenchymal, immune etc) using the Human Protein Atlas classification system (https://www.proteinatlas.org/humanproteome/single+cell+type) ^91^. The mean raw expression of selected exemplar genes within each cluster was then calculated and visualised as dotplots using the ggplot2 R package (version 4.0.0) ^92^. Mean expression values and cluster cell group assignments can be found in Supplementary Table 9.

### Genetically modified mice

Animal experiments were performed and reported in accordance with ARRIVE guidelines ^93^. All mouse skeletal phenotyping studies were undertaken under licence at Imperial College (project licence PPL70/8785 and PP1540664) and the Wellcome Trust Sanger Institute Mouse Genetics Project as part of the International Mouse Phenotyping Consortium and licenced by the UK Home Office (PPLs 80/2485 and P77453634) in accordance with the 1986 Animals (Scientific Procedures) Act and the recommendations of the Weatherall report. Animal experiments were approved by the Sanger or Imperial College Hammersmith Campus Animal Welfare Ethical Review Bodies (AWERB) as appropriate.

### *Pls3* knockout mouse line

C57BL/6N mice carrying a *Pls3^tm1a(EUCOMM)Wtsi^* knockout first allele (MGI:104807) were obtained from the Wellcome Trust Sanger Institute EMMA mouse repository and re-derived from frozen embryos. Genotyping was performed according to supplier’s protocols. The *Tm1a* knockout allele contains an internal ribosome entry site (IRES)-lacZ cassette in intron 3. The skeletal phenotype was studied in hemizygous male (*Pls3^y/-^*) mice and compared to WT (*Pls3^y+^*) male littermates. Following sacrifice, skeletal tissues were placed immediately into 70% ethanol or fixed in 10% neutral buffered formalin for 24 hours prior to storage at 4°C in 70% ethanol until analysis. All studies were approved by the Imperial College London AWERB and performed in accordance with the UK Animal (Scientific Procedures) Act 1986, the ARRIVE guidelines and EU Directive 2010/63/EU. All analyses were performed blind to sample genotype identification.

### Wild Type and mutant mice generated by the International Mouse Phenotyping Consortium

Samples from 16-week old wild-type (WT) and genetically-modified mice designed with deletion alleles on the C57BL/6Brd-*Tyr^c-Brd^*, C57BL/6Dnk, and C57BL/6N backgrounds were generated as part of the Wellcome Trust Sanger Institute’s (WTSI) Mouse Genetics Project (MGP), part of the International Mouse Phenotyping Consortium (IMPC; http://www.mousephenotype.org). Details on back-crossing status, weight, health status, administered drug and procedures, husbandry and specific conditions (including housing, food, temperature and cage conditions) have been reported previously ^94,95^. All mice generated by the WTSI MGP underwent a broad primary phenotype screen using consortium-wide protocols available at the IMPC portal (www.mousephenotype.org/impress). This includes body length, x-ray skeletal survey and biochemical measures of mineral metabolism performed between 14-16 weeks of age.

### Origins of Bone and Cartilage Disease bone phenotyping pipeline

Female mice from all WTSI MGP lines included in this study were sacrificed at 16 weeks of age, with the lower limb and tail fixed in 70% ethanol and subsequently phenotyped by the Origins of Bone and Cartilage Disease programme (OBCD) at Imperial College London. All samples were anonymised and randomly assigned to batches of 75 samples for rapid throughput analysis in a blinded and unselected fashion. The mouse line and genotype of each sample was only unblinded once all analyses had been completed for a batch. At least 5 contemporaneous WT samples were included in each batch to allow the identification of any systematic batch errors. The OBCD skeletal phenotyping programme determined 19 parameters of bone mass and strength^7,8,43^. The key phenotype definitions are as follow:

- Structural phenotypes – Significant difference in bone structural parameters measured by digital X-ray microradiography or micro-computerised tomography.
- Functional phenotypes –Significant differences in mechanical strength parameters measured by biomechanical testing
- Structural and functional phenotypes – Significant differences in at least one structural and one functional parameter
- Bone quality phenotypes – lines with outlier functional phenotypes that did not correlate with bone mineral content
- Mahalanobis phenotypes – lines that were significant outliers due to smaller differences in multiple skeletal parameters.

### Digital x-ray microradiography

Soft tissue was removed from the fixed bones and digital X-ray images recorded at a 10μm resolution using a Faxitron MX20 variable kV point projection x-ray source and digital imaging system (Qados, Cross Technologies plc, Sandhurst, Berkshire, UK) operating at 26kV, 15s, and 5x magnification. The magnification was calibrated by imaging a digital micrometer. For each sample the cleaned lower limb and caudal vertebrae Ca6 and Ca7 were imaged in frame with 3 standards; a 1mm diameter steel wire, a 1mm diameter spectrographically pure aluminium wire, and a 1mm diameter polyester fibre. Relative bone mineral content, a two dimensional parameter similar to areal bone mineral density, and bone lengths were determined as previously described^96^. Briefly, 2368×2340 16-bit DICOM images were converted to 8-bit Tiff images in ImageJ, the grey levels for the polyester and steel standards were used to stretch each image across the 256 grey levels with polyester at grey level 0 and steel at grey level 255 using macros described in^97^. Increasing gradations of mineralisation density were represented in 16 equal intervals by applying a pseudocolour lookup table to each image. For each sample the median grey level (0-255) of the femur and caudal vertebrae 6 and 7 was calculated by determining the number of pixels of each grey level in the stretched X-ray image. Lengths were determined using ImageJ 1.44 software. The results for caudal vertebrae 6 and 7 were averaged. For male *Pls3^y/-^* mice, cleaned lower limb, caudal vertebrae Ca6 and Ca7, and lumbar vertebrae L5 were imaged at 10µm pixel resolution using a Faxitron UltraFocus digital radiography system (Faxitron Bioptics LLC, Arizona, USA).

### Micro-computerised tomography (µCT)

A Scanco µCT50 (Scanco medical, Zurich, Switzerland) was used to determine the three-dimensional cortical and trabecular structural parameters of femurs. Samples were scanned at 70kV, 200μA, with a 0.5mm aluminium filter, 1 second integration time, no averaging, and images captured every 0.36◦ though 180◦ rotation. The Scanco Medical software suite (µCT Tomography v6.4-2/Open VMS) was used for reconstructions, ROI selection and analyses. 10μm voxel resolution scans of a 1.5 mm long region of mid-shaft cortical bone were used to calculate cortical bone parameters (Cortical thickness Ct.Th, internal endosteal diameter ID, and Bone Mineral Density BMD). The ROI was centred at 56% along the length of the femur, distal to the femoral head, to provide a circular cross section avoiding the linea aspera. Trabecular parameters (Trabecular bone volume per tissue volume BV/TV, trabecular number Tb.N, Trabecular thickness Tb.Th, and trabecular spacing Tb.Sp) were calculated from 5μm voxel resolution scans with an ROI consisting of a 1mm long region of the trabecular compartment beginning 100μm proximal to the distal femoral growth plate. For male *Pls3^y/-^* mice femurs and lumbar vertebrae L5 were imaged. The entire volume of the body of the L5 vertebra was included in the trabecular analysis, with contouring used to exclude the thin layer of cortical bone. Trabecular bone BMD was not included because of the potential for partial volume artifacts due to the broad range of trabecular thickness relative to voxel resolution. Furthermore, cortical porosity (Ct.Po) and canal diameter (Ca.Dm) were determined in a 1 µm voxel resolution scan of a 500 µm ROI centred in the midshaft 56% along the length of the femur distal to the femoral head^60^.

Additional cortical vascular and cortical osteocyte lacuna analysis were performed in femur samples from P112 male mice. A 250 μm ROI centered in the midshaft, 56% along the length of the femur distal to the femoral head, was scanned at 1 μm voxel resolution and analysed using previously optimized segmentation thresholds^60^. DICOM images were processed using Fiji and lacunae rendered using its Volume Viewer (https://imagej.nih.gov/ij/plugins/volume-viewer.html) or Drishti-3.2. Vascular canals and osteocyte lacunae parameters were analyzed using BoneJ Particle Analyzer^98^. Particles with a volume >2000 µm^3^ were designated vascular channels and particles between 100 and 2000µm3 were designated osteocyte lacunae^99^. Vascular canal area (Va.Ca.V/BV) and number (VaA.Ca.N/BV) per bone volume, and mean vascular canal volume (Mean Va.Ca.V) in mutants were compared to WT controls. The distribution of vascular canal volumes was compared to WT by randomly selecting the maximum equivalent number vascular canals from each sample (90 per sample) and performing Kolmogorov-Smirnov analysis (GraphPad Prism version 10). Similarly cortical microporosity (Ct.µPo), osteocyte lacunae number per bone volume (Lc.N/BV), mean lacuna volume (Mean Lc.V) in mutants were compared to WT. The distribution of Lacuna volumes was compared to WT by randomly selecting the maximum equivalent number vascular canals from each sample (6275 per sample) and performing Kolmogorov-Smirnov analysis. Differences in volume distributions between mutant and WT were considered valid if significant in all 10 permutations performed.

### Biomechanical Testing

An Instron 5543 materials testing load frame (Instron Limited, High Wycombe, UK) was used to perform destructive 3-point bend tests and 2-point compression tests respectively on femurs and caudal vertebrae Ca6 and Ca7 (100N load cell for femurs and 500N load cell for vertebrae) (Instron Limited, High Wycombe, UK) as described^7,100^. Femurs were positioned horizontally with the anterior surface upwards between two custom mount points with rounded ends and a total span of 8mm. Individual vertebrae were bonded in vertical alignment to a custom anvil support using cyanoacrylate glue. Load was applied vertically to the mid-shaft of the femur, or evenly across the vertebral end plate with a constant rate of displacement of 0.03mm/second until femoral fracture, or ∼1 mm of displacement had occurred in the vertebra. Biomechanical properties were calculated by plotting load displacement curves for each biomechanical test and determining yield, maximum and, for femurs only, fracture loads. Stiffness was calculated from the slope of the linear portion of the load displacement curve using the “least squares” method. Percentage energy dissipated at fracture load was calculated, as previously described, by subtracting the elastic stored energy at fracture from the total work energy at fracture^43^. The results for caudal vertebrae 6 and 7 were averaged for each parameter. For male *Pls3^y/-^* mice, adjoining spinous processes were removed from lumbar vertebrae and the vertebral body bonded in vertical alignment to a custom anvil support using cyanoacrylate glue and load was applied vertically at a constant rate of displacement of 0.03 mm/s until approximately 0.5 mm of displacement had occurred. Vertebral yield load, maximum load, and stiffness were derived from load displacement curves.

### Whole-mount skeletal staining

P1 neonates were prepared and stained with Alizarin red (mineralised bone) and Alcian blue (cartilage) by standard techniques^101^. Whole mount preparations were imaged in glycerol using a Leica MZ75 binocular microscope, Leica KL1500 light source, and Leica DFC320 digital camera (Leica Microsystems Ltd).

### Growth plate histomorphometry

Tibias were fixed for 24 hours in 10% neutral buffered formalin, decalcified in 10% EDTA pH 7.4 and decalcification was confirmed by X-ray microradiography. Samples were embedded in paraffin and 5µm sections stained with Alcian blue (cartilage) and van Gieson (osteoid) ^102^. Images were obtained using a Leica DM LB2 microscope and DFC320 camera. The width of the proximal tibial growth plate reserve, proliferative, and hypertrophic zones were determined in at least four locations for each section using ImageJ (http://rsb.info.nih.gov/ij/).

### Osteoclast static histomorphometry

Proximal humeri were fixed for 24 hours in 10% neutral buffered formalin, decalcified in 10% EDTA pH 7.4 and decalcification was confirmed by X-ray microradiography. Samples were embedded in paraffin and 5µm sections stained for tartrate-resistant acid phosphatase (TRAP) (osteoclasts) and counter stained with aniline blue (bone) ^103^. Histomorphometry analysis was performed with TrapHisto software (https://www.liverpool.ac.uk/ageing-and-chronic-disease/bone-hist/trap-hist/) in a 750×750 µm area of measurement commencing 250 µm distal to the proximal humeral growth plate^103,104^.

### Osteoblast dynamic histomorphometry

Mice were double-labelled with 15 mg/kg calcein (Sigma C0875) by tail-vein injection of calcein (2.5 µg/mL phosphate-buffered saline) at six and two days prior to collection at P70. Femurs and lumbar vertebrae were embedded in methacrylate and midcoronal block faces were cut and polished to an optically flat surface. To quantify osteoblastic bone formation parameters fluorescent calcein labelling was imaged using confocal autofluorescence scanning light microscopy (CSLM). A Leica SP5 scanning confocal microscope at 488 nm excitation with x40/1.25 objective^105^. Montages of images containing trabecular and cortical endosteal bone surfaces were analysed using ImageJ to determine total bone surfaces, calcein-labelled surfaces and the separation between calcein double-labels according to the American Society for Bone and Mineral Research system of nomenclature^104^. Femoral trabecular bone was analysed in a 1.5×1.5 mm region of interest (ROI) commencing 500 µm proximal to the distal femoral growth plate, and cortical endosteal bone surfaces were analysed 2 mm below the growth plate for a length of 1 mm. Lumbar vertebral trabecular bone was analysed from the entire inner region of a vertebra body from region L1-3 excluding the peripheral cortical bone.

### CD31 Immunohistochemistry

Femurs, from P21 male mice, were fixed for 24 hours in 10% neutral buffered formalin, decalcified in 10% EDTA pH 7.4 and decalcification was confirmed by X-ray microradiography. Samples were embedded in paraffin and platelet endothelial cell adhesion molecule (PECAM-1, CD31) immunohistochemistry was performed using 5µm longitudinal midline sections. PECAM-1 antigen retrieval was performed in citrate buffer pH 6 at 60°C for 2h. The primary rabbit anti-CD31 antibody ab182981 (Abcam; Cambridge UK) was diluted 1:200. Visualization was performed using Leica PowerVision Poly-HRP anti-Rabbit IHC Detection System (Leica Biosystems PV6113) according to the manufacturer’s instructions. Mouse lung sections were used as a positive control tissue and no primary antibody controls were also included. Sections were counterstained with haematoxylin and imaged using a Leica DM LB2 microscope, Leica Flexacam C1 camera and LAS X software. Images were captured at a resolution of 2016 pixels/mm and full femur montages generated. A 2cm long mid femur cortical bone ROI was defined, and the two cortical ROI images from each sample randomized. ImageJ was calibrated and each cortical ROI image was analysis blind to determine, total cortical area, number of vascular canals, and area of each vascular canal. Individual images were subsequently unblinded and the mean values for the two cortical ROIs form each femur calculated and mutants compared to WT controls.

### Iodine Contrast Enhanced BSE-SEM (ICE-BSE-SEM)

Formalin fixed femurs, from P70 male mice, were embedded in polymethyl methacrylate (PMMA), polished to an optically flat finish, and stained with potassium iodide. Stock Lugol’s Iodine solution (Pro-Lab Diagnostics) was diluted with an equal volume of absolute ethanol and pipetted directly onto the block surface. After 15 min, blocks were washed with distilled water and air dried. Blocks were carbon coated and imaged by SEM (Tescan UK, Cambridge, UK) at high vacuum with a 4-quadrant back-scattered electron detector (Deben,UK) (Male, n=6 per genotype at P70). Images were captured at 20 kV, 0.4 nA and montages of each sample generated at a resolution of 819 pixels/mm. Some higher resolution images were also acquired at 2560 pixels/mm. A 3 cm long cortical bone ROI was defined starting 1cm above the distal femoral growth plate and the two cortical ROI images from each sample randomized. ImageJ was calibrated and each cortical ROI image was analysed blind to determine total cortical area, number of vascular canals, and area of vascular canals. Individual images were subsequently unblinded and the mean values for the two cortical ROIs from each femur calculated and compared to WT controls.

### Serum analysis

Serum was obtained by centrifugation of blood obtained by terminal cardiac puncture. Serum bone resorption (C-terminal telopeptides of type 1 collagen (CTX)) and bone formation (N-terminal propeptide of type I procollagen (P1NP)) were determined by ELISA at P70 (Immunodiagnostic Systems Ltd, Boldon, Tyne & Wear, UK). The inhibitors of Wnt-mediated osteoblastic bone formation Dickkopf related protein-1 (DKK-1) and Sclerostin (SOST) were determined by ELISA at P183 (R&D Systems Europe Ltd, Abingdon, Oxfordshire, UK).

### OBCD bone phenotyping pipeline quantification and statistical analysis

#### Determination of Wild Type Reference Ranges

To determine if the KO lines had an abnormal skeletal phenotype, skeletal parameters were compared to those of one of two WT cohorts depending on their genetic background. 918 KO lines were compared to the reference range of the primary WT cohort composed of 320 C57BL/6N and C57BL/6NTac mice. 132 lines were evaluated against the reference range of a secondary WT cohort composed of 80 C57BL/6Brd-Tyr^c-Brd^ and C57BL/6Dnk mice. Frequency distribution of datasets was assessed by D’Agostino and Pearson test. For the primary WT cohort, the normally distributed parameters were: Femoral Bone Mineral Content, Trabecular Bone Volume per Tissue Volume, Trabecular Number, Cortical Thickness, Cortical Internal Diameter, Femur Yield Load, Maximum Load, and Stiffness, Vertebral Bone Mineral Content, Length, Yield Load, Maximum Load and Stiffness. Non-normally distributed parameters were: Femur Length, Trabecular Thickness, Trabecular Spacing, Cortical Bone Mineral Density, Femur Fracture Load and Toughness (Energy dissipated at fracture). For the secondary WT cohort, the normally distributed parameters were: Femoral Bone Mineral Content, Length, Trabecular Number, Trabecular Thickness Cortical Internal Diameter, Femur Yield Load, Maximum Load, Fracture Load and Stiffness, Vertebral Bone Mineral Content, Length, Yield Load, Maximum Load and Stiffness. Non-normally distributed parameters were: Femur Trabecular Bone Volume per Tissue Volume, Trabecular Spacing, Cortical Thickness, Cortical Bone Mineral Density, and Femur Toughness (Energy dissipated at fracture). Comparisons of WT primary and secondary cohort values with unpaired Student’s *t*-test or Kolmogorov-Smirnov tests demonstrated statistically significant differences among the two genetic background strains. All mice analysed during this study are females of age 15.3-16.7 weeks. Statistical analyses and plots for the WT cohorts have been performed using GraphPad Prism 9.

#### Reference Range Analysis and Permutation Testing of Outliers

The Reference Range Analysis involved two distinct approaches to identify KO lines with outlier skeletal phenotypes. Firstly, a KO line was considered as an outlier if the average value of any of the 19 parameters was outside of the WT reference range defined as 2 standard deviations above or below the mean (normally distributed parameters), or outside the 2.5–97.5th percentiles (non-normally distributed parameters). Secondly, for each parameter of each KO line we used permutation testing (100,000 permutations using Julia (https://juliastats.org/) to determine the probability of selecting an equal number of WT samples, from the appropriate cohort, with a parameter value as or more divergent than that observed in the KO line from the WT mean/median. The *P*-value was determined by dividing the number times the WT sample’s parameter was as or more divergent than that of the KO line by 100,000. Significance thresholds were calculated by applying a Bonferroni correction for the number of effective tests to a 5×10^-2^ significance threshold. We determined the number of effective tests by obtaining the eigenvalues of the correlation matrix of the 19 skeletal parameters of the WT baselines using the eigen function in Julia. We then estimated the number of effective tests (Neff) as 11.4 for the primary genetic background and 10.7 for the secondary genetic background using the formula:

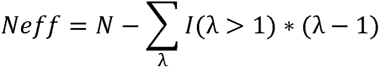

Where N = 19 as the number of the skeletal parameters and λ indicates the eigenvalues. The outlier *P*-value significance threshold was 4.37×10^-3^ for the KO lines on the primary genetic background and 4.69×10^-3^ for the KO lines on the secondary genetic background.

#### Mahalanobis distance outliers

To ensure that significant abnormal phenotypes resulting from simultaneous but smaller variances in any of the 19 parameters were not overlooked, we performed a multivariate analysis defining the Squared Robust Mahalanobis distances (MDi2) for each skeletal sample. Only samples with values for all the 19 phenotype parameters were evaluated. The MD¬i2 were computed using the heplots R package (https://friendly.github.io/heplots/reference/Mahalanobis.html) with the minimum volume ellipsoid method for the robust estimate of the centre. We tested for multivariate normality by performing a Mardia’s test combining kurtosis and skewness tests (MVN R package) and by visually inspecting the Chi-Square Q-Q plots. Under the assumption of multivariate normality, the distribution of MD¬i2 is approximately chi-squared with p degrees of freedom, with p determined by the number of variables (19). A sample was designated outlier if its MDi2 > χ2p; 0.975 (i.e. MDi2 > 32.852, determined in Microsoft Excel with formula CHISQ.INV(0.975,19)). A KO line was designated a Mahalanobis Outlier if >50% of the KO lines’ samples were outliers by Mahalanobis analysis.

#### Bone quality outliers

Using the results from primary or secondary WT cohorts, linear regression analysis was performed comparing femur BMC to each femur strength parameters, and vertebra BMC to each vertebra strength parameters. Lines of best fit, R^2^, and *P-*values were calculated. 95% prediction intervals were determined for all parameter pairs where the linear regression *P-*values was <0.05. The results for KO lines were compared to the 95% prediction intervals. A KO line was designated a Bone Quality Outlier if the mean/median value of any of its parameter pairs were not contained within the WT 95% prediction interval for that parameter pair.

### Statistical analysis of male *Pls3^y/-^* samples

Data from hemizygous male *Pls3^y/-^* mice were compared to male WT littermates by an unpaired two-tailed Student’s *t*-test. *P*-values < 0.05 were considered statistically significant. The Kolmogorov-Smirnov test was used to compare cumulative frequency distributions of BMC^43,106^.

### Zebrafish husbandry

All zebrafish experimentation adhered to the guidelines of the animal ethics committee of the University of Queensland (Permit 2022/AE000091). Zebrafish transgenic line utilized in this study is *Tg(kdrl:EGFP)^s843^* ^107^. Embryos were obtained through natural paired matings and were incubated at 28°C in a dark-phase incubator. Embryos were maintained in 10 cm petri dishes containing 1X E3 media (5 mM NaCl, 0.17 mM KCl, 0.33 mM CaCl_2_, 0.33 mM MgSO_4_) at a maximum density of n=60. At 24 hpf, all embryos were changed to E3 media supplemented with 0.0003% phenylthiourea (PTU) to prevent pigmentation.

### Genome editing and genotyping of pls3 mutants in zebrafish

To generate G_0_ mosaic crispants, three crRNAs targeting *pls3* were designed using IDT Alt-R Custom Cas9 Design Tool. Sequences of the three crRNAs are as follows:

*pls3* crRNA1: 5’ – caagactataccatcaccta – 3’;
*pls3* crRNA2: 5’ – gagcaagtgaaagtaagccc – 3’;
*pls3* crRNA3: 5’ – acaaaggccaagttgagttt – 3’.

gRNAs were prepared according to the IDT synthesis protocol. In brief, 0.4 μL of each 100 μM crRNA was mixed with 1.2 μL of 100 μM tracrRNA and 0.95 μL of Nuclease-free Duplex Buffer. The mix was incubated at 95°C for 5 minutes and cooled down to room temperature for 3 minutes. Ribonucleoprotein injection mix was prepared with 0.45 μL Alt-R^TM^ S.p. Cas9 Nuclease V3 (100 μg, IDT, 1081058), 3 μL annealed gRNAs cocktail, 2.25 μL 1M KCl, 0.75 μL phenol red and 1.05 μL UltraPure Water, followed by incubating at 37°C for an hour. Zebrafish embryos were injected at the one-cell-stage with ∼1 nL of ribonucleoprotein injection mix. Both uninjected control and injected embryos were grown up to 72 hpf for live imaging.

Following live imaging, DNA was extracted from uninjected control and injected embryos, and genotyped to determine cutting efficiency of the gRNAs. Primers sequences for amplification of amplicons flanking crRNA target sites:

*pls3* 1F: 5’ – aaactgcccatgaaccctaa – 3’
*pls3* 1R: 5’ – tgaaaccatggacaatatgttaaa – 3’
*pls3* 2F: 5’ – aagctacattttcacttaaccttttt – 3’
*pls3* 2R: 5’ – tgacctttaggtgcaatctgg – 3’
*pls3* 3F: 5’ – tgcagacagtttgtcacgtct – 3’
*pls3* 3R: 5’ – tttcgggttttgtcagtgct – 3’

### Confocal imaging

Live zebrafish embryos were mounted laterally in 1% low-melting point agarose and imaged on a Zeiss LSM 710 confocal microscope using a PlnApo 20x/0.8 DICII objective. Image analysis was conducted using Image J. ISV luminal diameter was quantified by application of a Vasometrics macro^108^.

## Data availability

scRNA-seq raw data has been deposited in the GEO database and will be made available upon publication. Published human spatial transcriptomic data generated by Yip and Er *et al* ^52^ that was reanalysed is available on GEO: GSE299207. Genome-wide association summary result statistics will be made publicly available via the NHGRI-EBI GWAS Catalog upon publication. Source data for Figs. 1-7 and Extended Data Figs. 1-9 are included in the supplementary notes. All other data are available from the corresponding authors upon reasonable request.

## Code availability

scRNA-seq was analysed using the Seurat software (https://satijalab.org/seurat/). Code for constructing cell type clusters, trajectories and gene programs from scRNA-seq will be made available upon publication via GitHub (https://github.com/). Code used for genetic association analyses and gene set analyses will also be made available upon publication via GitHub.

## Acknowledgements

This work was supported by a Wellcome Trust Strategic Award (101123/Z/13/A) to G.R.W., P.I.C. and J.H.D.B. and by Mrs. Janice Gibson and the Ernest Heine Family Foundation. J.P.K. is funded by a National Health and Medical Research Council (Australia) Investigator Grant (GNT2026272) and Project Grant (GNT1158758); the Lions Medical Research Foundation (2020 Lions Dunning-Orlich Investigator Award), the American Society of Bone and Mineral Research (2020 ASBMR Rising Star Award) and the Mater Foundation. P.I.C and T.G.P are supported by NHMRC Investigator Grants (GNT2009010 and GNT1155678). P.I.C. is supported by a National Breast Cancer Foundation Collaborative Research Accelerator grant (2024/CRA0020). J.H.D.B and G.R.W are supported by a Wellcome Trust Joint Investigator Award (110141/Z/ 15/Z and 110140/Z/15/Z) and European Union H2020 THYRAGE Grant (666869). M.L. was supported by a University of Queensland Research Training Scholarship from The University of Queensland (UQ) and a Postgraduate Top-Up Scholarship from the Commonwealth Scientific and Industrial Research Organisation (CSIRO). J.T.S. was funded by a UNSW Sydney International Postgraduate Award. A.P.C. and S.E.Y. were funded by the Australian Government Research Training Program Scholarships. R.E.M. was supported by the Finnish ORL–HNS Foundation, the Orion Research Foundation and the Juhani Aho Foundation. D.M.E. is supported by a NHMRC Investigator Grant (GNT2017942). C.A.L. was supported by a European Research Council advanced grant (ERC-2018-ADG 834057). E.D.H is supported by a fellowship from the National Health and Medical Research Council (GNT2008652). D.P.K. were supported by a grant from the National Institute of Arthritis and Musculoskeletal and Skin Diseases (R01 AR041398). Additional support came from the UNSW Cellular Genomics Futures Institute, Cancer Council NSW and the Leukemia Research Foundation. This research has been conducted using the UK Biobank Resource under application number 53641. We thank Eric Lam, Sunny Wu, Chia-Ling Chan, Nona Farbehi, Yasmin Husaini, Zun Han Lu, Orion Tong, Alejandro Rios Villamil, Samira Samarfard, Chris O’Keefe and the staff of the Garvan Genomics Platform at the Garvan Institute of Medical Research for support with FACS, scRNA-seq and snATAC-seq; and the Australian BioResource Biological Testing Facility. This study was supported by phenotyping expertise provided by Anne-Tounsia Adoum, Rebecca Allen, Jayashree Bagchi-Chakraborty, Katharine F. Curry, Hannah F. Dewhurst, Apostolos Gogakos, Fiona Kussy, Naila Mannan, Justyna Miszkiewicz, Hayley J Protheroe, Penny C Sparkes and Valerie Vancollie. We also thank Benjamin Mullin, Shannon D’Urso, Laura Scott, Chrissy Hammond and David Hume for their intellectual contributions as well as Luke Hargreaves, Owen Powell, Leslie Elliott and the UQ and IMB ITS support teams for assistance with high performance computing.

## Author contributions

R.C.C., J.H.D.B., G.R.W., P.I.C. and J.P.K. conceived the study. R.C.C., J.H.D.B., G.R.W., P.I.C. and J.P.K. designed experiments. R.C.C., A.C-M., S.G.Y. and E.S. isolated cells from mouse and human bone for scRNA-seq and snATAC-seq. R.C.C., J.T.S., A.P.C. and W.H.K. performed analyses on the scRNA-seq data. M.L. and J.P.K. processed and analysed the GWAS data. R.C.C., M.L., J.T.S. and J.P.K. performed integrated analysis of scRNA-seq and GWAS. M.F., Y.Z., K.A.F. performed additional analyses on the GWAS data. B.F., D.K., M.R.G.D., S.E.G., J.G.L., N.C.B., V.D.L., A.S.P., R.E.M. performed skeletal phenotyping of mutant mice. A.P.B. and Y.Z. designed and implemented the interactive portal. Y.C., R.K.H.Y., J.E., E.D.H. generated and analysed spatial transcriptomics data. S.Z.T. and A.K.L. generated and performed vascular phenotyping of mutant zebrafish. N.B. and H.L.O recruited patients for human scRNA-seq. L.R.-M, A.V.-Z, H.L.O., R.M., J.P.T.-K., E.M.H., M.M.M., S.E.Y., C.M.S., D.M.E., J.E.P., C.A.L., A.D.E., J.A.E., P.A.B., R.D.B., T.G.P., E.L.D. and IFMRS Big Data Working Group provided expert guidance and feedback on experimental design, analysis and results. R.C.C., M.L., J.T.S., J.H.D.B., G.R.W., P.I.C. and J.P.K. wrote the manuscript. All authors reviewed and revised the manuscript.

## IFMRS Big Data Working Group

Cheryl L. Ackert-Bicknell^1^, Douglas P. Kiel^2,9,12^, Fernando Rivadeneira^3^, Jennifer J. Westendorf^4^, David Karasik^5^, Yuuki Imai^6,7^, Ralph Müller^8^, Jason Flannick^9,10^, Lynda Bonewald^11^, Noël P. Burtt^12^, Jonathan H. Tobias^13,14^, Carolina Medina-Gomez^3^, Qing Wu^15^, Maria C. Costanzo^12^ and Charles R. Farber^16^

1. Department of Orthopedics, University of Colorado Anschutz Medical Campus, Aurora, CO, USA
2. Hinda and Arthur Marcus Institute for Aging Research, Hebrew SeniorLife Department of Medicine, Beth Israel Deaconess Medical Center and Harvard Medical School, Boston, MA, USA
3. Department of Internal Medicine, Erasmus Medical Centre, Rotterdam, the Netherlands
4. Department of Orthopedic Surgery, Mayo Clinic, Rochester, MN, USA
5. The Musculoskeletal Genetics Laboratory, The Azrieli Faculty of Medicine, Bar-Ilan University, Safed, Israel
6. Department of Pathophysiology, Ehime University Graduate School of Medicine, Toon, Ehime, Japan
7. Division of Integrative Pathophysiology, Proteo-Science Center (PROS), Ehime University, Toon, Ehime, Japan
8. Institute for Biomechanics, Department of Health Sciences and Technology, ETH Zürich, Zürich, Switzerland
9. Harvard Medical School, Boston, MA, USA
10. Division of Genetics and Genomics at Boston Children’s Hospital, Boston, MA, USA
11. Indiana Center for Musculoskeletal Health, Indiana University, Indianapolis, IN, USA
12. Broad Institute of MIT and Harvard, Cambridge, MA, USA
13. MRC Integrative Epidemiology Unit, University of Bristol, Bristol, UK
14. Musculoskeletal Research Unit, Translational Health Sciences, Southmead Hospital, University of Bristol, Bristol, United Kingdom
15. Department of Biomedical Informatics, College of Medicine, The Ohio State University, Columbus, OH, USA
16. Department of Genome Sciences, University of Virginia, Charlottesville, VA, USA

---

**Extended Data Fig. 1. Genes and gene programs that define murine chondrocytes, endothelial cells, vascular smooth muscle cells and osteoclasts**

Analysis of gene programs that define the chondrocyte (cluster 27), endothelial cell (cluster 31), VSMC (cluster 32) and osteoclast (cluster 30) clusters from Fig. 1c.

(**a**) Density plots of log_2_fold-change [log_2_(FC)] in expression of genes (top) and total number of clusters expressing genes (bottom) that define the chondrocyte gene program (cluster 27).

(**b**) Waffle plots for individual genes (coloured squares) in the chondrocyte gene program with log_2_(FC) 0.5-1 (top), log_2_(FC) 1-2 (middle) and log_2_(FC) >2 (bottom) in expression. Genes annotated with a skeletal process in the GO database (green), MGI database (purple), in both databases (blue) or are unannotated (orange) are shown. Circos plot shows individual genes with a log_2_(FC) >2, with those found only in the chondrocyte cluster gene program indicated in bold.

(**c**) Density plots of log_2_(FC) in expression of genes (top) and total number of clusters expressing genes (bottom) that define the endothelial cell gene program (cluster 31).

(**d**) Waffle plots and circos plot for individual genes in the endothelial gene program. Annotated as described in (**b**).

(**e**) Density plots of log_2_(FC) in expression of genes (top) and total number of clusters expressing genes (bottom) that define the VSMC gene program (cluster 32).

(**f**) Waffle plots and circos plot for individual genes in the VSMC gene program. Annotated as described in (**b**).

(**g**) Density plots of log_2_(FC) in expression of genes (top) and total number of clusters expressing genes (bottom) that define the osteoclast gene program (cluster 30).

(**h**) Waffle plots and circos plot for individual genes in the osteoclast gene program. Annotated as described in (**b**).

**Extended Data Fig. 2. Cell-cell interaction of osteoblast lineage cells with other non-haematopoietic cells and osteoclasts**

(**a**) Circos plots displaying predicted interactions between osteoblast lineage cells and selected cell types. Genes are arranged according to the annotated pathways present within CellPhoneDB and labelled accordingly in the outer layer. Interaction types are indicated by coloured boxes. Line colour indicates which cell type is the “sender” cell type for a given interaction, whilst thickness reflects interaction specificity. For clarity, only interactions with a specificity score ≥50% are included within the plots.

(**b**) Bar plots displaying enriched ReactomeDB pathways identified amongst interactions between osteoblast lineage cells and selected cell types. Pathways are grouped according to their root term within the ReactomeDB database (see Methods). Adjoining numbers indicate the proportion of genes associated with a term that were present within the results. Terms that are uniquely enriched for one cell type are indicated in grey boxes and red outlines/text.

**Extended Data Fig. 3. High-resolution clustering of non-haematopoietic cells**

(**a**) UMAP of cell clusters identified in osteoblast lineage, chondrocyte, endothelial cell, VSMC and neuronal cell types.

(**b**) Heatmap of the top 10 genes expressed by each of the non-haematopoietic sub-clusters ranked by log_2_(FC). Two example genes that define each sub-cluster are annotated.

(**c**) Predicted relationship between individual osteoblast sub-clusters determined by pseudo-time trajectory analysis.

(**d**) Expression of exemplar genes of each osteoblast sub-cluster in pseudo-time.

(**e**) Gene ontology analysis of the gene programs for each osteoblast sub-cluster.

(**f**) Bar plot showing the proportion of genes in osteoblast sub-cluster gene programs. Genes annotated with a skeletal process in the GO database (green), MGI database (purple), in both databases (blue) or are unannotated (orange) are shown.

(**g**) Predicted relationship between individual chondrocyte sub-clusters determined by pseudo-time trajectory analysis.

(**h**) Expression of exemplar genes of each chondrocyte sub-cluster in pseudo-time.

(**i**) Predicted relationship between individual osteoblast and chondrocyte sub-clusters in diaphysis and metaphysis determined by pseudo-time trajectory analysis.

**Extended Data Fig. 4. Transcription factors implicated in regulating non-haematopoietic sub-clusters.**

(**a**) Heatmap displaying the mean activity of the target genes (regulons) for the top 5 transcription factors predicted by SCENIC to regulate each non-haematopoietic sub-cluster. Scale bar represents AUCell scores which indicate relative expression levels of the regulon across different cell types.

(**b**) UMAP of cell clusters identified by snATAC-seq.

(**c**) Barplots displaying the top enriched transcription factor binding motifs within differentially accessible peaks identified within the 2 osteoblast lineage cell (OLC) clusters.

(**d**) Venn diagram showing overlap of identified transcription factors from SCENIC and snATAC-seq analyses.

(**e**) Barplot displaying skeletal annotations of all transcription factors identified from snATAC-seq analysis. Top transcription factors for both osteoblast lineage clusters identified from snATAC-seq analysis in (**c**) are annotated. Common transcription factors identified from both snATAC-seq and SCENIC analyses are highlighted in bold and underlined. Circos plot displaying known skeletal annotations of all common transcription factors identified from both methods. Motifs involving genes annotated with a skeletal process in the gene ontology (GO) database (green), the mouse genome informatics (MGI) database (purple), in both databases (blue) or are not annotated in either (orange) are shown. Motifs are ordered by strength of enrichment with snATAC-seq analysis, as indicated by the lower bar in barplot and inner circle of circus plot.

**Extended Data Fig. 5. Gene programs of non-haematopoietic cells and osteoclasts are enriched with monogenic skeletal disorder genes**

(**a**) Dotplot showing the mean expression level (log_10_ mean+1) of causative genes of rare monogenic skeletal disorders for individual non-haematopoietic sub-cluster gene programs. Boxes identify exemplar causative genes present in a cell type gene program. Red text denotes a gene found only in a single gene program and not shared with other gene programs. Scale bar in bar plot and UMAP plot indicates the *P* value. Light blue bars in bar plot and light blue dots in UMAP correspond to observations that have nominal evidence of enrichment: *P* value of <0.05 [-log_10_(*P* value) of >1.3]. Dark blue bars and asterisks in bar plot and dark blue dots in UMAP correspond to observations that have robust evidence of enrichment and meet the *Bonferroni*-corrected significance threshold: *P* value of < 3.1 × 10^-3^ [-log_10_(*P* value) > 2.5].

Number and proportion of causative genes within rare disorder groups found in gene programs for osteoblast (**b**) and chondrocyte (**c**) sub-clusters. 27 and 26 disorder groups are identified for osteoblast and chondrocyte sub-clusters respectively, with those enriched with disorder-causing genes in at least one of the sub-clusters indicated in bold. Size of the circles represent the number of genes in each disorder group present within the gene program. Scale bar indicates the *P* value of enrichment, as determined by hypergeometric tests of over-representation. Light blue dots indicate nominal evidence of enrichment: *P* value of <0.05 [-log_10_(*P* value) of >1.3]. Dark blue dots denote robust evidence of enrichment with *Bonferroni*-corrected threshold of 2 × 10^-4^ (**b**) and 4.1 × 10^-4^ (**c**) [-log_10_(*P* value) of >3.7 (**b**) or > 3.4 (**c**)].

(**d**) Network plot showing unique and shared causative genes identified in the gene programs of osteoblast lineage and chondrocyte sub-clusters. Genes are colour coded based on disorder group and the magnitude of expression log_2_(FC) is indicated by the thickness of the connecting lines.

**Extended Data Fig. 6. Genome-wide association study of eBMD in the UK Biobank study**

(**a**) Manhattan plot summarising the results of eBMD GWAS performed on 448,010 participants in the UK Biobank study. The dashed line represents the threshold to declare genome wide significance (*P* <□6.6□×□10^-9^). Novel lead eBMD associated variants are coloured yellow and annotated with the name of the protein coding gene. The y axis is capped at −log_10_(P-value) = 600 and excludes 81 variants located within the *WNT16*/*CPED1* locus on chromosome 7. Variants with *P* >□5.5□×□10^-5^ have also been excluded.

(**b**) Bivariate scatter plot contrasting the magnitude of the genetic effect on eBMD and minor allele frequency of lead eBMD associated variants. Orange circles correspond to lead variants that map to known BMD associated loci, and yellow circles correspond to lead variants that map to novel loci. Gene symbols of monogenic skeletal disorder genes are annotated if they are located closest a lead eBMD associated variant.

(**c**) Bubble plot showing the number, and proportion of monogenic skeletal disorder genes from each disorder group that are present in the set of 901 protein coding genes located closest to lead eBMD associated variants. Size of the circles represent the number of genes in each disorder group present within the gene list. Scale bar indicates the *P* value of enrichment, as determined by hypergeometric over-representation testing. Light blue dots indicate nominal evidence of enrichment: *P* value of <0.05 [-log_10_(*P* value) of >1.3]. Dark blue dots denote robust evidence of enrichment with *Bonferroni*-corrected threshold of 1.2 × 10^-3^ [-log_10_(*P* value) of >2.9].

**Extended Data Fig. 7. Gene programs of non-haematopoietic cell sub-clusters are enriched with eBMD-associated genes**

Bar plot and UMAP plot showing enrichment of eBMD-associated genes in gene programs for different cell types identified in scRNA-seq. Scale bar in bar plot and UMAP plot indicates the *P* value. Light blue bars in bar plot and light blue dots in UMAP correspond to observations that have nominal evidence of enrichment: *P* value of <0.05 [-log_10_(*P* value) of >1.3]. Dark blue bars and asterisks in bar plot and dark blue dots in UMAP correspond to observations that have robust evidence of enrichment and meet the *Bonferroni*-corrected significance threshold: *P* value of < 3 ×10^-3^ [-log_10_(*P* value) > 2.5].

**Extended Data Fig. 8. Gene programs of non-haematopoietic cells and osteoclasts are enriched with genes that cause abnormal bone structure when mutated in mice.**

(**a**) Enrichment of gene programs for genes that cause abnormal bone structure when mutated in mice in the MGI database. Light blue bars represent all genes in each gene program and dark blue bars represent genes in each gene program that are associated with eBMD in the MAGMA analysis. Numbers illustrate numbers of genes with abnormal bone structure phenotypes that are present in the gene program (top) and the total number of genes in each program (bottom). Enrichment was determined by hypergeometric over-representation testing. *Bonferroni*-corrected significance: **** *P*<0.0001, *** *P*<0.001, ** *P*<0.01, * *P*<0.05.

(**b**) Enrichment of gene programs, excluding genes that are known to cause monogenic skeletal disorders, for genes that cause abnormal bone structure when deleted in mice in the MGI database. Light blue bars represent all genes in each gene program excluding causative genes for monogenic skeletal disorder; dark blue bars represent genes in each gene program that are not causative for monogenic skeletal disorders and are associated with eBMD in the MAGMA analysis. Numbers illustrate numbers of genes with abnormal bone structure phenotypes (top) and the total number of genes in each group (bottom). Enrichment was determined by hypergeometric over-representation testing.

(**c**) Enrichment for genes that cause abnormal BMD when deleted in mice in the gene programs for non-haematopoietic cell types and osteoclasts, stratified by magnitude of fold-change in differential expression. Panel (i) includes all genes in the gene programs and panel (ii) the genes that are associated with eBMD in the gene programs. Genes within gene programs are allocated to 6 nested groups based on the magnitude of fold-change which increases from left to right. Numbers illustrate numbers of genes with abnormal bone structure phenotypes (top) and the total number of genes allocated to each group (bottom). Enrichment was determined by hypergeometric over-representation testing. *Bonferroni*-corrected significance: **** *P*<0.0001, *** *P*<0.001, ** *P*<0.01, * *P*<0.05.

(**d**) Enrichment for genes that cause abnormal bone structure when mutated in mice in the gene programs of sub-clusters in selected non-haematopoietic cell types identified in (a) in the MGI database. Light blue bars represent all genes in each gene program and dark blue bars represent genes in each gene program that are associated with eBMD in the MAGMA analysis. Numbers illustrate numbers of genes with abnormal bone structure phenotypes (top) and the total number of genes in each program (bottom). Enrichment was determined by hypergeometric over-representation testing.

(**e**) Enrichment for genes that cause abnormal bone structure when mutated in mice in the gene programs of sub-clusters in selected non-haematopoietic cell types, stratified by magnitude of fold-change in expression. Panel (i) includes all genes in the gene programs and panel (ii) includes genes that are associated with eBMD in the gene programs. Genes within gene programs are allocated to 6 nested groups based on the magnitude of fold-change which increases from left to right. Numbers illustrate numbers of genes with abnormal bone structure phenotypes (top) and the total number of genes allocated to each group (bottom). Enrichment was determined by hypergeometric over-representation testing. *Bonferroni*-corrected significance: **** *P*<0.0001, *** *P*<0.001, ** *P*<0.01, * *P*<0.05.

**Extended Data Fig. 9. Cellular mechanism of skeletal phenotype in *Pls3* deficient mice.**

(**a**) Left panel shows representative upper limb paws from P1 WT and *Pls3^y/-^* mice stained with alcian blue (cartilage) and alizarin red (bone); scale bar = 1mm. Second panel shows decalcified proximal tibia growth plate sections from P21 (WT n=5; *Pls3^y/-^* n=4) mice stained with alcian blue (cartilage) and van Gieson (bone osteoid); scale bar = 100μm. Left graph shows the absolute widths of the resting zone (RZ), proliferating zone (PZ) and hypertrophic zone (HZ) chondrocytes in the growth plate. Graphs show femur length from (P1 (WT n=6; *Pls3^y/-^* n=4), P21 (WT n=17; *Pls3^y/-^*n=8), P70 (WT n=10; *Pls3^y/-^* n=6), and P183 (WT n=12; *Pls3^y/-^* n=7) and nose to tail length from P21 (WT n=19; *Pls3^y/-^* n=8), P70 (WT n=8; *Pls3^y/-^* n=6), and P183 (WT n=12; *Pls3^y/-^* n=7); mean±SD;. Students’ *t*-test; \**P*<0.05; \*\**P*<0.01.

(**b**) Confocal images of mid femur cortical bone, distal femur trabecular bone, and lumbar vertebral trabecular bone (L1-L3) from P70 (WT n=6; *Pls3^y/-^*n=6) mice, double-labelled with calcein; scale bar=100μm. Graphs show serum procollagen type 1 N propeptide (P1NP) levels P70 (WT n=10; *Pls3^y/-^* n=6) mice, femur cortical bone mineral apposition rate (MAR), femur trabecular bone mineralising surface per bone surface (MS/BS), MAR and bone formation rate (BFR), and lumbar vertebral trabecular bone MS/BS, MAR and BFR in P70 (WT n=6; *Pls3^y/-^* n=6) mice; mean±SD.

(**c**) Decalcified sections of proximal tibia stained for tartrate-resistant acid phosphatase (TRAP) (osteoclasts) and aniline blue (bone) from P70 from (WT n=10; *Pls3^y/-^* n=6) mice; scale bar=100μm. Graphs show serum C-terminal telopeptide of type 1 collagen (CTX) levels P70 (WT n=10; *Pls3^y/-^* n=6) mice, numbers of osteoclasts per mm bone surface (OcN/BS), and osteoclast surface per mm bone surface (OcS/BS) in P70 (WT n=8; *Pls3^y/-^* n=8) mice; mean±SD.

(**d**) Graphs show serum sclerostin (SOST) and dickkopf-related protein 1 (DKK1) levels in P183 (WT n=11; *Pls3^y/-^*n=7) mice.

(**e**) Micro-CT images of a 250µm mid-femur ROI from P112 (WT n=6; *Pls3^y/-^* n=6) mice, showing small osteocyte lacunae in blue (100-300µm^3^) and large osteocyte lacunae in yellow (301-2000 µm^3^); scale bar = 100μm. Graphs show cortical microporosity (Ct.µPo), lacuna number per bone volume (Lc.N/BV), mean lacuna volume (Mean Lc.V), and lacuna number by lacuna volume; mean±SD; Students’ *t*-test;. \**P*<0.05. The violin plot shows the distribution of lc.V; Kolmogorov-Smirnov test; \*\*\*\**P*<0.0001.

**Extended Data Fig. 10. Spatial transcriptomics of human bone samples.**

(**a**) UMAP plot and of non-haematopoietic cell types and osteoclasts identified in trephine biopsies of 4 control patients as identified by Yip and Er *et al*^52^. MSCs = mesenchymal stromal cells; VSMCs = vascular smooth muscle cells; SECs = sinusoidal endothelial cells; AECs = arterial endothelial cells.

(**b**) Haematoxylin and eosin (H&E) images and spatial plots of individual samples used for spatial transcriptomics. Spatial plots display only non-haematopoietic cells and osteoclasts for clarity and are coloured according to (**a**).

Bar plots showing enrichment of gene programs for (**c**) causative genes of monogenic skeletal disorders and (**d**) eBMD-associated genes. Scale bars in bar plots indicate the *P* value. Light blue bars in bar plot correspond to observations that have nominal evidence of enrichment: *P* value of <0.05 [-log_10_(*P* value) of >1.3]. Dark blue bars and asterisks in bar plot correspond to observations that have robust evidence of enrichment and meet the *Bonferroni*-corrected significance threshold: *P* value of < 1.6 ×10^-3^ [-log_10_(*P* value) > 2.3].

**Supplementary Fig. 1. Validation of cell populations enriched in the endosteal compartment of mouse bone.**

(**a**) UMAP plots displaying expression of *Cd14* within the mouse scRNA-Seq dataset. Clusters “7. Monocytes (*Cd14^+^*)” and “1. Mature neutrophils (*Cxcl2^+^*)” are indicated.

(**b**) Top marker genes for clusters “7. Monocytes (*Cd14^+^*)” and “1. Mature neutrophils (*Cxcl2^+^*)”. *Cd14* is highlighted as a strong marker for both clusters.

(**c**) Gating strategy used to identify neutrophils and monocyte populations in mouse bone marrow and endosteal samples and to determine *CD14* expression within these populations.

(**d**) Quantification of *CD14* expression in selected populations. Left bar plots show the proportion of cells with detectable *CD14* expression. Right bar plots show median fluorescence intensity of *CD14* within selected populations. Mean ± SEM are shown; Students’ *t*-test; *** *P*<0.001, ** *P*<0.01, * *P*<0.05.

**Supplementary Fig. 2. Distribution of non-haematopoietic cell sub-clusters in diaphysis and metaphysis.**

Bar plots showing the fraction of all non-haematopoietic sub-clusters in diaphysis and metaphysis. Mean ± SEM are shown; Students’ *t*-test; **** *P*<0.0001, *** *P*<0.001, ** *P*<0.01, * *P*<0.05.

**Supplementary Fig. 3. Genes associated with eBMD, but not pulse rate, are enriched with monogenic skeletal disorder genes.**

Bubble plot showing the number, and proportion of monogenic skeletal disorder genes from each disorder group that are present in the set of protein-coding genes associated with (**a**) eBMD and (**b**) pulse rate. Size of the circles represent the number of genes in each disorder group present within the gene list. Scale bar indicates the *P* value of enrichment, as determined by hypergeometric over-representation testing. Light blue dots indicate nominal evidence of enrichment: *P* value of <0.05 [-log_10_(*P* value) of >1.3]. Dark blue dots denote robust evidence of enrichment with *Bonferroni*-corrected threshold of 1.2 × 10^-3^ [-log_10_(*P* value) of >2.9].

**Supplementary Fig. 4. Conditional GSA analysis to determine whether enrichment was confounded by shared sets of genes.**

(**a**) UpSet plot showing the number of restricted and shared genes associated with eBMD in non-haematopoietic cell types.

(**b**) Histograms showing a pairwise conditional GSA analysis used to determine whether enrichment was confounded by shared sets of genes. Each histogram quantifies the strength of evidence of enrichment. Dark grey bars correspond to strength of evidence of enrichment in marginal (original) analyses and light grey bars correspond to the strength of evidence of enrichment after adjusting for the effect of genes that are shared between two cell types (conditional analyses). Numbers above each bar correspond to the GSA point estimate (i.e. β). **** *P*<0.001, *** *P*<0.005** *P*<0.01, * *P*<0.05. Dotted line corresponds to the threshold of statistical significance (*P* < 0.05).

(**c**) Post-hoc permutation analyses showing QQ-plots of Z-scores of genes in each gene program of different cell clusters. Plots show residualised Z-scores from the null model for each gene program, with the expected values based on the quantiles across all genes in the data. The black points denote the 25th, 50th and 75th percentile. The dashed black line represents the one-sided (upper) 95% confidence band. Genes are coloured red if they exceed the confidence band, and grey if they do not. The proportion of genes in each program exceeding the confidence band is indicated on each plot.

**Supplementary Fig. 5. Expression of exemplar genes in tissues outside of the skeleton.** Dotplots showing expression of exemplar genes across cell types isolated from different mouse (**a**) and human (**b**) tissues, using the TabulaMuris^88^ and TabulaSapiens^89^ datasets respectively. Individual cell types were manually assigned a classification using the Human Protein Atlas^91^ and coloured accordingly. Left dotted lines indicate the mean expression value of a gene across the whole dataset; right dotted lines indicate the 90^th^ percentile value.

**Supplementary Fig. 6. Skeletal phenotype of younger male *Pls3^y/-^* mice.**

(**a**) Pseudocoloured X-ray microradiography of lumbar vertebrae (L5) from P70 (WT n=5; *Pls3^y/-^* n=5) mice; scale bar = 1mm. Relative frequency histograms of BMC are shown for each age comparison; Kolmogorov-Smirnov test, \*\*\**P*<0.001. Micro-CT images of mid coronal sections of lumbar vertebrae (L5) from P70 (WT n=5; *Pls3^y/-^*n=5) mice; scale bar = 1mm. Graphs show bone volume as a proportion of tissue volume (BV/TV), trabecular number (Tb.N), trabecular thickness (Tb.Th), and trabecular separation (Tb.Sp); mean±SD; Students’ *t*-test.

(**b**) Representative load displacement curves from compression testing of lumbar vertebrae (L5) from P70 (WT n=5; *Pls3^y/-^*n=5) mice. Graphs show yield load, maximum load, and stiffness; mean±SD; Students’ *t*-test; \**P*<0.05.

(**c**) Pseudocoloured X-ray microradiography of femurs from P21 (WT, blue, n=12; *Pls3^y/-^*, orange, n=4), and P70 (WT n=7; *Pls3^y/-^* n=6). Low bone mineral content (BMC) is blue/green and high BMC is pink; scale bar = 1mm. Relative frequency histograms of BMC are shown for each age comparison; Kolmogorov-Smirnov test, \*\*\**P*<0.001.

(**d**) Micro-CT images of distal femur trabecular bone from P21 (WT n=14; *Pls3^y/-^* n=4), and P70 (WT n=10; *Pls3^y/-^* n=6) mice; scale bar = 100μm. Graphs show bone volume as a proportion of tissue volume (BV/TV), trabecular number (Tb.N), trabecular thickness (Tb.Th), and trabecular separation (Tb.Sp); mean±SD; Students’ *t*-test; \**P*<0.05.

(**e**) Micro-CT images of femur mid-shaft cortical bone from P21 (WT n=14; *Pls3^y/-^* n=4), P70 (WT n=10; *Pls3^y/-^* n=6) mice; scale bar = 100μm. Graphs show cortical thickness (Ct.Th), internal diameter (Int.Di), and bone mineral density (Ct.BMD); mean±SD; Students’ *t*-test; \*\*\*\**P*<0.0001.

(**f**) Representative load displacement curves from three-point bend testing of femurs from P70 (WT, blue, n=10; *Pls3^y/-^*, orange, n=6) mice. Graphs show yield load, maximum load, fracture load, and stiffness; mean±SD; Students’ *t*-test; \**P*<0.05, \*\*\**P*<0.001.

**Supplementary Fig. 7. Skeletal phenotype of female *Pls3^-/-^* mice.**

(**a**) Greyscale and pseudocoloured X-ray microradiography of femurs from P112 WT and *Pls3^-/-^*mice. Low bone mineral content (BMC) is blue/green and high BMC is pink; scale bar = 1mm. Graphs show relative bone mineral content (BMC, mean±SD) and femur length (median±2.5th and 97.5th percentiles) with WT reference range represented as grey violin plots with individual values shown as white dots (n=320). Individual values, together with the mean or median, from *Pls3^-/-^* (orange n=2) are shown. Significant *P* values after permutation testing are indicated.

(**b**) Caudal vertebrae from P112 WT and *Pls3^-/-^* mice; scale bar = 1mm. Graphs show BMC (mean±SD), and vertebral length (mean±SD) with WT reference range represented as grey violin plots with individual values shown as white dots (n=320). Individual values and mean from *Pls3^-/-^* (orange n=2) mice are shown.

(**c**) Micro-CT images of distal femur trabecular bone from P112 WT and *Pls3^-/-^* mice; scale bar = 100μm. Graphs show bone volume as a proportion of tissue volume (BV/TV, mean±SD), trabecular number (Tb.N, mean±SD), trabecular thickness (Tb.Th, mean±SD), and trabecular separation (Tb.Sp, median±2.5th and 97.5th percentiles). Significant *P* values after permutation testing are indicated.

(**d**) Micro-CT images of femur mid-shaft cortical bone from P112 WT and *Pls3^-/-^* mice; scale bar = 100μm. Graphs show cortical thickness (Ct.Th, mean±SD), internal diameter (Int.Di, mean±SD), and bone mineral density (Ct.BMD, median±2.5th and 97.5th percentiles). Significant *P* values after permutation testing are indicated.

(**e**) Representative load displacement curves from three-point bend testing of femurs from P112 WT (blue) and *Pls3^-/-^*(orange) mice. Graphs show yield load (mean±SD), maximum load (mean±SD), fracture load (median±2.5th and 97.5th percentiles), and stiffness (mean±SD). Significant *P* values after permutation testing are indicated.

(**f**) Representative load displacement curves from compression testing of caudal vertebrae from P112 WT (blue) and *Pls3^-/-^* (orange) mice. Graphs show yield load, maximum load, and stiffness (mean±SD).

**Supplementary Fig. 8. Skeletal vascular phenotype of male Pls3y/- mice**.

(**a**) Micro-CT images (1.0 µm^3^ voxel resolution) of mid-femur cortical bone (250 µm long ROI) from P112 (WT n=6; *Pls3^y/-^* n=6) mice; scale bar = 100μm.

(**b**) Osteocyte lacunae and cortical vascular canals, within the cortical bone ROI, with a volume greater than 100 µm^3^ identified using BoneJ Particle Analyser.

(**c**) Cortical vascular canals (Red), within the cortical bone ROI, with a volume greater than 2000 µm^3^ identified using BoneJ Particle Analyser.

(**e**) Overlay of mid-femur cortical bone ROI and vascular canals

(**f**) Higher power image of cortical bone and vascular canals.

## References

1. Dieleman, J.L. et al. US Health Care Spending by Payer and Health Condition, 1996-2016. Jama 323, 863-884 (2020).

2. Wu, A.-M. et al. Global, regional, and national burden of bone fractures in 204 countries and territories, 1990–2019: a systematic analysis from the Global Burden of Disease Study 2019. The Lancet Healthy Longevity 2, e580-e592 (2021).

3. Ayub, N. et al. The Treatment Gap in Osteoporosis. J Clin Med 10(2021).

4. Minikel, E.V., Painter, J.L., Dong, C.C. & Nelson, M.R. Refining the impact of genetic evidence on clinical success. Nature 629, 624–629 (2024).

5. Rivadeneira, F. & Mäkitie, O. Osteoporosis and Bone Mass Disorders: From Gene Pathways to Treatments. Trends Endocrinol Metab 27, 262–281 (2016).

6. Zhu, X., Bai, W. & Zheng, H. Twelve years of GWAS discoveries for osteoporosis and related traits: advances, challenges and applications. Bone Res 9, 23 (2021).

7. Kemp, J.P. et al. Identification of 153 new loci associated with heel bone mineral density and functional involvement of GPC6 in osteoporosis. Nat Genet 49, 1468–1475 (2017).

8. Morris, J.A. et al. An atlas of genetic influences on osteoporosis in humans and mice. Nat Genet 51, 258–266 (2019).

9. Kim, S.K. Identification of 613 new loci associated with heel bone mineral density and a polygenic risk score for bone mineral density, osteoporosis and fracture. PLOS ONE 13, e0200785 (2018).

10. Aguet, F. et al. Molecular quantitative trait loci. Nature Reviews Methods Primers 3, 4 (2023).

11. Mullin, B.H. et al. Characterisation of genetic regulatory effects for osteoporosis risk variants in human osteoclasts. Genome Biol 21, 80 (2020).

12. Grundberg, E. et al. Population genomics in a disease targeted primary cell model. Genome Res 19, 1942–52 (2009).

13. Al-Barghouthi, B.M. et al. Transcriptome-wide association study and eQTL colocalization identify potentially causal genes responsible for human bone mineral density GWAS associations. eLife 11, e77285 (2022).

14. Yazar, S. et al. Single-cell eQTL mapping identifies cell type–specific genetic control of autoimmune disease. Science 376, eabf3041 (2022).

15. Bandyopadhyay, S. et al. Mapping the cellular biogeography of human bone marrow niches using single-cell transcriptomics and proteomic imaging. Cell 187, 3120–3140.e29 (2024).

16. Rood, J.E., Maartens, A., Hupalowska, A., Teichmann, S.A. & Regev, A. Impact of the Human Cell Atlas on medicine. Nature Medicine 28, 2486–2496 (2022).

17. Consortium, T.G. et al. The GTEx Consortium atlas of genetic regulatory effects across human tissues. Science 369, 1318–1330 (2020).

18. Chinwalla, A.T. et al. Initial sequencing and comparative analysis of the mouse genome. Nature 420, 520–562 (2002).

19. Brommage, R. & Ohlsson, C. High Fidelity of Mouse Models Mimicking Human Genetic Skeletal Disorders. Frontiers in Endocrinology 10(2020).

20. Kaya, S., Schurman, C.A., Dole, N.S., Evans, D.S. & Alliston, T. Prioritization of Genes Relevant to Bone Fragility Through the Unbiased Integration of Aging Mouse Bone Transcriptomics and Human GWAS Analyses. J Bone Miner Res 37, 804–817 (2022).

21. Doolittle, M.L., Khosla, S. & Saul, D. Single-Cell Integration of BMD GWAS Results Prioritize Candidate Genes Influencing Age-Related Bone Loss. JBMR Plus 7, e10795 (2023).

22. Unger, S. et al. Nosology of genetic skeletal disorders: 2023 revision. American Journal of Medical Genetics Part A 191, 1164–1209 (2023).

23. Batoon, L. et al. CD169+ macrophages are critical for osteoblast maintenance and promote intramembranous and endochondral ossification during bone repair. Biomaterials 196, 51–66 (2019).

24. McDonald, M.M. et al. Osteoclasts recycle via osteomorphs during RANKL-stimulated bone resorption. Cell 184, 1330–1347.e13 (2021).

25. Millard, S.M. et al. Fragmentation of tissue-resident macrophages during isolation confounds analysis of single-cell preparations from mouse hematopoietic tissues. Cell Reports 37, 110058 (2021).

26. Hume, D.A., Millard, S.M. & Pettit, A.R. Macrophage heterogeneity in the single-cell era: facts and artifacts. Blood 142, 1339–1347 (2023).

27. Filipowska, J., Tomaszewski, K.A., Niedźwiedzki, Ł., Walocha, J.A. & Niedźwiedzki, T. The role of vasculature in bone development, regeneration and proper systemic functioning. Angiogenesis 20, 291–302 (2017).

28. Lane, M.D., Tang, Q.-Q. & Jiang, M.-S. Role of the CCAAT Enhancer Binding Proteins (C/EBPs) in Adipocyte Differentiation. Biochemical and Biophysical Research Communications 266, 677–683 (1999).

29. Ouyang, N. et al. The Transcription Factor Foxc1 Promotes Osteogenesis by Directly Regulating Runx2 in Response of Intermittent Parathyroid Hormone (1-34) Treatment. Front Pharmacol 11, 592 (2020).

30. Rashid, H. et al. Sp7 and Runx2 molecular complex synergistically regulate expression of target genes. Connect Tissue Res 55 Suppl 1, 83–7 (2014).

31. Wang, S. et al. Regulon active landscape reveals cell development and functional state changes of human primary osteoblasts in vivo. Human Genomics 17, 11 (2023).

32. Zhang, J. et al. Neural tube, skeletal and body wall defects in mice lacking transcription factor AP-2. Nature 381, 238–41 (1996).

33. Kieslinger, M. et al. EBF2 regulates osteoblast-dependent differentiation of osteoclasts. Dev Cell 9, 757–67 (2005).

34. Thottappillil, N. et al. ZIC1 Dictates Osteogenesis Versus Adipogenesis in Human Mesenchymal Progenitor Cells Via a Hedgehog Dependent Mechanism. Stem Cells 41, 862–876 (2023).

35. Kusumbe, A.P., Ramasamy, S.K. & Adams, R.H. Coupling of angiogenesis and osteogenesis by a specific vessel subtype in bone. Nature 507, 323–328 (2014).

36. Klingseisen, A. & Jackson, A.P. Mechanisms and pathways of growth failure in primordial dwarfism. Genes Dev 25, 2011–24 (2011).

37. Zhou, S. et al. Converging evidence from exome sequencing and common variants implicates target genes for osteoporosis. Nature Genetics 55, 1277–1287 (2023).

38. Wang, Q. et al. Rare variant contribution to human disease in 281,104 UK Biobank exomes. Nature 597, 527–532 (2021).

39. Dijk, F.S.v., et al. *PLS3* Mutations in X-Linked Osteoporosis with Fractures. New England Journal of Medicine 369, 1529–1536 (2013).

40. Döffinger, R. et al. X-linked anhidrotic ectodermal dysplasia with immunodeficiency is caused by impaired NF-κB signaling. Nature Genetics 27, 277–285 (2001).

41. Francis, F. et al. A gene (PEX) with homologies to endopeptidases is mutated in patients with X–linked hypophosphatemic rickets. Nature Genetics 11, 130–136 (1995).

42. de Leeuw, C.A., Mooij, J.M., Heskes, T. & Posthuma, D. MAGMA: generalized gene-set analysis of GWAS data. PLoS Comput Biol 11, e1004219 (2015).

43. Bassett, J.H. et al. Rapid-throughput skeletal phenotyping of 100 knockout mice identifies 9 new genes that determine bone strength. PLoS Genet 8, e1002858 (2012).

44. Bianco, P., Fisher, L.W., Young, M.F., Termine, J.D. & Robey, P.G. Expression and localization of the two small proteoglycans biglycan and decorin in developing human skeletal and non-skeletal tissues. Journal of Histochemistry & Cytochemistry 38, 1549–1563 (1990).

45. Keller, R.B. et al. Monoallelic and biallelic CREB3L1 variant causes mild and severe osteogenesis imperfecta, respectively. Genet Med 20, 411–419 (2018).

46. Schwebach, C.L. et al. Osteogenesis imperfecta mutations in plastin 3 lead to impaired calcium regulation of actin bundling. Bone Research 8, 21 (2020).

47. Chin, S.M., Unnold-Cofre, C., Naismith, T. & Jansen, S. The actin-bundling protein, PLS3, is part of the mechanoresponsive machinery that regulates osteoblast mineralization. Front Cell Dev Biol 11, 1141738 (2023).

48. Maus, I., et al. Osteoclast-specific Plastin 3 knockout in mice fail to develop osteoporosis despite dramatic increased osteoclast resorption activity. JBMR Plus 8(2024).

49. Yorgan, T.A. et al. Mice lacking plastin-3 display a specific defect of cortical bone acquisition. Bone 130, 115062 (2020).

50. Wu, R.S. et al. A Rapid Method for Directed Gene Knockout for Screening in G0 Zebrafish. Dev Cell 46, 112–125.e4 (2018).

51. To, K. et al. A multi-omic atlas of human embryonic skeletal development. Nature 635, 657–667 (2024).

52. Yip, R.K.H. et al. Profiling the spatial architecture of multiple myeloma in human bone marrow trephine biopsy specimens with spatial transcriptomics. Blood (2025).

53. Perrin, S. et al. Single nuclei transcriptomics reveal the differentiation trajectories of periosteal skeletal/stem progenitor cells in bone regeneration. (eLife Sciences Publications, Ltd, 2024).

54. Tikhonova, A.N. et al. The bone marrow microenvironment at single-cell resolution. Nature 569, 222–228 (2019).

55. Baryawno, N. et al. A Cellular Taxonomy of the Bone Marrow Stroma in Homeostasis and Leukemia. Cell 177, 1915–1932.e16 (2019).

56. Baccin, C. et al. Combined single-cell and spatial transcriptomics reveal the molecular, cellular and spatial bone marrow niche organization. Nature Cell Biology 22, 38–48 (2020).

57. Zhong, L. et al. Single cell transcriptomics identifies a unique adipose lineage cell population that regulates bone marrow environment. eLife 9, e54695 (2020).

58. Nookaew, I. et al. Refining the identity of mesenchymal cell types associated with murine periosteal and endosteal bone. Journal of Biological Chemistry 300, 107158 (2024).

59. Ayturk, U.M. et al. SingleCCell RNA Sequencing of Calvarial and LongCBone Endocortical Cells. Journal of Bone and Mineral Research 35, 1981–1991 (2020).

60. Youlten, S.E. et al. Osteocyte transcriptome mapping identifies a molecular landscape controlling skeletal homeostasis and susceptibility to skeletal disease. Nat Commun 12, 2444 (2021).

61. Hao, Y. et al. Integrated analysis of multimodal single-cell data. Cell 184, 3573–3587.e29 (2021).

62. Korsunsky, I. et al. Fast, sensitive and accurate integration of single-cell data with Harmony. Nature Methods 16, 1289–1296 (2019).

63. Menezes, S. et al. The Heterogeneity of Ly6C(hi) Monocytes Controls Their Differentiation into iNOS(+) Macrophages or Monocyte-Derived Dendritic Cells. Immunity 45, 1205–1218 (2016).

64. Xie, X. et al. Single-cell transcriptome profiling reveals neutrophil heterogeneity in homeostasis and infection. Nature Immunology 21, 1119–1133 (2020).

65. Lee, R.D. et al. Single-cell analysis identifies dynamic gene expression networks that govern B cell development and transformation. Nature Communications 12, 6843 (2021).

66. Troulé, K. et al. CellPhoneDB v5: inferring cell–cell communication from single-cell multiomics data. Nature Protocols (2025).

67. Drost, H.G. & Paszkowski, J. Biomartr: genomic data retrieval with R. Bioinformatics 33, 1216–1217 (2017).

68. Milacic, M. et al. The Reactome Pathway Knowledgebase 2024. Nucleic Acids Research 52, D672–D678 (2024).

69. Yu, G. & He, Q.-Y. ReactomePA: an R/Bioconductor package for reactome pathway analysis and visualization. Molecular BioSystems 12, 477–479 (2016).

70. Wickham, H. ggplot2: elegant graphics for data analysis. (Springer-Verlag, New York, 2016).

71. Trapnell, C. et al. The dynamics and regulators of cell fate decisions are revealed by pseudotemporal ordering of single cells. Nat Biotechnol 32, 381–386 (2014).

72. Stuart, T. et al. Comprehensive Integration of Single-Cell Data. Cell 177, 1888–1902.e21 (2019).

73. Aibar, S. et al. SCENIC: single-cell regulatory network inference and clustering. Nat Methods 14, 1083–1086 (2017).

74. Ashburner, M. et al. Gene Ontology: tool for the unification of biology. Nature Genetics 25, 25–29 (2000).

75. Smith, C.L., Blake, J.A., Kadin, J.A., Richardson, J.E. & Bult, C.J. Mouse Genome Database (MGD)-2018: knowledgebase for the laboratory mouse. Nucleic Acids Res 46, D836–d842 (2018).

76. Durinck, S., Spellman, P.T., Birney, E. & Huber, W. Mapping identifiers for the integration of genomic datasets with the R/Bioconductor package biomaRt. Nature Protocols 4, 1184–1191 (2009).

77. Zimmerman, M.T., Kabat, B., Grill, D.E., Kennedy, R.B. & Poland, G.A. RITAN: rapid integration of term annotation and network resources. PeerJ 7(2019).

78. Shannon, P. et al. Cytoscape: a software environment for integrated models of biomolecular interaction networks. Genome Res 13, 2498–504 (2003).

79. Deng, J.E., Sham, P.C. & Li, M.X. SNPTracker: A Swift Tool for Comprehensive Tracking and Unifying dbSNP rs IDs and Genomic Coordinates of Massive Sequence Variants. G3 (Bethesda) 6, 205–7 (2015).

80. Cingolani, P. et al. A program for annotating and predicting the effects of single nucleotide polymorphisms, SnpEff: SNPs in the genome of Drosophila melanogaster strain w1118; iso-2; iso-3. Fly (Austin) 6, 80–92 (2012).

81. McLaren, W. et al. The Ensembl Variant Effect Predictor. Genome Biol 17, 122 (2016).

82. Yang, J. et al. Conditional and joint multiple-SNP analysis of GWAS summary statistics identifies additional variants influencing complex traits. Nat Genet 44, 369–75, s1-3 (2012).

83. Yang, J., Lee, S.H., Goddard, M.E. & Visscher, P.M. GCTA: a tool for genome-wide complex trait analysis. Am J Hum Genet 88, 76–82 (2011).

84. Bycroft, C. et al. The UK Biobank resource with deep phenotyping and genomic data. Nature 562, 203–209 (2018).

85. Manichaikul, A. et al. Robust relationship inference in genome-wide association studies. Bioinformatics 26, 2867–73 (2010).

86. Purcell, S. et al. PLINK: a tool set for whole-genome association and population-based linkage analyses. Am J Hum Genet 81, 559–75 (2007).

87. Quinlan, A.R. & Hall, I.M. BEDTools: a flexible suite of utilities for comparing genomic features. Bioinformatics 26, 841–2 (2010).

88. Schaum, N. et al. Single-cell transcriptomics of 20 mouse organs creates a Tabula Muris. Nature 562, 367–372 (2018).

89. Consortium, T.T.S. & Quake, S.R. Tabula Sapiens reveals transcription factor expression, senescence effects, and sex-specific features in cell types from 28 human organs and tissues. bioRxiv, 2024.12.03.626516 (2025).

90. Huang, N. sceasy: A package to help convert different single-cell data formats to each other. 0.0.7 edn (2025).

91. Uhlén, M. et al. Tissue-based map of the human proteome. Science 347, 1260419 (2015).

92. Kolde, R. & Kolde, M.R. Package ‘pheatmap’. R package 1, 790 (2015).

93. Kilkenny, C., Browne, W.J., Cuthill, I.C., Emerson, M. & Altman, D.G. Improving bioscience research reporting: the ARRIVE guidelines for reporting animal research. PLoS Biol 8, e1000412 (2010).

94. Skarnes, W.C. et al. A conditional knockout resource for the genome-wide study of mouse gene function. Nature 474, 337–42 (2011).

95. White, J.K. et al. Genome-wide generation and systematic phenotyping of knockout mice reveals new roles for many genes. Cell 154, 452–64 (2013).

96. Butterfield, N.C., Logan, J.G., Waung, J., Williams, G.R. & Bassett, J.H.D. Quantitative X-Ray Imaging of Mouse Bone by Faxitron. Methods Mol Biol 1914, 559–569 (2019).

97. Butterfield, N.C. et al. Accelerating functional gene discovery in osteoarthritis. Nat Commun 12, 467 (2021).

98. Doube, M. et al. BoneJ: Free and extensible bone image analysis in ImageJ. Bone 47, 1076–1079 (2010).

99. Hemmatian, H. et al. Age-related changes in female mouse cortical bone microporosity. Bone 113, 1–8 (2018).

100. Esapa, C.T. et al. Bone Mineral Content and Density. Curr Protoc Mouse Biol 2, 365–400 (2012).

101. Rigueur, D. & Lyons, K.M. Whole-mount skeletal staining. Methods Mol Biol 1130, 113–121 (2014).

102. Bassett, J.H. et al. Mice lacking the calcineurin inhibitor Rcan2 have an isolated defect of osteoblast function. Endocrinology 153, 3537–48 (2012).

103. van’t Hof, R.J., Rose, L., Bassonga, E. & Daroszewska, A. Open source software for semi-automated histomorphometry of bone resorption and formation parameters. Bone 99, 69–79 (2017).

104. Dempster, D.W. et al. Standardized nomenclature, symbols, and units for bone histomorphometry: a 2012 update of the report of the ASBMR Histomorphometry Nomenclature Committee. J Bone Miner Res 28, 2–17 (2013).

105. Bassett, J.H. et al. Thyrostimulin Regulates Osteoblastic Bone Formation During Early Skeletal Development. Endocrinology 156, 3098–113 (2015).

106. Bassett, J.H., van der Spek, A., Gogakos, A. & Williams, G.R. Quantitative X-ray imaging of rodent bone by Faxitron. Methods Mol Biol 816, 499–506 (2012).

107. Beis, D. et al. Genetic and cellular analyses of zebrafish atrioventricular cushion and valve development. Development 132, 4193–204 (2005).

108. McDowell, K.P., Berthiaume, A.A., Tieu, T., Hartmann, D.A. & Shih, A.Y. VasoMetrics: unbiased spatiotemporal analysis of microvascular diameter in multi-photon imaging applications. Quant Imaging Med Surg 11, 969–982 (2021).

