## Extended Data Figure 1 for "Multiscale analysis and functional validation of the cellular and genetic determinants of skeletal disease"

### Extended Data Fig. 1. Genes and gene programs that define chondrocytes, endothelial cells, vascular smooth muscle cells and osteoclasts

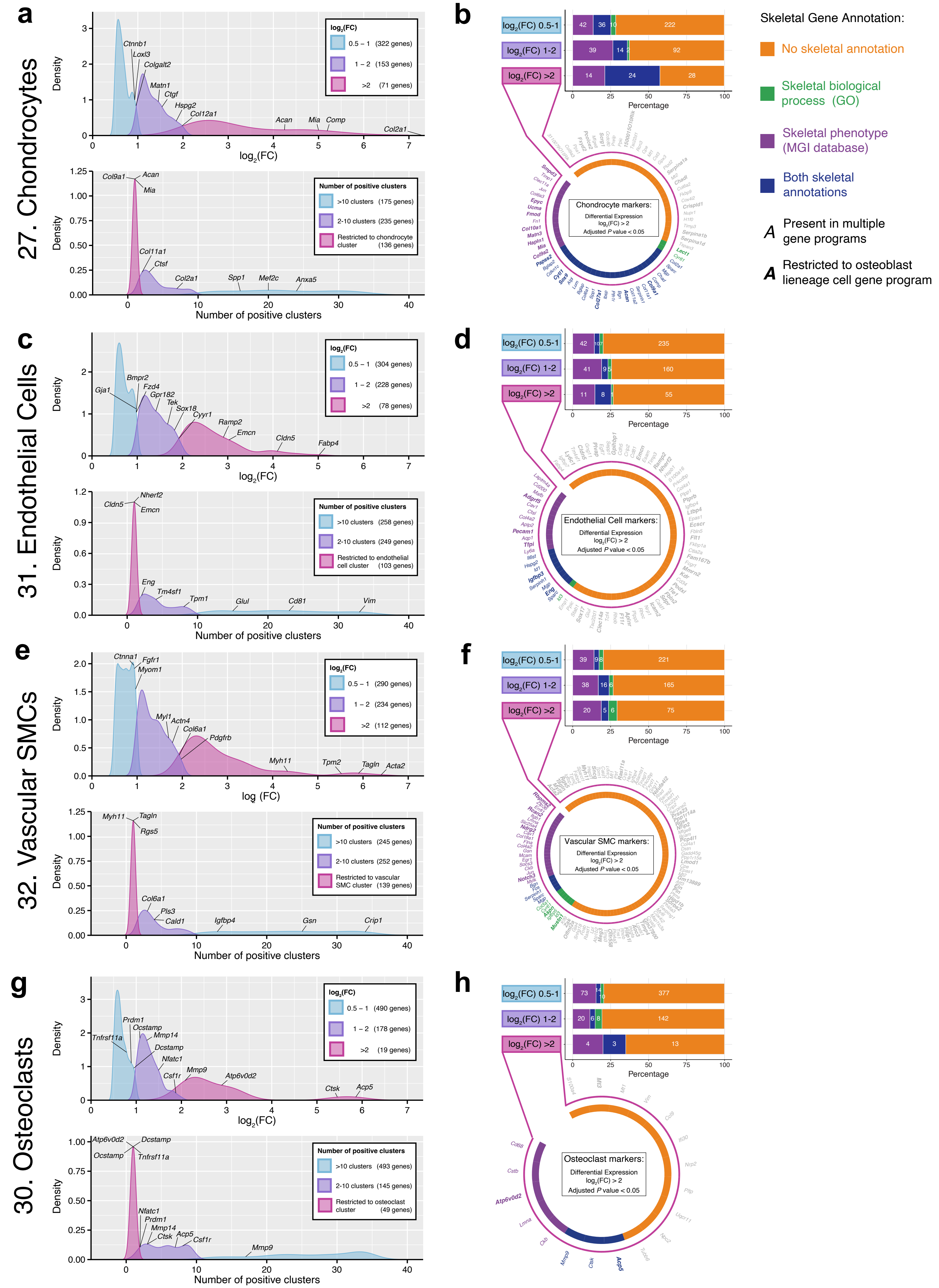
