## Extended Data Figure 2 for "Multiscale analysis and functional validation of the cellular and genetic determinants of skeletal disease"

### Extended Data Fig. 2. Cell-cell interaction of osteoblast lineage cells with other non-haematopoietic cells and osteoclasts

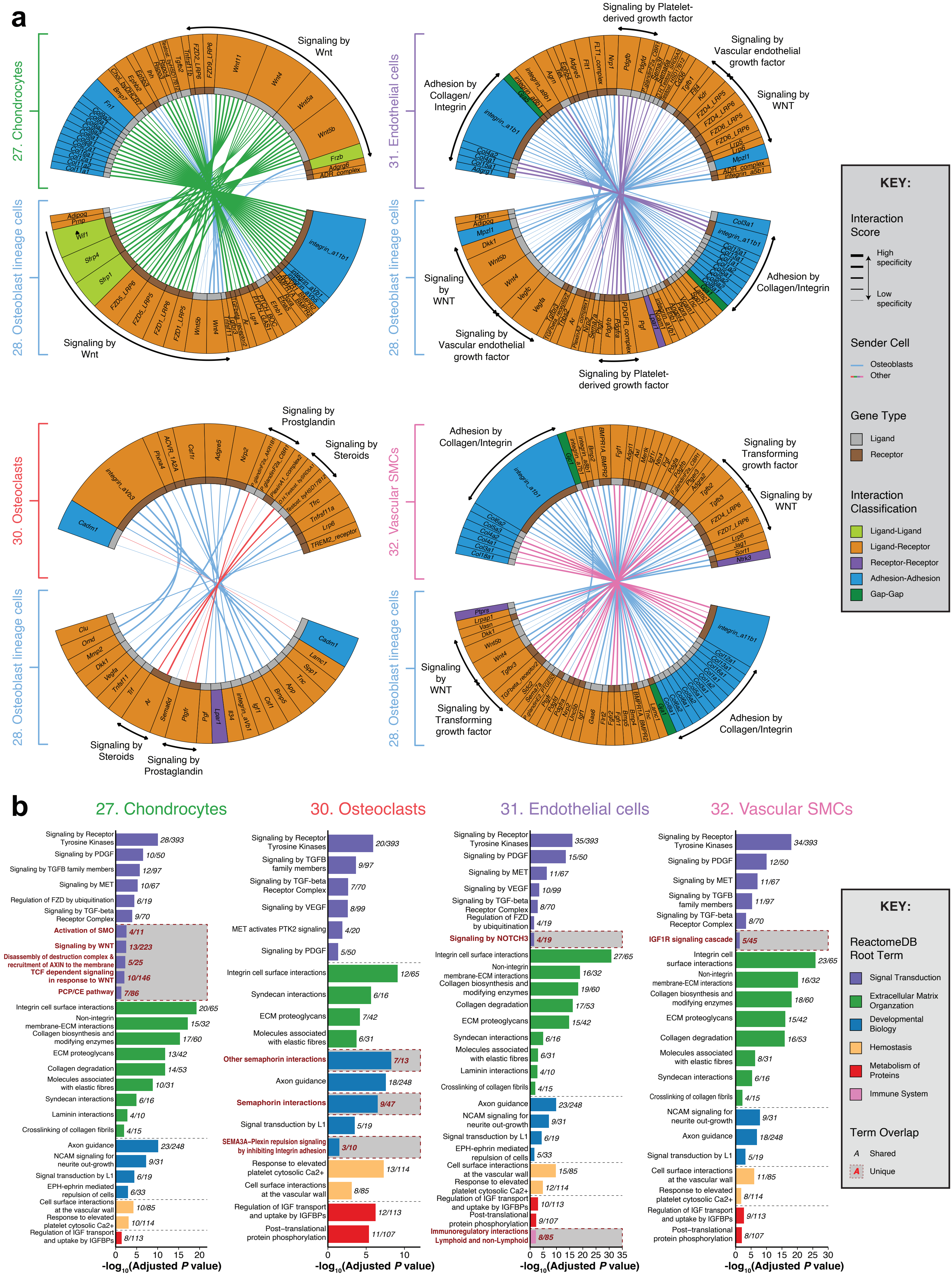
