## Extended Data Figure 3 for "Multiscale analysis and functional validation of the cellular and genetic determinants of skeletal disease"

### Extended Data Fig. 3. High-resolution clustering of non-haematopoietic cells

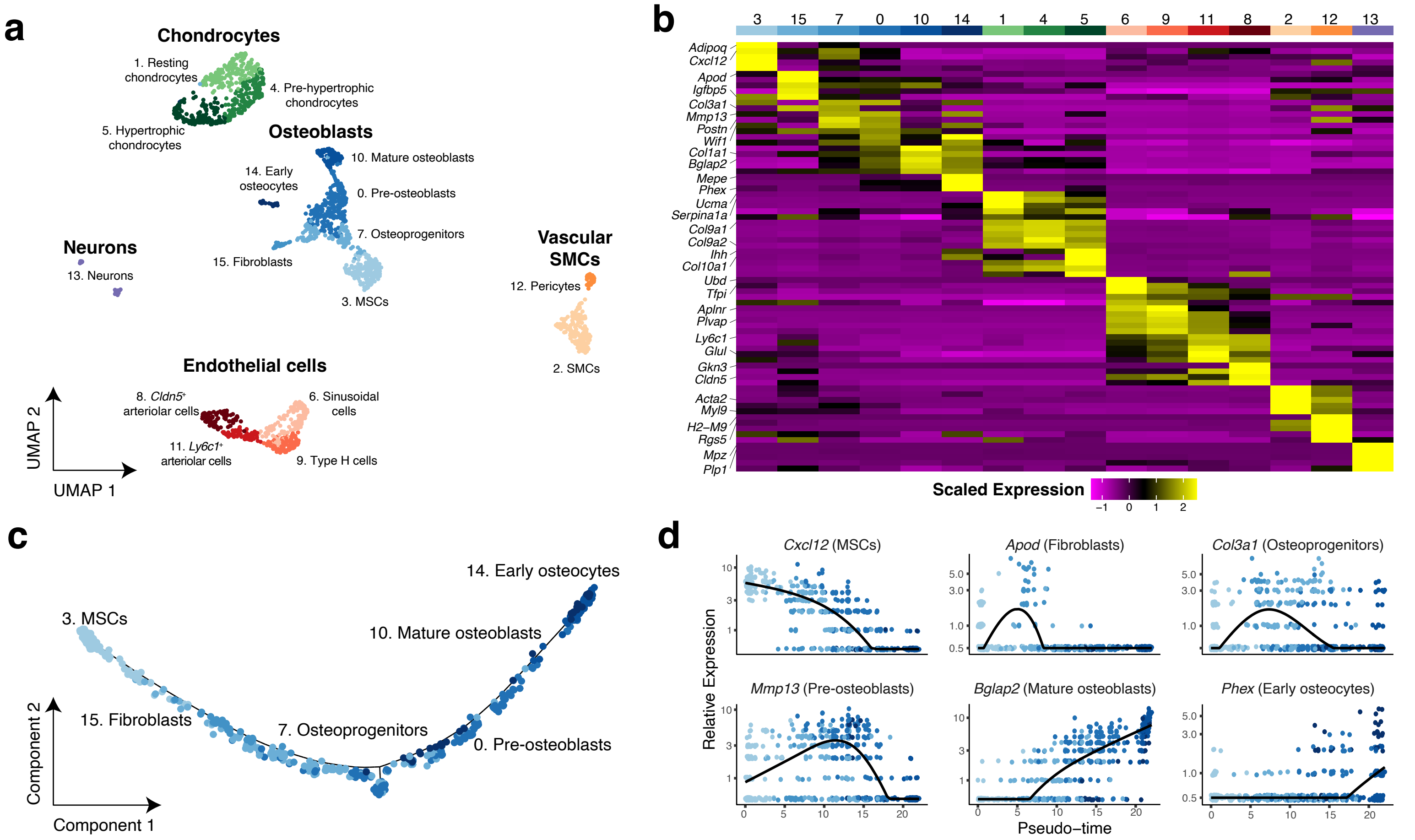


e

| 3. MSCs | | Adjusted *P* value |
|---|---|---|
| GO:0008284 | Positive Regulation Of Cell Population Proliferation | $4.30\times10^{-11}$ |
| GO:0071345 | Cellular Response To Cytokine Stimulus | $9.33\times10^{-11}$ |
| GO:0048522 | Positive Regulation Of Cellular Process | $5.74\times10^{-8}$ |
| GO:0042127 | Regulation Of Cell Population Proliferation | $7.95\times10^{-8}$ |

| 15. Fibroblasts | | Adjusted *P* value |
|---|---|---|
| GO:0030334 | Regulation Of Cell Migration | $1.93\times10^{-5}$ |
| GO:2000146 | Negative Regulation Of Cell Motility | $4.70\times10^{-4}$ |
| GO:0030178 | Negative Regulation Of Wnt Signaling Pathway | $6.00\times10^{-4}$ |
| GO:0010466 | Negative Regulation Of Peptidase Activity | $1.79\times10^{-3}$ |

| 7. Osteoprogenitors | | Adjusted *P* value |
|---|---|---|
| GO:0030198 | Extracellular Matrix Organization | $1.08\times10^{-11}$ |
| GO:0030334 | Regulation Of Cell Migration | $5.98\times10^{-9}$ |
| GO:0006508 | Proteolysis | $1.24\times10^{-5}$ |
| GO:0060828 | Regulation Of Canonical Wnt Signaling Pathway | $2.57\times10^{-4}$ |

| 0. Pre-osteoblasts | | Adjusted *P* value |
|---|---|---|
| GO:0030198 | Extracellular Matrix Organization | $1.88\times10^{-19}$ |
| GO:0043062 | Extracellular Structure Organization | $4.14\times10^{-14}$ |
| GO:0001501 | Skeletal System Development | $1.74\times10^{-8}$ |
| GO:0060348 | Bone Development | $7.53\times10^{-4}$ |

| 10. Mature osteoblasts | | Adjusted *P* value |
|---|---|---|
| GO:0030199 | Collagen Fibril Organization | $1.65\times10^{-9}$ |
| GO:0001501 | Skeletal System Development | $5.40\times10^{-6}$ |
| GO:0006621 | Protein Retention In ER Lumen | $1.05\times10^{-3}$ |
| GO:0060348 | Bone Development | $1.12\times10^{-2}$ |

| 14. Early osteocytes | | Adjusted *P* value |
|---|---|---|
| GO:0060395 | SMAD Protein Signal Transduction | $2.51\times10^{-4}$ |
| GO:0031214 | Biomineral Tissue Development | $3.99\times10^{-4}$ |
| GO:0030278 | Regulation Of Ossification | $8.79\times10^{-4}$ |
| GO:0030501 | Positive Regulation Of Bone Mineralization | $9.22\times10^{-4}$ |

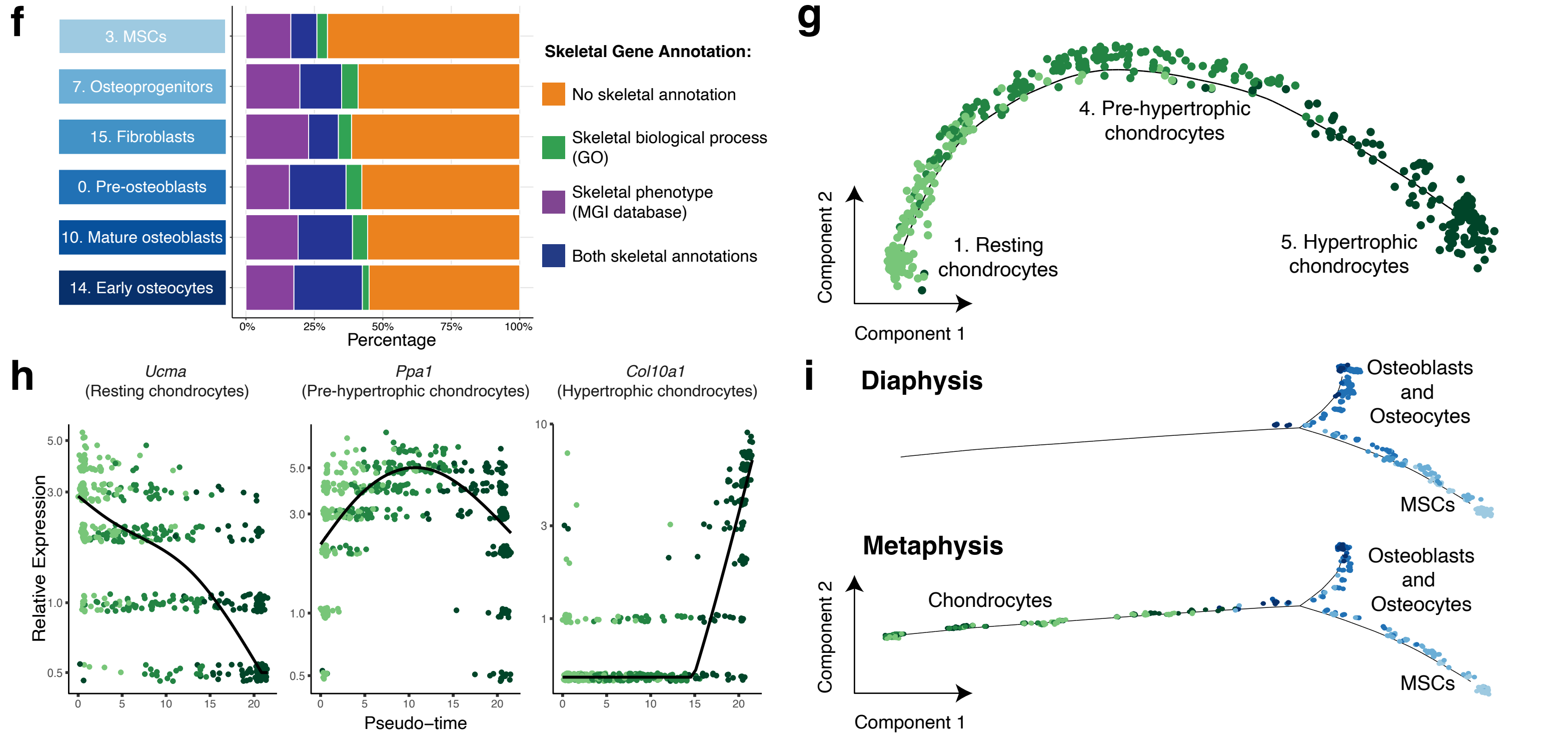
