## Supplementary figures and images for "Multiscale analysis and functional validation of the cellular and genetic determinants of skeletal disease"

### Extended Data Figure 4

#### Extended Data Fig. 4. Analysis of transcription factor activity in non-haematopoietic cells

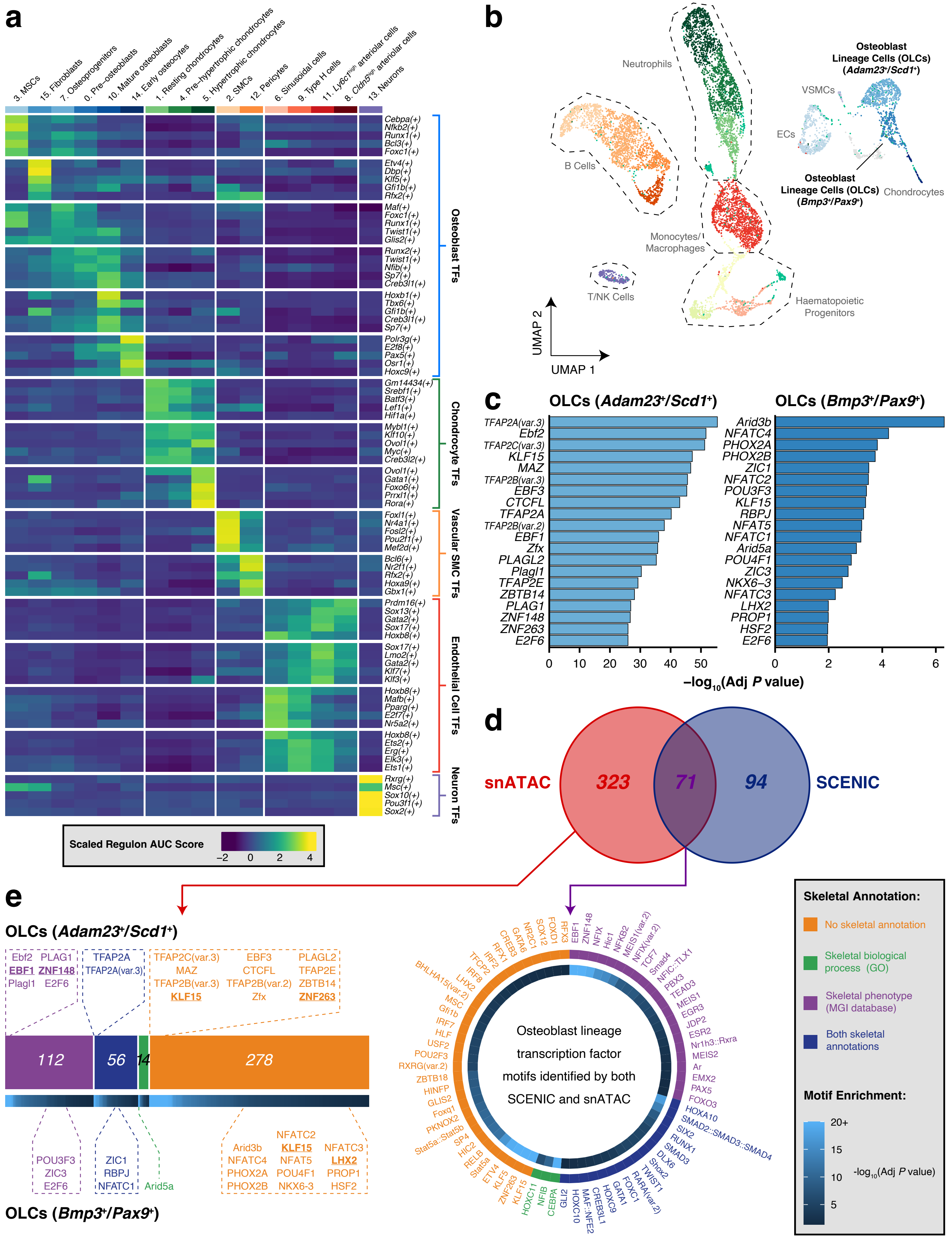

### Extended Data Figure 6

##### Extended Data Fig. 6. Genome-wide association study of eBMD in the UK Biobank study

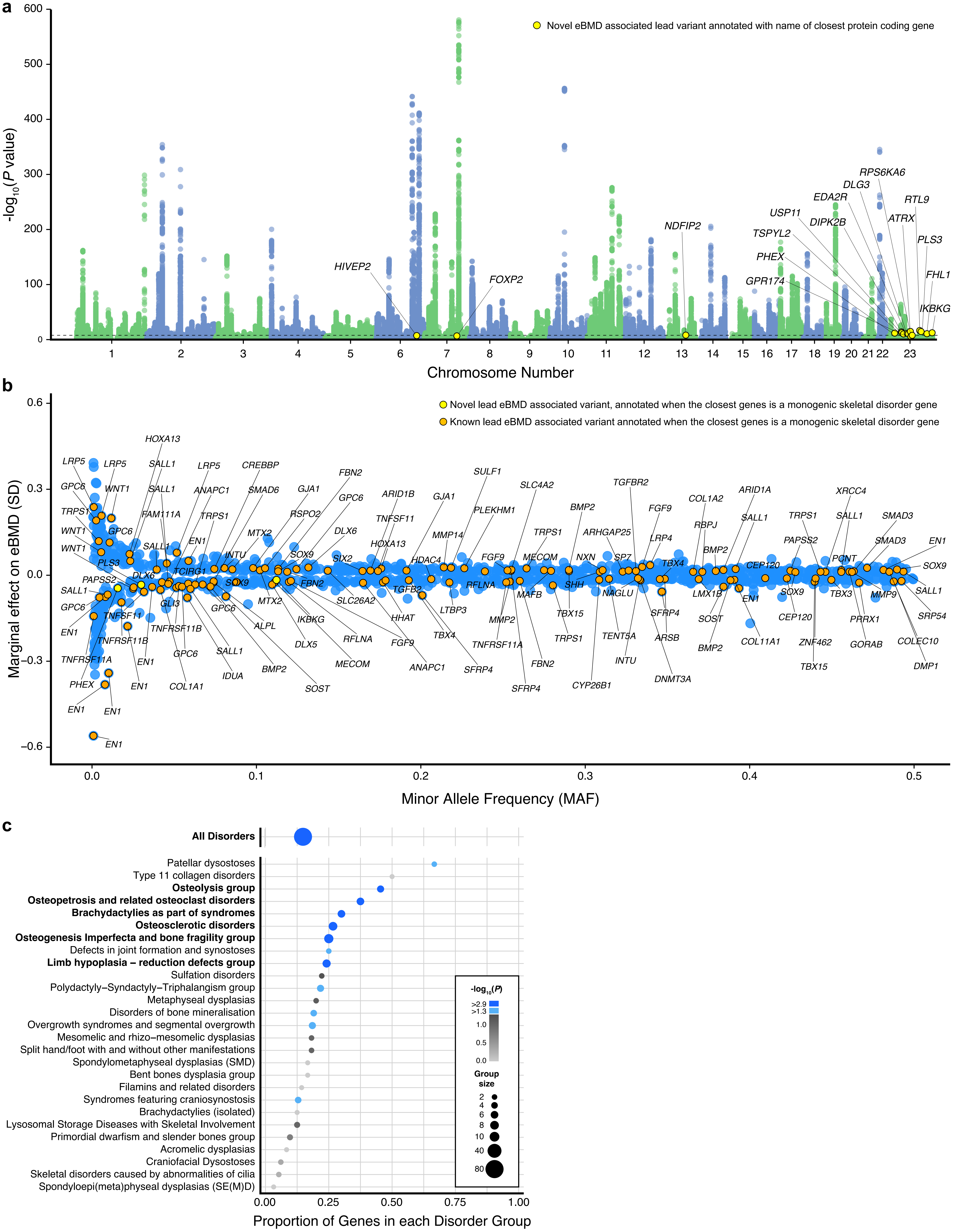

### Extended Data Figure 9

#### Extended Data Fig. 9. Cellular mechanism of skeletal phenotype in *Pls3* deficient mice

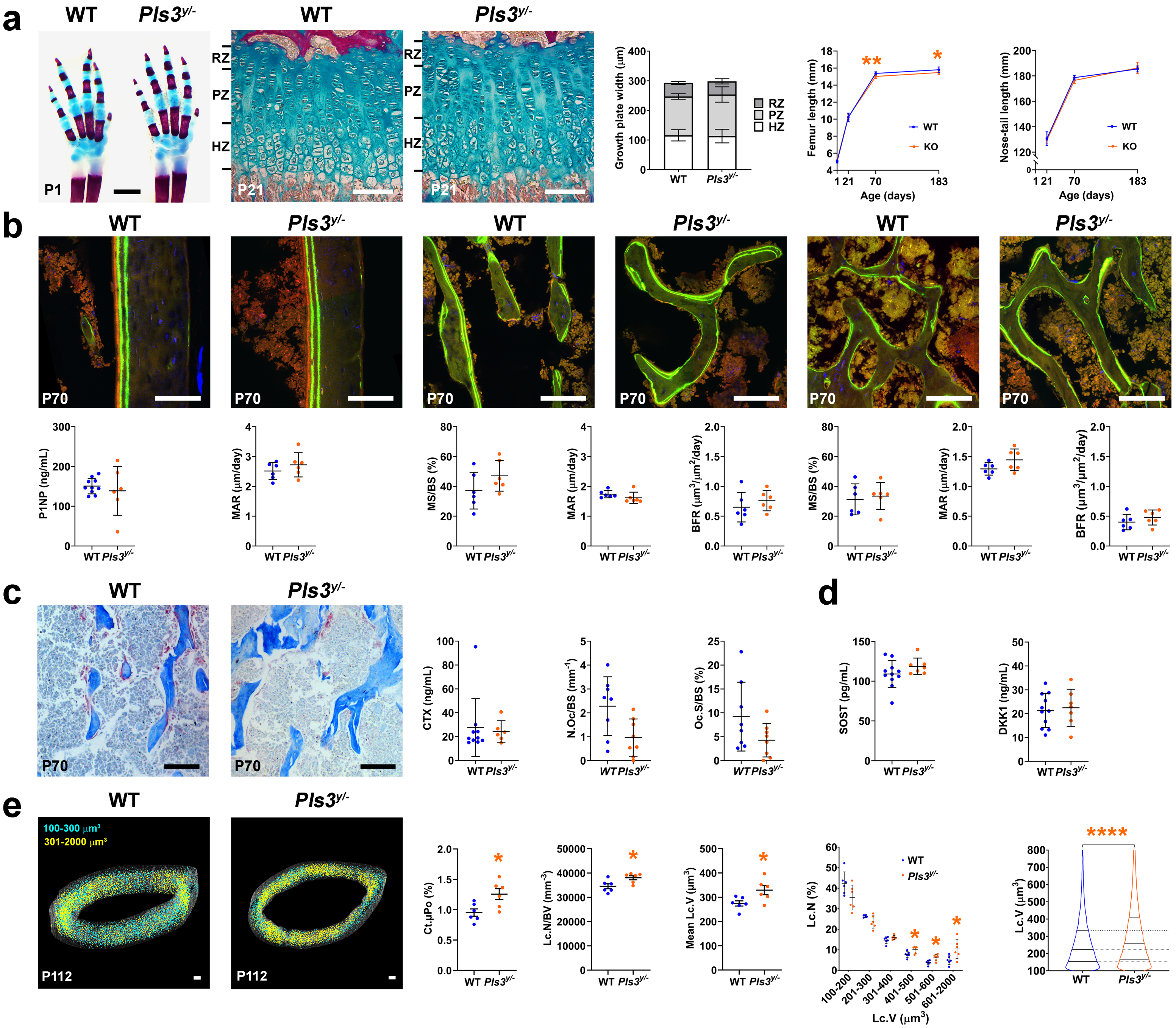

### Extended Data Figure 10

#### Extended Data Fig. 10. Spatial transcriptomics of human bone samples

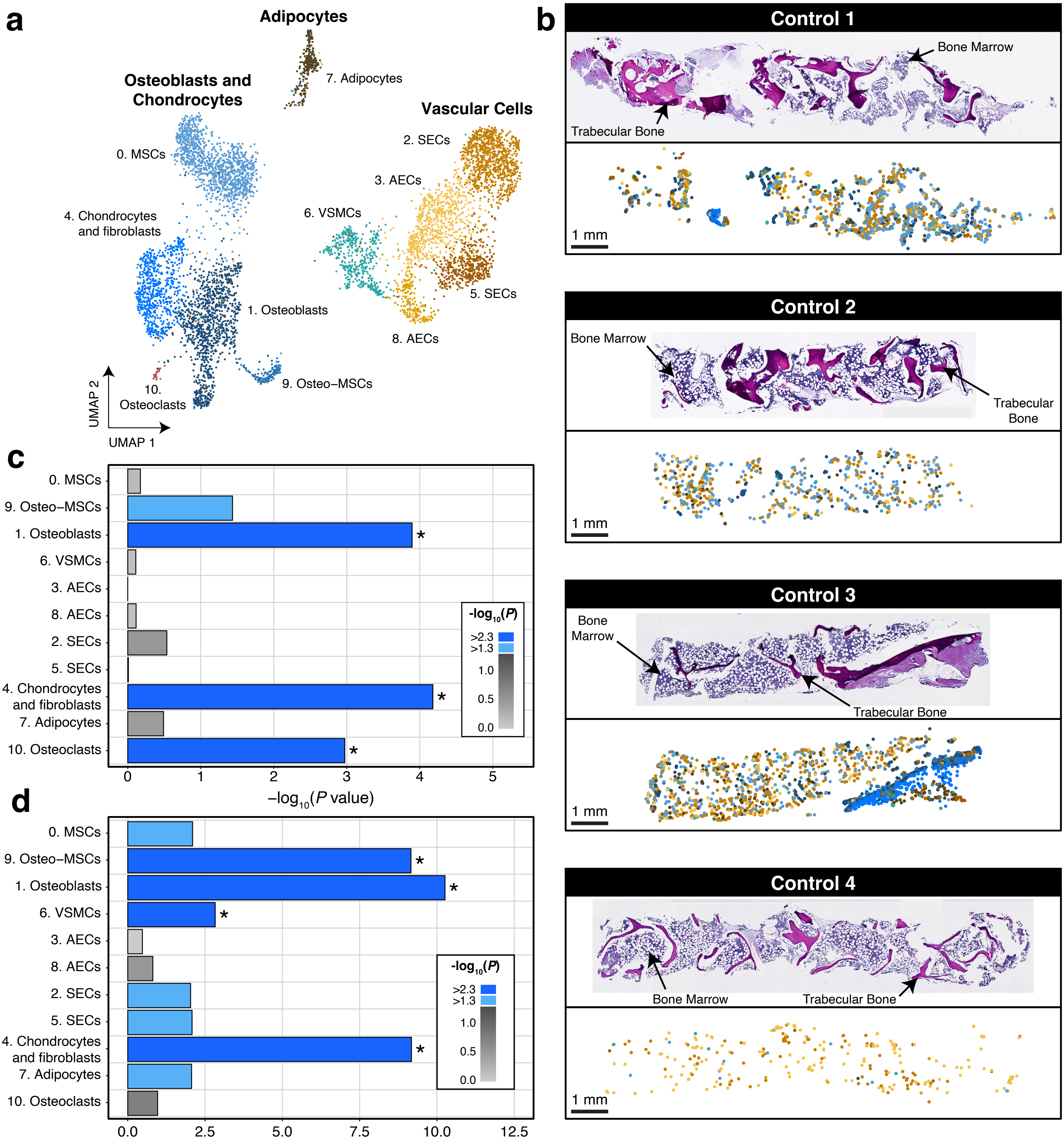

### Supplementary Figure 5

#### Supplementary Fig. 5. Expression of exemplar genes in tissues outside of the skeleton.

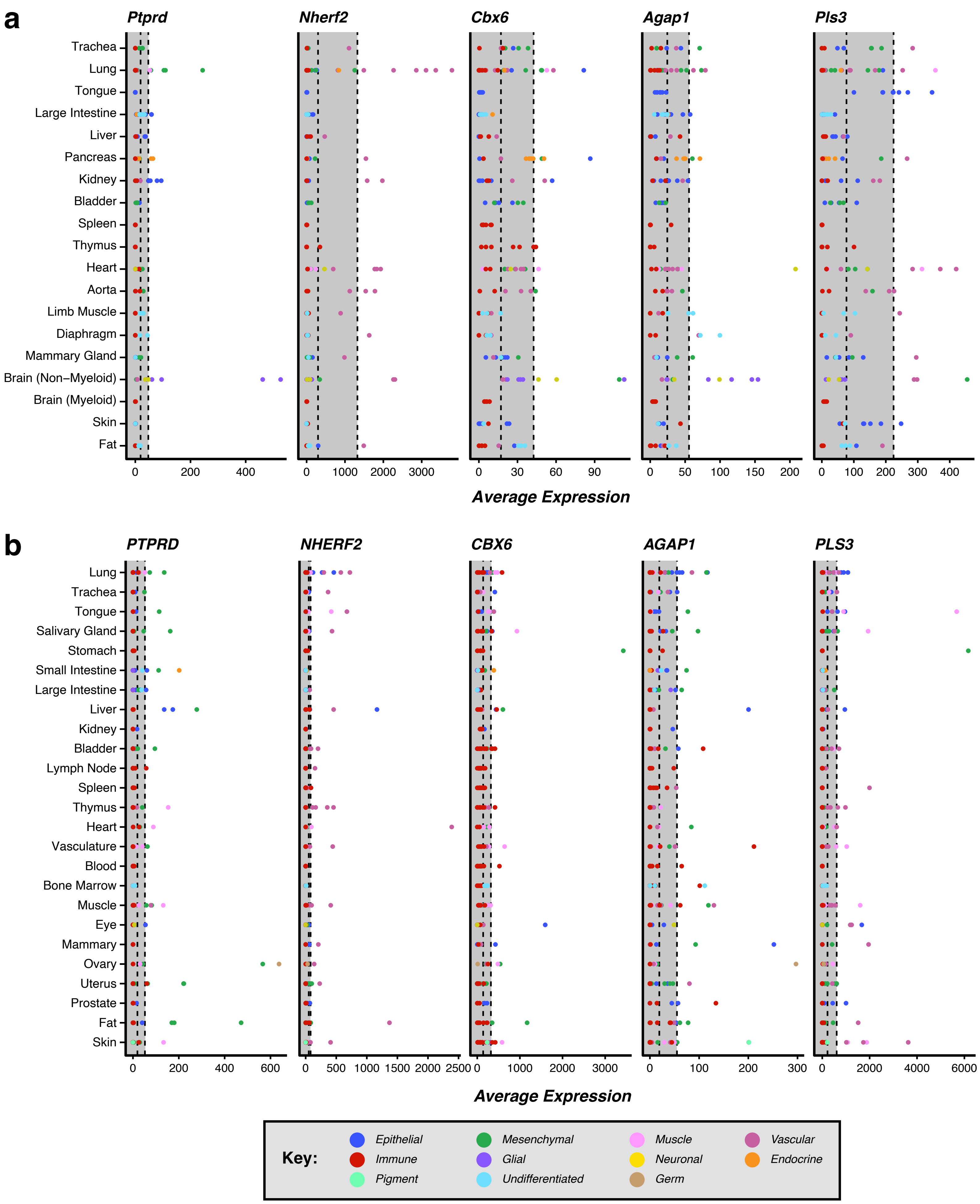

### Supplementary Figure 6

Supplementary Fig. 6. Skeletal phenotyping of younger male *Pls3*$^{y/-}$ mice

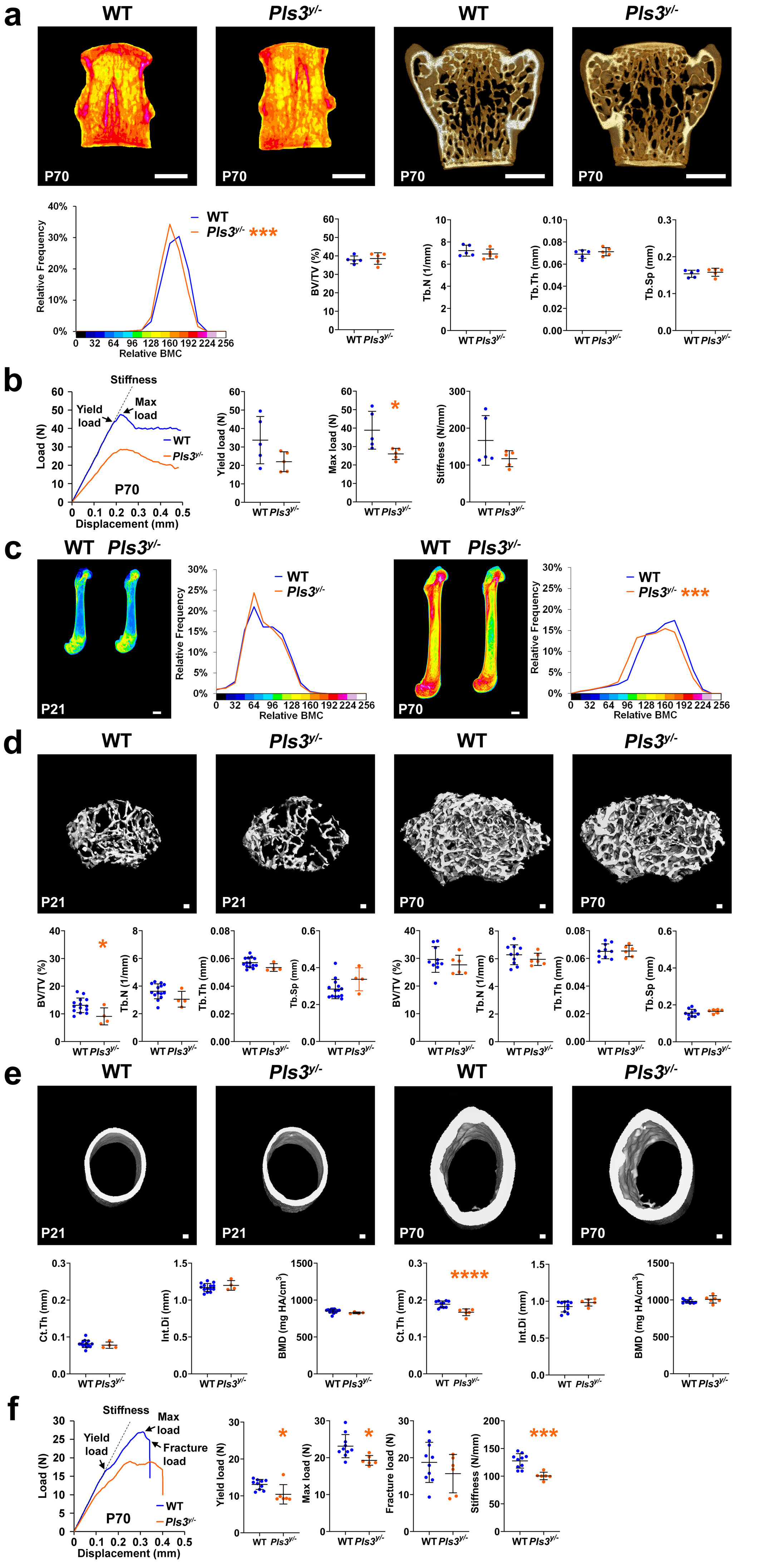

### Supplementary Figure 7

##### Supplementary Fig. 7. Skeletal phenotyping of female $Pls3^{y/-}$ mice

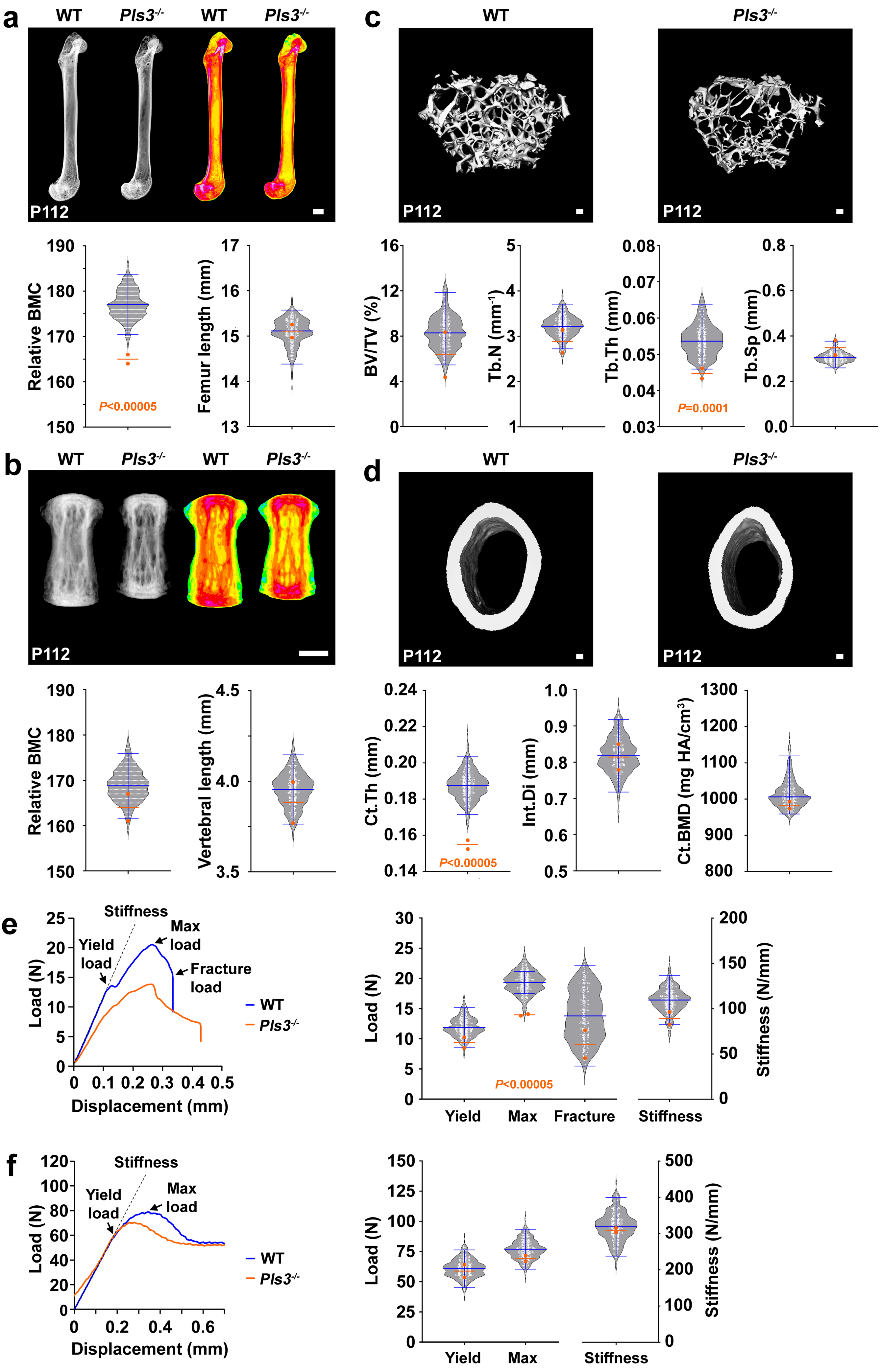

### Supplementary Figure 8

##### Supplementary Fig. 8. Skeletal vascular phenotyping of male *Pls3*$^{y/-}$ mice

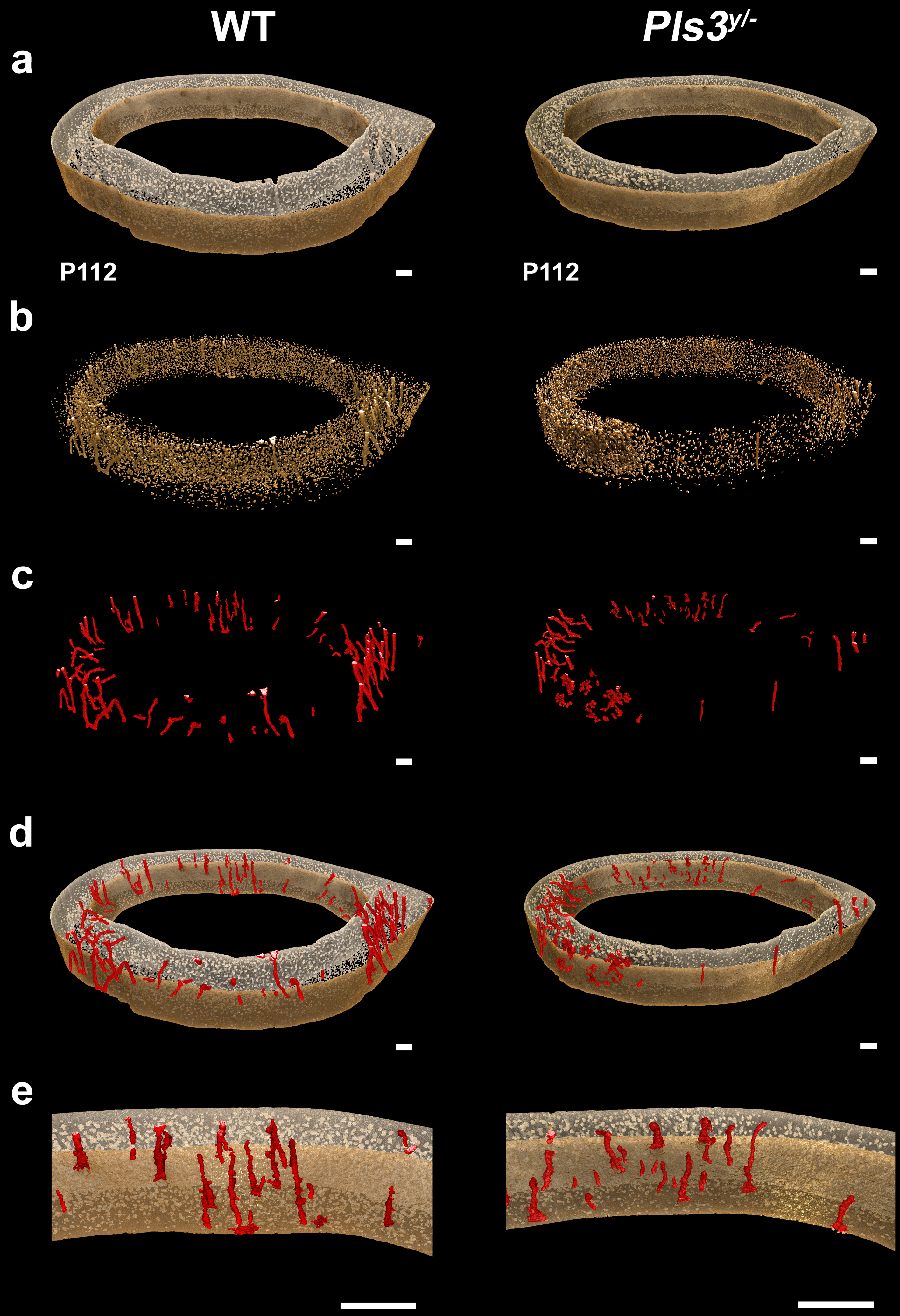

### Supplementary Note 1

#### Supplementary Note 1. Annotation of cell types in mouse scRNA-seq data

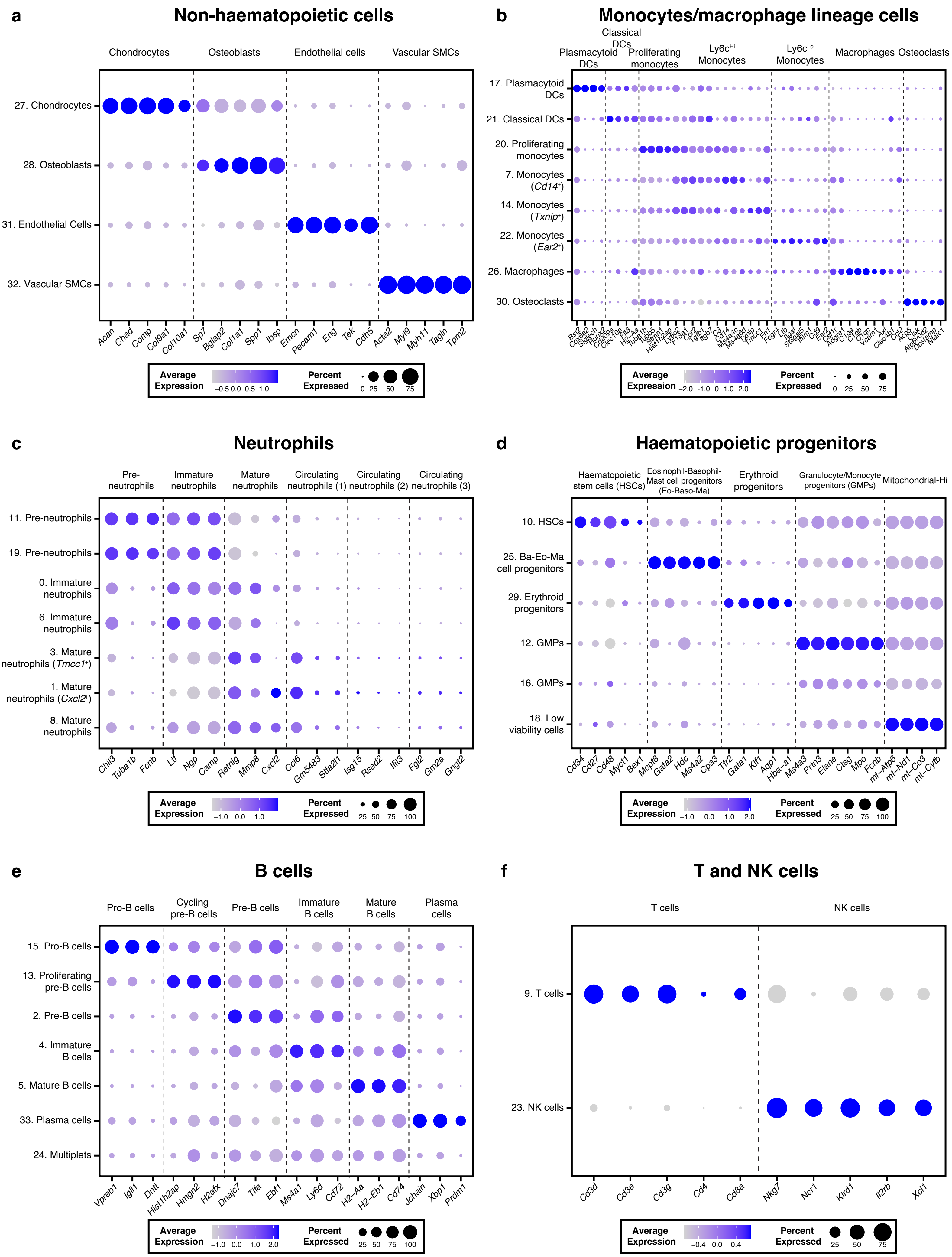
