## Extended Data Figure 5 for "Multiscale analysis and functional validation of the cellular and genetic determinants of skeletal disease"

### Extended Data Fig. 5. Gene programs of non-haematopoietic cell sub-clusters are enriched with monogenic skeletal disorder genes

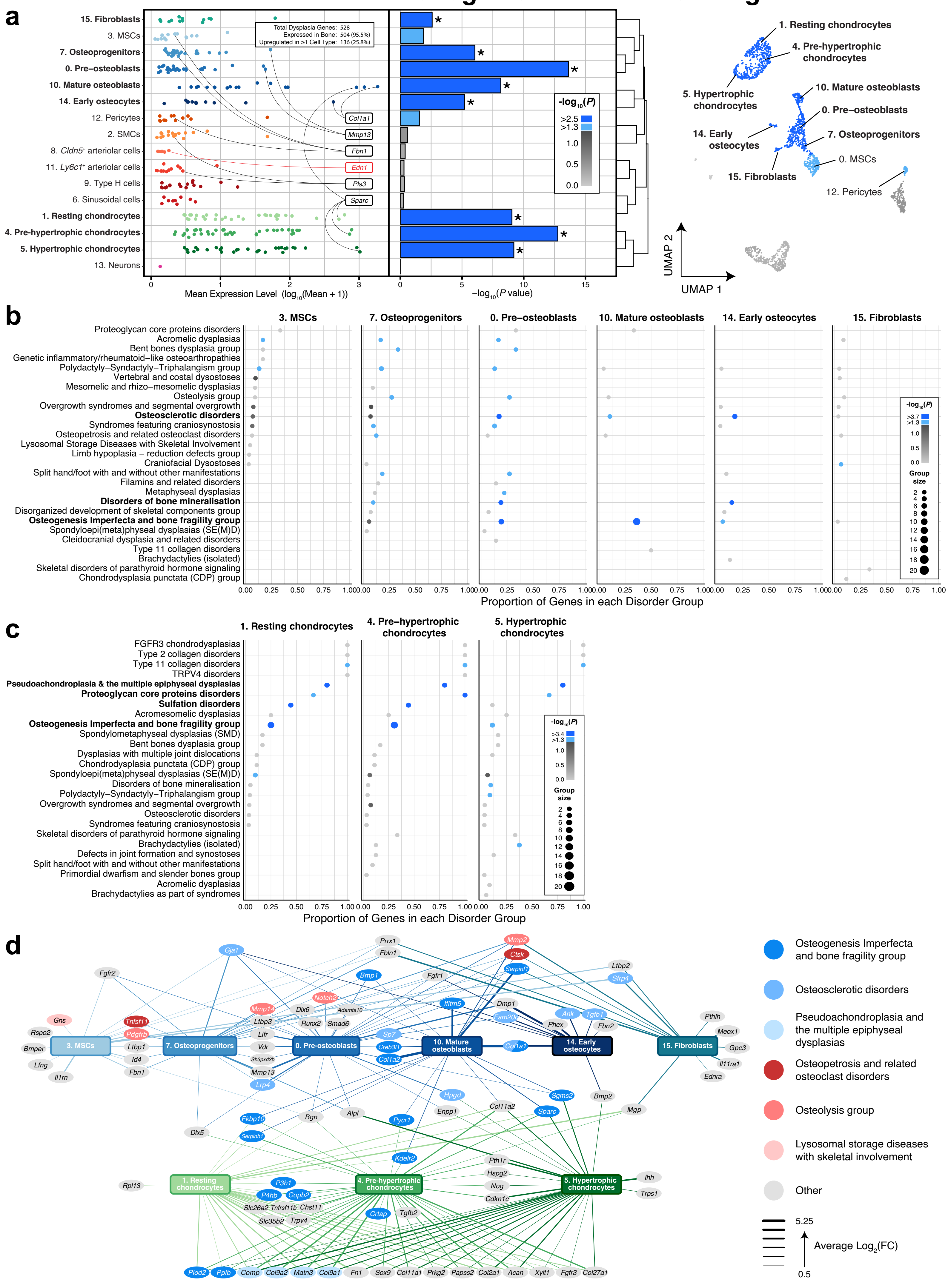
