## Extended Data Figure 7 for "Multiscale analysis and functional validation of the cellular and genetic determinants of skeletal disease"

### Extended Data Fig. 7. Gene programs of non-haematopoietic cell sub-clusters are enriched with eBMD-associated genes

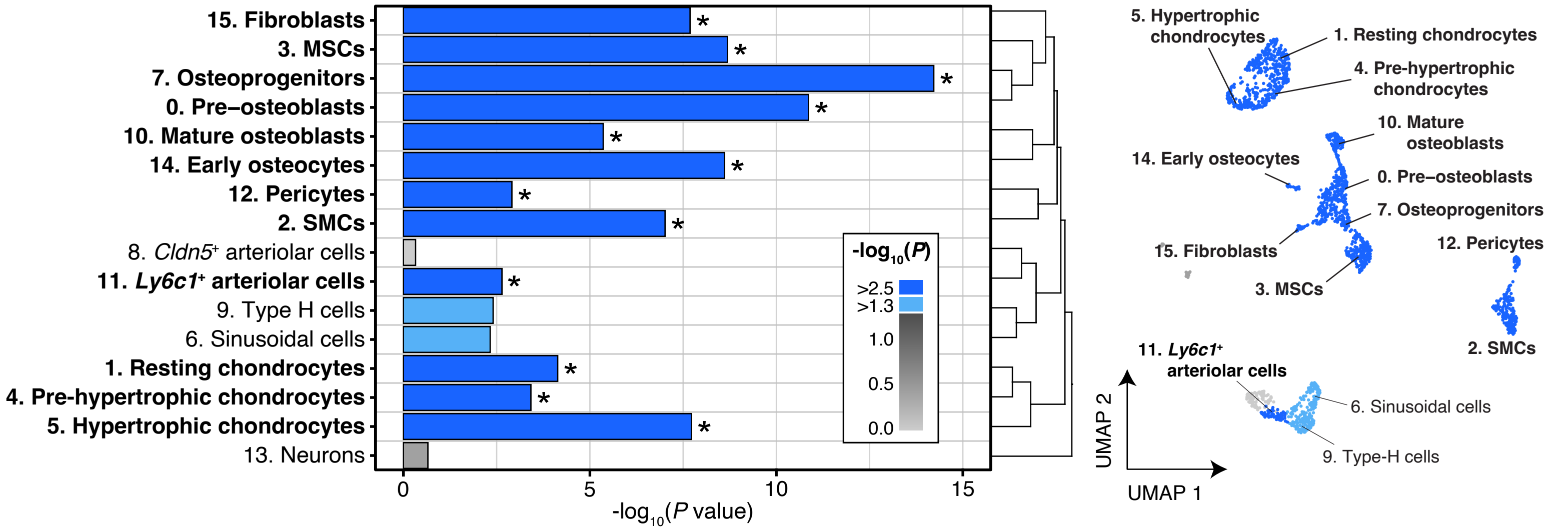
