## Extended Data Figure 8 for "Multiscale analysis and functional validation of the cellular and genetic determinants of skeletal disease"

### Extended Data Fig. 8. Gene programs of non-haematopoietic cells and osteoclasts are enriched with genes that cause abnormal bone structure when mutated in mice

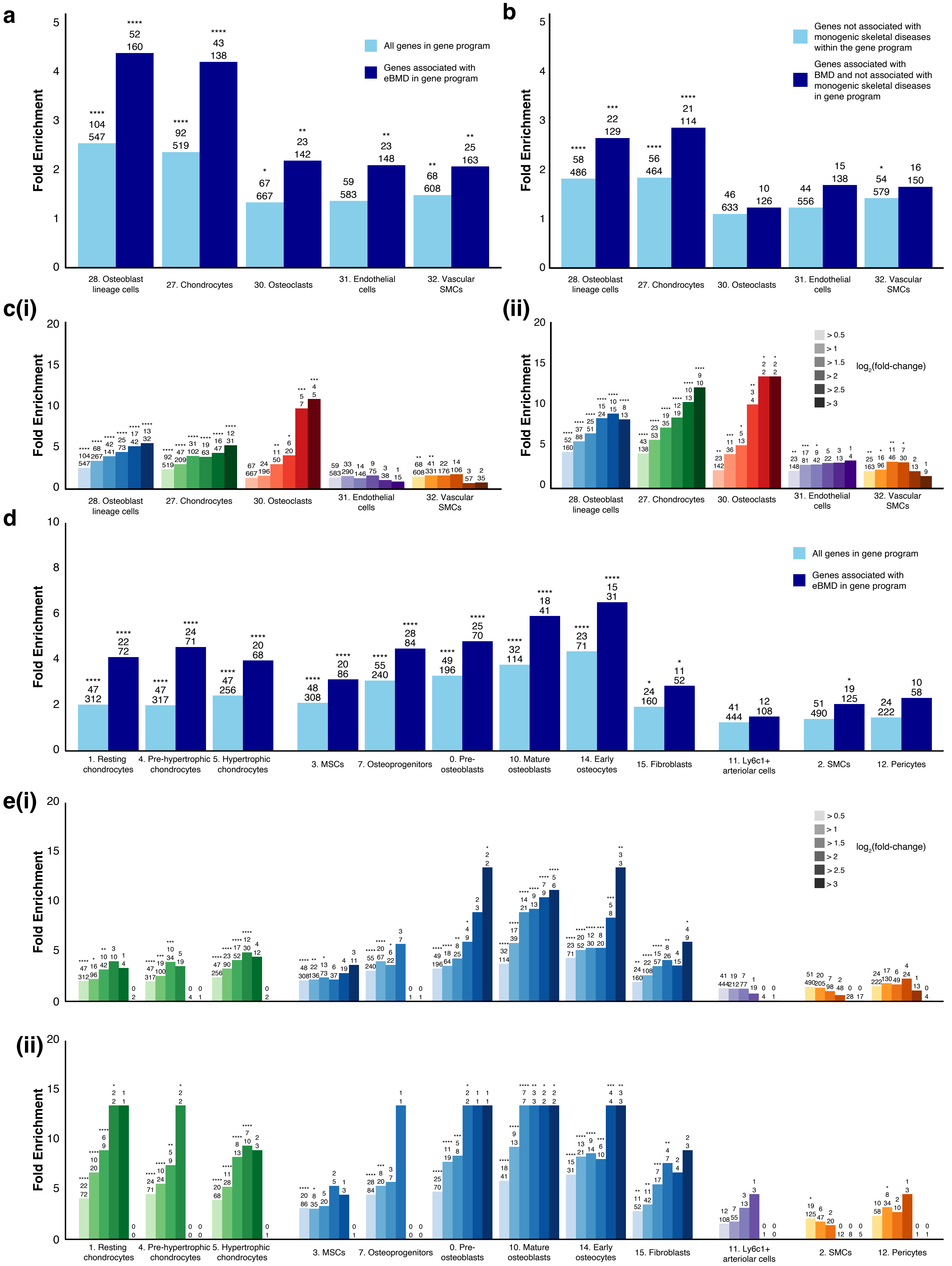
