## Supplementary Figure 1 for "Multiscale analysis and functional validation of the cellular and genetic determinants of skeletal disease"

### Supplementary Fig. 1. Validation of cell populations enriched in the endosteal compartment of mouse bone.

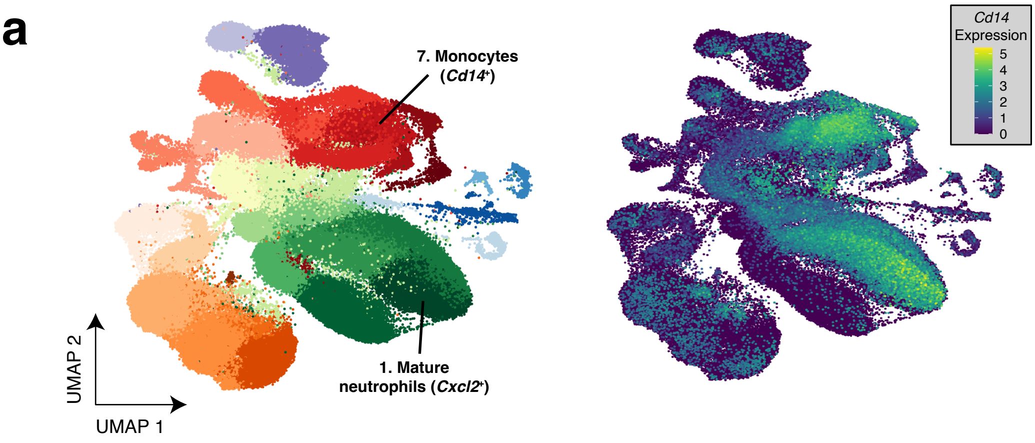


b

| | Gene | Average $Log_2FC$ | Adjusted p Value |
|---|---|---|---|
| 7. Monocytes (*Cd14*+) | *Ctss* | 2.76 | 0 |
| | *Ms4a6c* | 2.75 | 0 |
| | *Ms4a4c* | 2.64 | 0 |
| | *S100a4* | 2.63 | 0 |
| | *Ccr2* | 2.53 | 0 |
| | *Ccl9* | 2.53 | 0 |
| | *Apoe* | 2.40 | 0 |
| | *Ctsc* | 2.30 | 0 |
| | *Cd14* | 2.28 | 0 |
| 1. Mature neutrophils (*Cxcl2*+) | **Gene** | **Average $Log_2FC$** | **Adjusted p Value** |
| | *Il1b* | 3.39 | 0 |
| | *Dusp1* | 2.71 | 0 |
| | *Hcar2* | 2.58 | 0 |
| | *Ccl6* | 2.40 | 0 |
| | *Cebpb* | 2.40 | 0 |
| | *Egr1* | 2.28 | 0 |
| | *Slc7a11* | 2.27 | 0 |
| | ... | ... | ... |
| | *Cd14* | 1.18 | 0 |

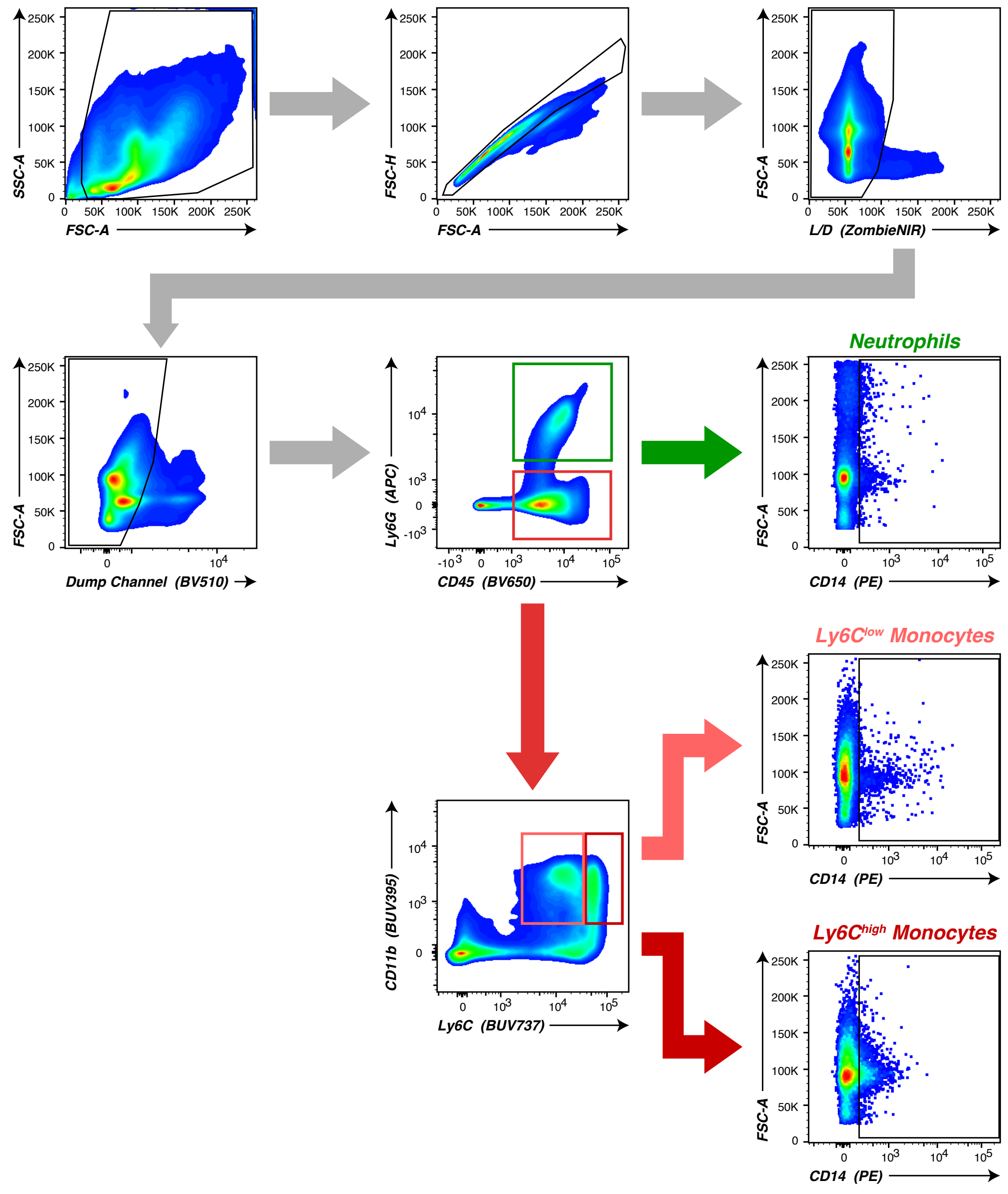


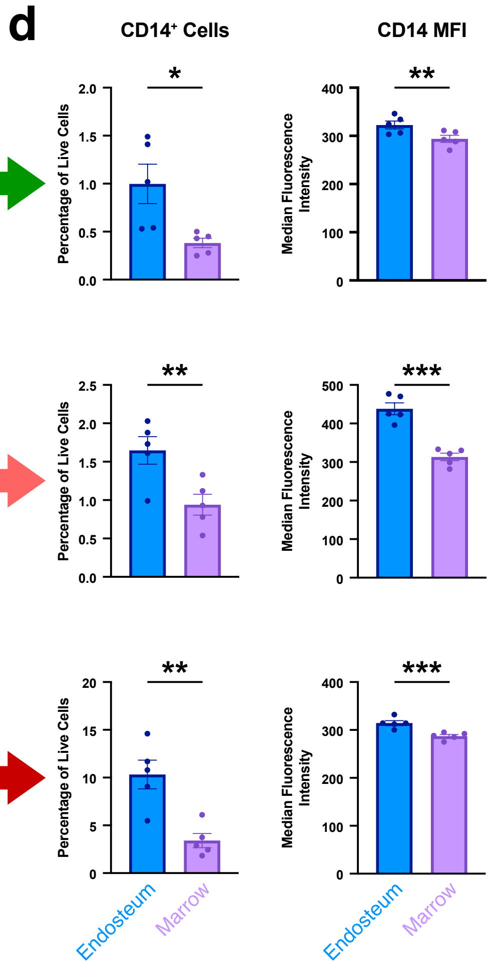
