## Supplementary Figure 2 for "Multiscale analysis and functional validation of the cellular and genetic determinants of skeletal disease"

### Supplementary Fig. 2. Distribution of non-haematopoietic cell sub-clusters in diaphysis and metaphysis

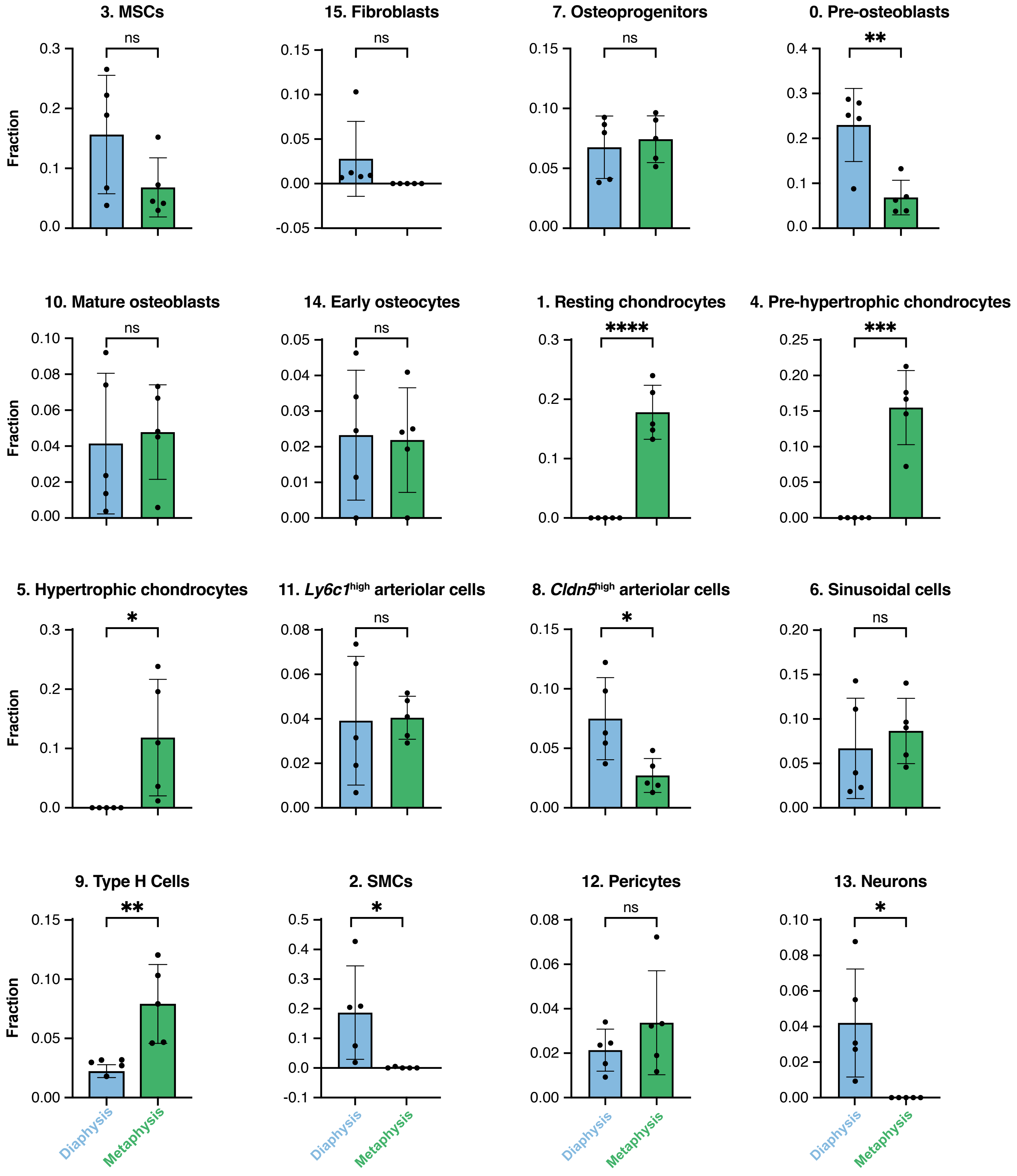
