## Supplementary Figure 3 for "Multiscale analysis and functional validation of the cellular and genetic determinants of skeletal disease"

### Supplementary Fig. 3. Genes associated with eBMD, but not pulse rate, are enriched with monogenic skeletal disorder genes

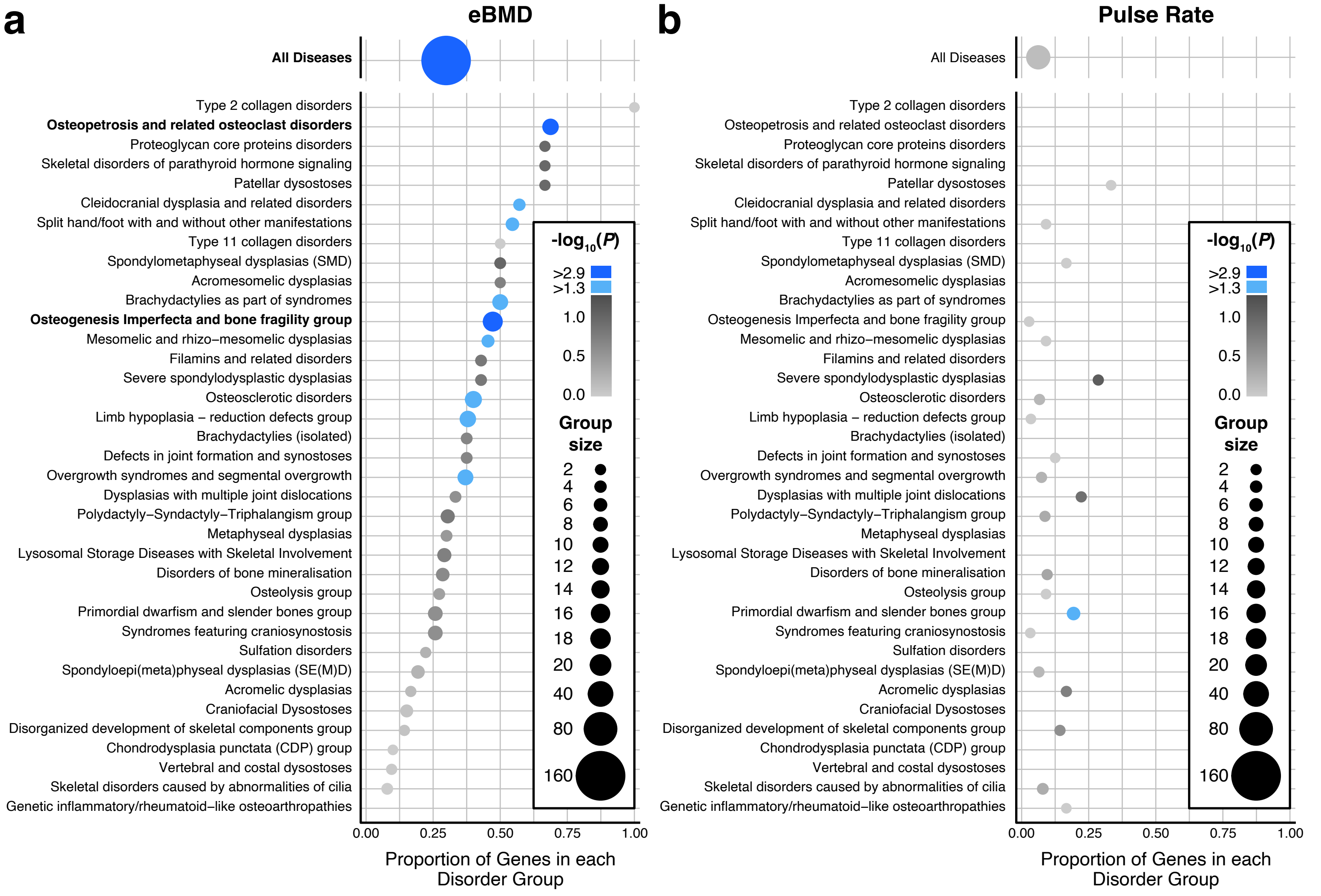
