## Supplementary Figure 4 for "Multiscale analysis and functional validation of the cellular and genetic determinants of skeletal disease"

### Supplementary Fig. 4. Conditional GSA analysis to determine whether enrichment was confounded by shared sets of genes

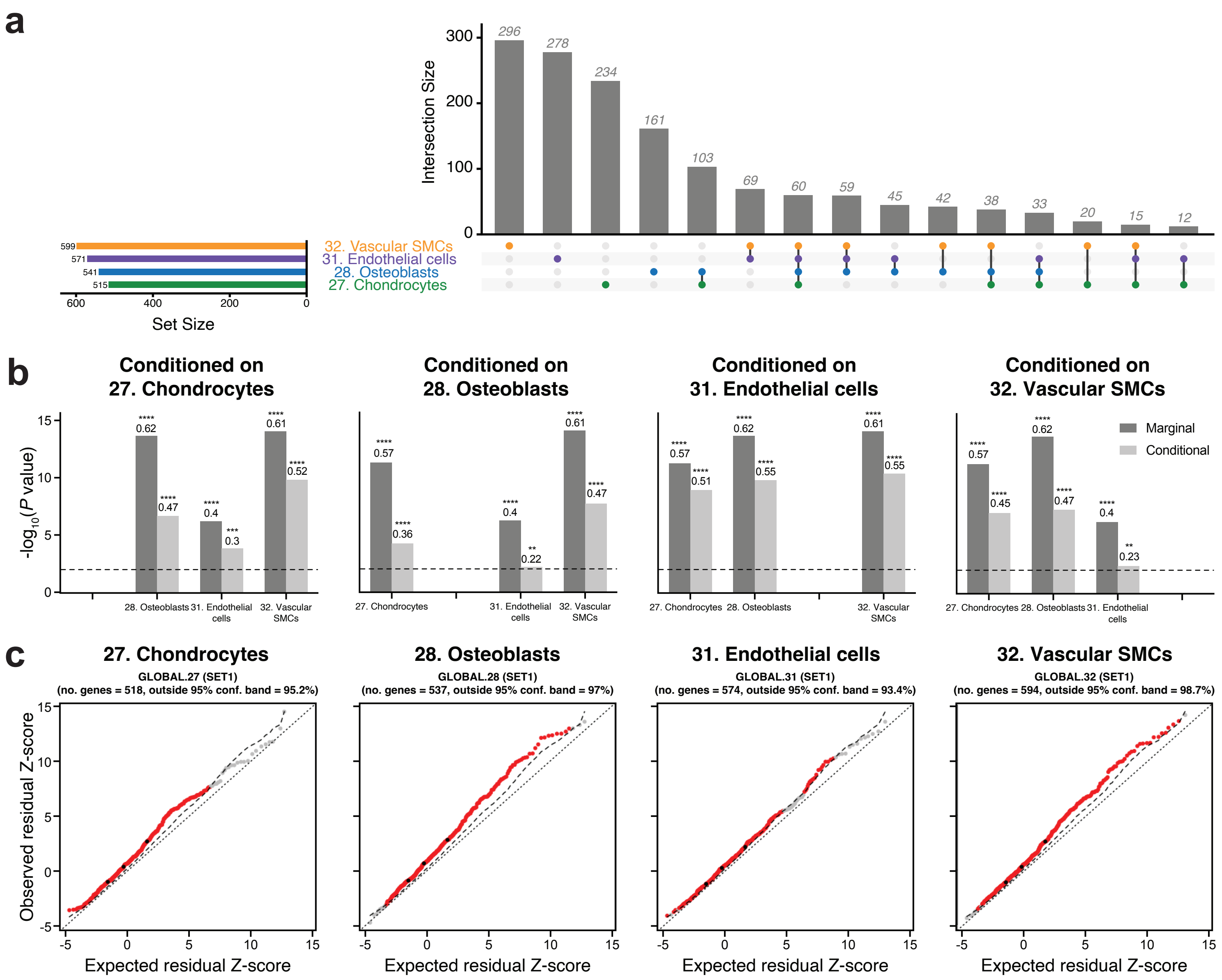
