## Supplementary Table 1 for "Multiscale analysis and functional validation of the cellular and genetic determinants of skeletal disease"

### Supplementary Table 1 - Glossary of Terms and Definitions:

This glossary defines key terms used in this manuscript. Detailed technical information are provided in the Materials and Methods.

| Term | Definition |
|---|---|
| Endosteal bone compartment | The layer of cells located at the interface between bone and the bone marrow. Includes cells that are adjacent to bone and those up to 10 cells away from the bone surface. |
| Bone marrow compartment | The cells in the bone marrow space that are not adjacent to bone and are not part of the endosteal bone compartment. |
| Cluster | A group of cells that share similar patterns of gene expression. Cluster membership is based on the magnitude of expression and restrictedness of ~3000 highly variable genes |
| Restrictedness | A term used to describe whether a gene is expressed in a single cluster or found in more than one cluster. |
| Gene program | The set of all genes that are differentially upregulated in a given cluster relative to all other cells in the dataset. |
| Pathogenic variant | A genetic variant that changes the function of a gene in a way that causes a monogenetic disorder. |
| Monogenetic | A trait or disorder resulting from pathogenic variants that occur within a single gene. |
| Causative gene | A gene that causes a rare monogenetic disorder when its function is altered by a pathogenic variant. |
| Nosology | A classification of individual monogenetic disorders caused by pathogenic variants in single genes. |
| Causal variant | A genetic variant that changes the function of a gene in a way that alters a polygenetic trait and/or susceptibility to disease. |
| Polygenetic | A trait or disorder that result from the contribution of many independently acting or interacting causal genetic variants. |
| Effector gene | A gene that causes a change in a polygenetic trait and/or susceptibility to disease when its function is altered by causal variants. |
| Abnormal bone structure from MGI | Anomaly in the composite material or the layered arrangement of the bony endoskeleton of the body |
| Gene set enrichment analysis | A family of statistical methods that can be used to identify whether a set of genes are over-represented among a gene program. |
| High impact variant | A genetic variant that is predicted to have a disruptive impact on the protein, causing protein |

| | |
|---|---|
| | truncation, loss of function or triggering nonsense mediated decay. |
| Moderate impact variant | A genetic variant that is non-disruptive, but likely to change protein expression level and/or function. |
| Low impact variant | A genetic variant predicted to be mostly non-deleterious or unlikely to change protein expression level and/or function. |
| Locus | A region in the genome that contains genetic variants that are associated with variation of a polygenetic trait and/or susceptibility to disease. |
| Lead variant | The genetic variant that is most strongly associated at a locus after accounting for all other associated genetic variants. |
| Structural phenotypes | Significant difference in bone structural parameters measured by digital X-ray microradiography or micro-computerised tomography. |
| Functional phenotypes | Significant differences in mechanical strength parameters measured by biomechanical testing |
| Structural and functional phenotypes | Significant differences in at least one structural and one functional parameter |
| Bone quality phenotypes | A term that encompasses various structural parameters of bone that affect fragility, including bone microarchitecture, mineralization, and material properties [Grynpas, M. D. (2003). The role of bone quality on bone loss and bone fragility. In Bone loss and osteoporosis: An anthropological perspective (pp. 33-44). Boston, MA: Springer US.]. In the OBCD study, it refers to lines with outlier functional phenotypes that did not correlate with bone mineral content |
| Mahalanobis phenotypes | Lines that were significant outliers due to smaller differences in multiple skeletal parameters. |
