## Supplementary Note 3 for "Multiscale analysis and functional validation of the cellular and genetic determinants of skeletal disease"

**Supplementary Note 3. Overview of the FACS strategy to collect live non-erythroid cells from mouse and human bones**

**a** **Mouse**

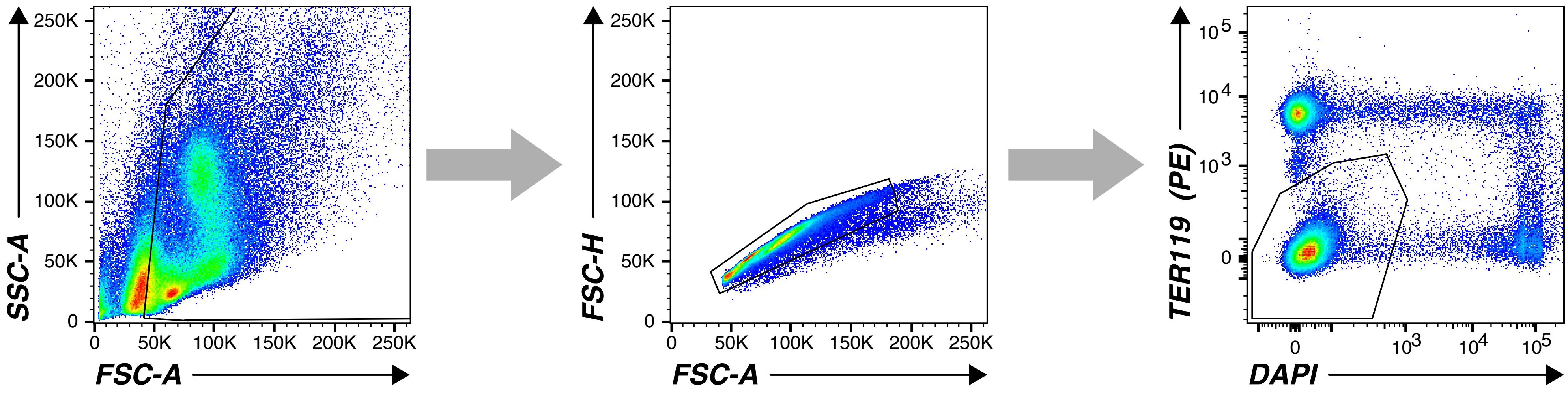


**b** **Human**

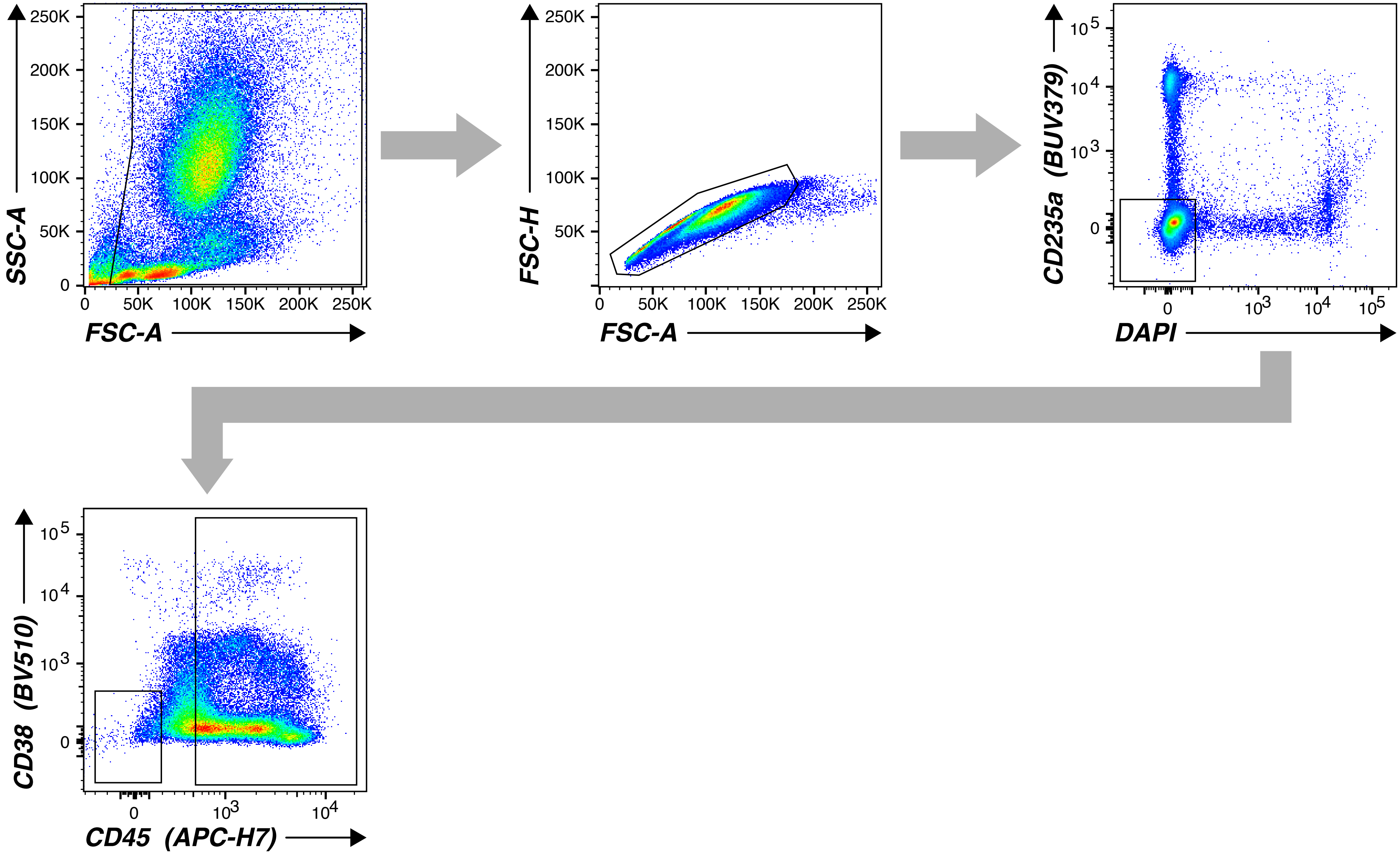
